# PIWI-piRNA pathway-mediated transposable element repression in *Hydra* somatic stem cells

**DOI:** 10.1101/731695

**Authors:** Bryan B. Teefy, Stefan Siebert, Jack F. Cazet, Haifan Lin, Celina E. Juliano

## Abstract

Transposable elements (TEs) can damage genomes, thus organisms employ a variety of mechanisms to repress TE expression. However, these mechanisms often fail over time leading to de-repression of TEs in aging tissues. The PIWI-piRNA pathway is a small RNA pathway that represses TE expression in the germline of animals. Here we explore the function of the pathway in the epithelial stem cells of *Hydra*, a long-lived freshwater cnidarian. *Hydra* have three stem cell populations; endodermal and ectodermal epithelial stem cells are strictly somatic, whereas the interstitial stem cells retain germline competence. In our previous study, we found that the PIWI proteins are expressed in all three *Hydra* stem cell types. In this study, we focus on the ectodermal and endodermal epithelial stem cells to understand the somatic function of the pathway. We isolated piRNAs from *Hydra* that lack the interstitial lineage and found that these somatic piRNAs map predominantly to TE transcripts and display the conserved sequence signatures typical of germline piRNAs. Three lines of evidence suggest that the PIWI-piRNA pathway represses TEs in *Hydra* epithelial stem cells. First, epithelial knockdown of the *Hydra* PIWI protein *hywi* resulted in upregulation of TE expression. Second, degradome sequencing revealed evidence of PIWI-mediated cleavage of TE RNAs in epithelial cells using the ping-pong mechanism. Finally, we demonstrated a direct association between Hywi protein and TE transcripts in epithelial cells using RNA immunoprecipitation. Interestingly, we found that RNAi knockdown of *hywi* leads to an upregulation of genes involved in innate immunity, which may be in response to TE upregulation; this is consistent with recent studies on TE expression in mammalian cells. Altogether, this study suggests a function for the PIWI-piRNA pathway in maintaining the long-lived somatic cell lineages of *Hydra* and may point to a broader role for this pathway in protecting somatic tissue from TE-induced damage.

## Introduction

A prominent feature of somatic aging is the breakdown of heterochromatic domains and expression of transposable elements (TEs), which are typically repressed (López-Otín et al. 2013; Cecco et al. 2013; Benayoun et al. 2018; Jones et al. 2016; Li et al. 2013; Patterson et al. 2015; Hendrickson et al. 2018; Sousa-Victor et al. 2017). Although TE expression is likely a secondary effect of heterochromatin disruption, mounting evidence suggests that suppressing TE mobilization can extend lifespan (Wood et al. 2016; Jones et al. 2016; Wang et al. 2011; Cecco et al. 2019; Simon et al. 2019). Therefore, it is of interest to understand how long-lived organisms, such as *Hydra vulgaris*, repress TE expression in somatic cells. A longevity study demonstrated that mortality in *Hydra* does not increase with age, suggesting a lack of senescence (Schaible et al. 2015), but the mechanisms by which *Hydra* escapes aging are not understood. The best-studied function of the PIWI-piRNA pathway is to repress TE expression in the germline (Iwasaki et al. 2015). However, we found that this pathway is active in the strictly somatic epithelial stem cells of *Hydra* (Juliano et al. 2014). This suggested the possibility that the activity of the PIWI-piRNA pathway in *Hydra* somatic stem cells contributes to longevity through repression of TE expression. This may also be the case for other long-lived animals such as planarians (Sturm et al. 2017; Petralia et al. 2014; Shibata et al. 2016; Palakodeti et al. 2008; Reddien et al. 2005). However, the direct RNA targets of the pathway in the somatic cells of long-lived animals are largely unexplored.

*Hydra* is composed of three cell lineages: ectodermal epithelial, endodermal epithelial, and interstitial; each lineage is supported by its own population of stem cells (**Fig. 1A**). The basic *Hydra* body plan consists of two epithelial monolayers (**Fig. 1A**). All body column epithelial cells are mitotically active, leading to displacement of cells towards the extremities. At the aboral and oral ends, epithelial cells stop dividing and terminally differentiate to build the tentacles, hypostome, and basal disk. The mitotic ectodermal and endodermal body column cells simultaneously function as epitheliomuscular cells and as epithelial stem cells. No fully undifferentiated stem cells exist to renew the epithelium, but rather all body column epithelial cells are able to self-renew and terminally differentiate into the epithelial cell types found at the extremities. Cells of the interstitial lineage include neurons, gland cells, nematocytes, and germ cells (**Fig. 1A**). This lineage is supported by an undifferentiated multipotent interstitial stem cell (ISC) population that continually produces neurons, gland cells, and nematocytes in a homeostatic adult animal. ISCs are also capable of producing germline stem cells (GSCs) when GSCs are experimentally depleted (Bosch and David 1987; Nishimiya-Fujisawa and Kobayashi 2012). Therefore, while ISCs have germline potential, ectodermal and endodermal epithelial stem cells (ESCs) are strictly somatic.

**Figure 1.**
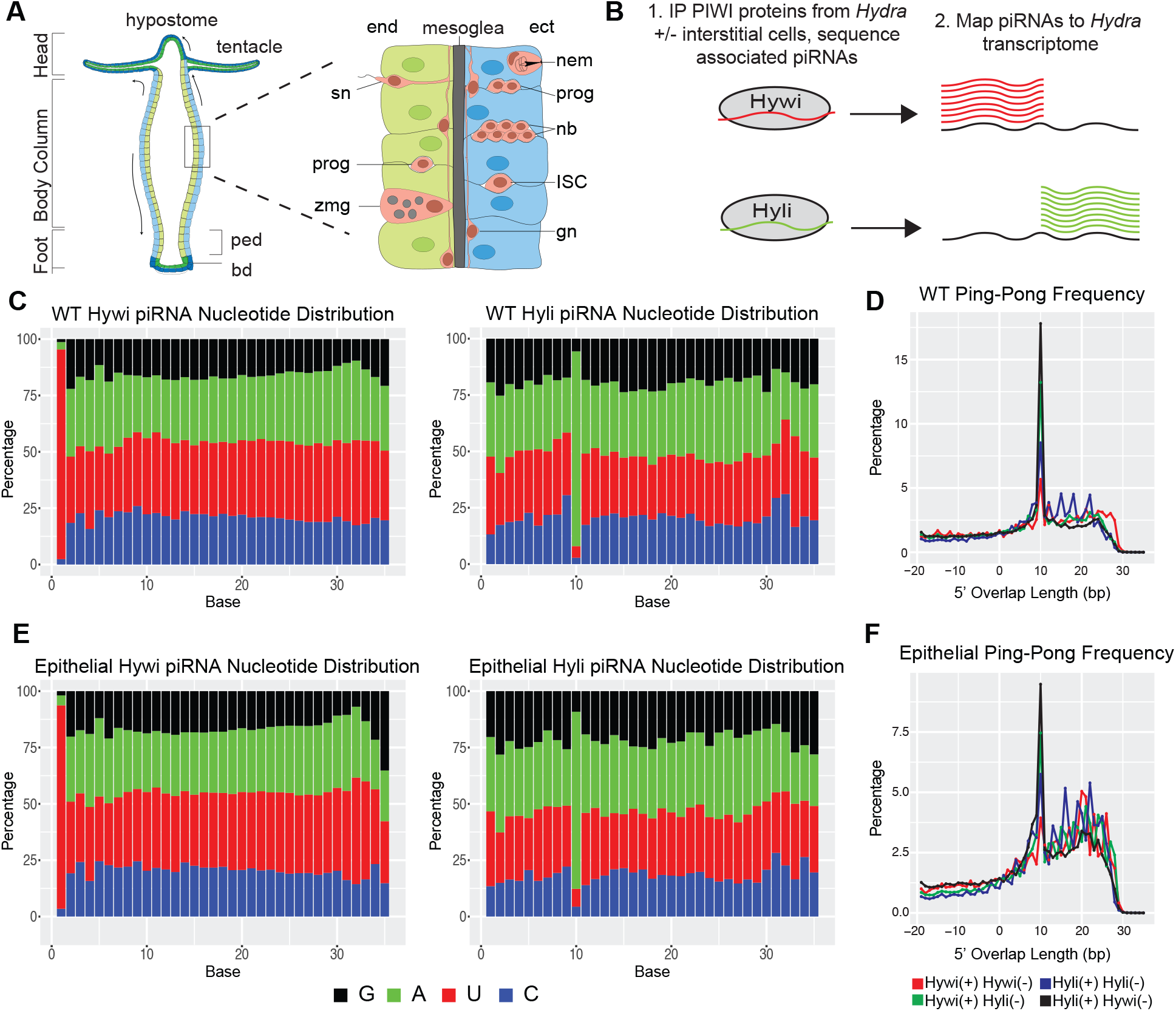
Epithelial *Hydra* piRNAs have sequence signatures typical of ping-pong biogenesis. (**A**) Schematic of the *Hydra* body plan and cell types adapted from (Siebert). *Hydra* is comprised of three cell lineages. The two epithelial cell lineages, endoderm (end, green) and ectoderm (ect, blue), form two monolayers separated by a basal lamina called the mesoglea. The cells of the interstitial lineage (pink) are embedded among the epithelial cells; this lineage is supported by a multipotent interstitial stem cell (ISC) which gives rise to three somatic cell types: neurons, gland cells, and nematocytes. Interstitial cells shown: progenitors (prog), nematoblasts (nb), nematocytes (nem), ganglion neuron (gn), sensory neuron (sg), and zymogen gland cell (zmg). The ISCs are also able to produce germline stem cells (not shown). By contrast, the epithelial cell lineages are strictly somatic. The epithelial cells of the body column (shown in lighter colors) are mitotically active, unipotent stem cells that express PIWI proteins (Juliano et al. 2014). Due to mitotic divisions, these cells are translocated towards the oral and aboral ends of the animal. When epithelial cells arrive at the extremities (shown in darker colors), the cells exit the cell cycle, lose PIWI expression, and terminally differentiate to build the hypostome and tentacles at the aboral end and the peduncle (ped) and basal disc (bd) at the aboral end. (**B**) Experimental design for Figures 1 and 2: piRNAs were extracted from either untreated (WT, germline-competent) or colchicine-treated (epithelial) *Hydra* by immunoprecipitation using Hywi- and Hyli-specific antibodies (Juliano et al. 2014). piRNAs were isolated from immunoprecipitated complexes, sequenced, and mapped to the *Hydra* transcriptome (Siebert et al. 2019). (**C,E**) Nucleotide distribution of Hywi- and Hyli-bound piRNAs isolated from (**C**) untreated wild type (WT) *Hydra* and (**E**) colchicine-treated epithelial *Hydra*. Hywi-bound piRNAs show a strong bias for a uridine at the 5’ end (1U), indicative of primary piRNAs. Hyli-bound piRNAs show a strong bias for an adenine 10 bases from the 5’ end (10A), indicative of secondary piRNAs formed via ping-pong biogenesis. (**D,F**) The frequency of complementary overlap between the 5’ ends of Hywi- and Hyli-bound piRNAs; all four possible combinations were tested. The high frequency of 10 base pair overlap between Hywi- and Hyli-bound piRNAs is indicative of ping-pong biogenesis. This signal is present in piRNAs isolated from both WT and epithelial *Hydra*, indicating that the ping-pong biogenesis pathway is active in ESCs.

*Hydra* has two cytoplasmic PIWI proteins, Hywi and Hyli, which are located in perinuclear, cytoplasmic granules found in ESCs and ISCs (Juliano et al. 2014; Lim et al. 2014). Hywi and Hyli exhibit the conserved “ping-pong” piRNA biogenesis pathway found in the germline. A large fraction of Hywi-bound piRNAs map to TEs in an antisense orientation and Hyli-bound piRNAs map to TEs in a sense orientation. *Hydra* piRNAs exhibit a typical “ping-pong signature,” which consists of a 10 base-pair overlap between the 5’ end of Hywi-bound piRNAs (primary piRNAs) and the 5’ end of Hyli-bound piRNAs (secondary piRNAs) (Juliano et al. 2014; Lim et al. 2014). These data indicate that Hywi and Hyli likely participate in ping-pong mediated cleavage of TE RNAs, a conserved mechanism that occurs in the cytoplasmic granules of germ cells (Brennecke et al. 2007; Aravin et al. 2008). Hywi knockdown in *Hydra* ESCs leads to loss of epithelial integrity and death within 12 days, thus demonstrating a necessary function for the PIWI-piRNA pathway in *Hydra* somatic stem cells (Juliano et al. 2014). However, our previous study focused on piRNAs isolated from whole animals and did not directly test the identity of PIWI-piRNA pathway targets in ESCs. Therefore, the identity of PIWI-piRNA targets in the strictly somatic ESCs is unknown and is the subject of this study.

Here we report that a primary target of the PIWI-piRNA pathway in *Hydra* ESCs are TE RNAs. This targeting occurs through the conserved ping-pong mechanism, which couples piRNA biogenesis to TE destruction (i.e. the RNA transcripts of TEs are processed into piRNAs). This is supported by our findings that TE-derived piRNAs are prevalent in *Hydra* ESCs and that TE transcripts are upregulated in response to epithelial *hywi* knockdown. Using degradome sequencing, we find evidence of PIWI-directed cleavage products of TE transcripts in somatic cells using the ping-pong mechanism, and we demonstrate a direct interaction between Hywi protein and target TE RNAs in ESCs. These observations demonstrate that the PIWI-piRNA pathway represses TEs in the somatic stem cells of *Hydra*, which suggests a role for the pathway in maintaining long-lived somatic cell lineages.

## Results and Discussion

### Hydra vulgaris AEP piRNAs have conserved sequence characteristics

In our previous study, we isolated and sequenced Hywi-bound and Hyli-bound piRNAs from *Hydra vulgaris* strain 105 and mapped these to genome and transcriptome assemblies generated from the same strain. These data suggested that the *Hydra* PIWI-piRNA pathway targets TEs through the conserved ping-pong mechanism (Juliano et al. 2014). However, we were not able to discern cell lineage-specific roles for the pathway because piRNAs were isolated from whole *Hydra vulgaris* strain 105. TE repression is a conserved function of the PIWI-piRNA pathway in germ cells (Brennecke et al. 2007; Aravin et al. 2007; Carmell et al. 2007; Das et al. 2008; Houwing et al. 2007; Kuramochi-Miyagawa et al. 2008), but targets in somatic cells are less well understood (Ross et al. 2014). Therefore, in this study we aimed to identify targets of the pathway specifically in the ESCs.

To understand lineage-specific functions of the PIWI-piRNA pathway in *Hydra*, we used transgenic lines with cell-lineage or cell-type specific expression of fluorescent proteins. However, our previous study of *Hydra* PIWI proteins and piRNAs was done using *Hydra vulgaris* strain 105, which does not produce eggs in the laboratory and cannot be used to make transgenic lines. By contrast, *Hydra vulgaris* strain AEP produces eggs in the laboratory and is routinely used for the production of transgenic lines (Wittlieb et al. 2006). Therefore, we began this study by testing if the two *Hydra* strains have the same piRNA characteristics. We first established that our Hywi and Hyli antibodies are able to immunoprecipitate PIWI-piRNA complexes in the AEP strain (**Fig S1A-D**). Next, we isolated and sequenced Hywi-bound and Hyli-bound piRNAs from the AEP strain and mapped these to our AEP transcriptome (**Fig. 1B**) (Siebert et al. 2019). To understand piRNA mapping preferences we categorized transcripts as: 1) TE transcripts (“TE”), 2) protein-coding transcripts with annotations (“gene”), 3) unannotated transcripts with open reading frames of at least 100 amino acids (“uncharacterized”), 4) and non-coding transcripts with open reading frames of less than 100 amino acids (“ncRNA”) (see Methods for further details on transcript annotations). The “uncharacterized” transcripts could include TEs that we were unable to annotate, we therefore kept this group as a separate category.

Similar to our previous data from the 105 strain (Juliano et al. 2014; Lim et al. 2014), we found evidence for ping-pong mediated TE repression in the AEP strain: 1) 93% of Hywi-bound piRNAs (primary piRNAs) have a uridine at the 5’ end and 86% of Hyli-bound piRNAs (secondary piRNAs) have an adenine at the 10th base from the 5’ end (**Fig. 1C**), 2) The presence of a 10 base-pair overlap between Hywi-bound and Hyli-bound piRNAs (**Fig. 1D**), and 3) piRNAs map to TE transcripts at a higher density than non-TE transcripts (**Fig. 2A**; **Table S1**, see Methods for details on transcript annotation). Therefore, we concluded that the PIWI-piRNA pathway functions similarly in the two commonly used *Hydra* strains. Subsequent experiments in this study were done with *Hydra vulgaris* strain AEP, hereafter referred to simply as “*Hydra*.”

**Figure 2.**
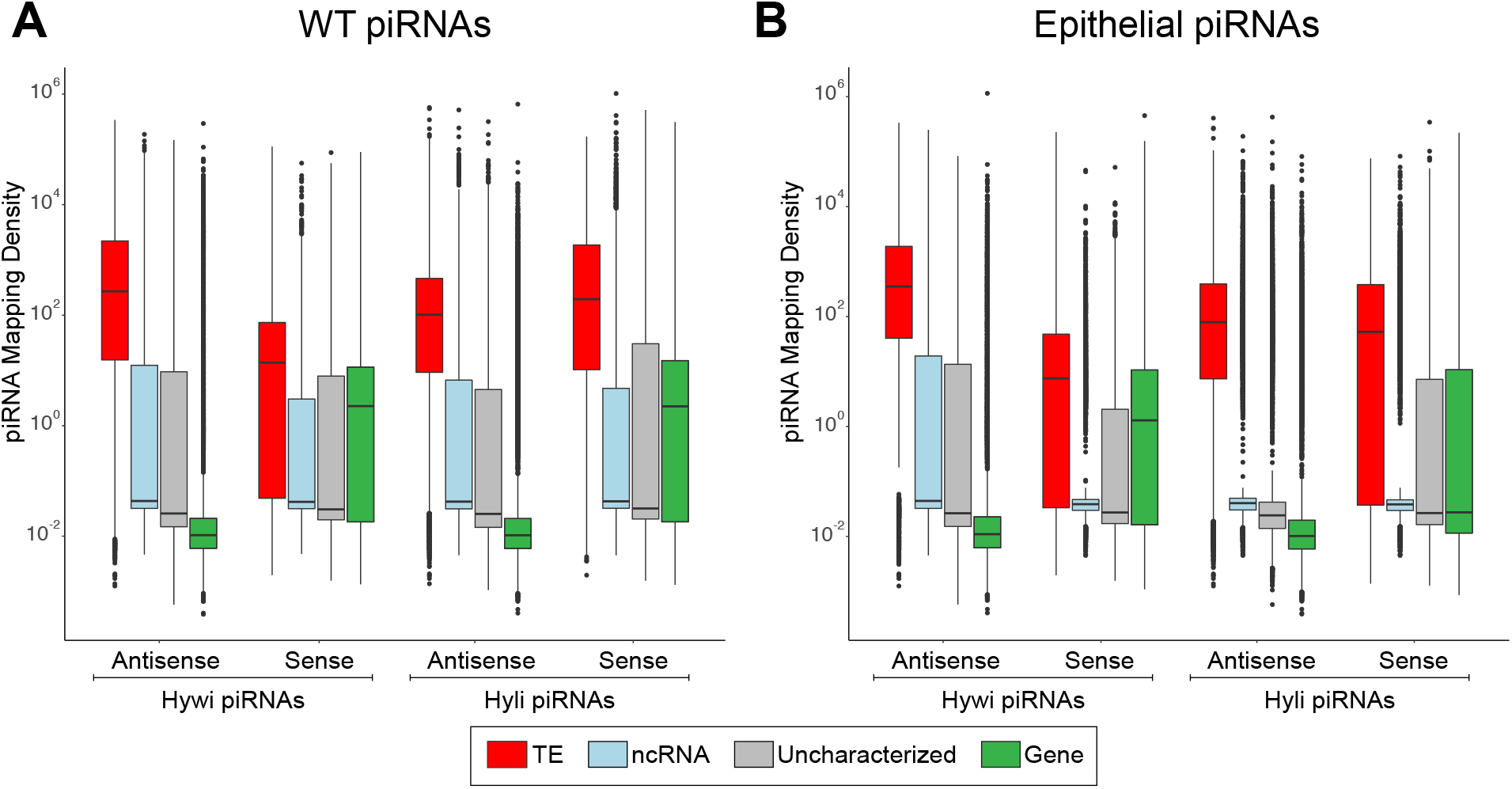
Epithelial piRNAs map predominantly to TE transcripts. Hywi- and Hyli-bound piRNAs isolated from (**A**) whole *Hydra* and (**B**) epithelial *Hydra* were mapped to the transcriptome. The piRNA mapping density (mapped piRNA counts per kb per million counts) is shown on the Y-axis of the box and whisker plots. The X-axis shows results subdivided by: 1) Hywi- or Hyli-bound piRNAs, 2) Mapped in a sense or antisense orientation, and 3) Transcript class (see legend). In all cases, TEs have a significantly higher mapping density than other transcript classes as determined using Tukey’s range test (p<0.001; Supplementary Analysis 1). For TE transcripts, the high density of antisense mapped Hywi-bound piRNAs and sense mapped Hyli-bound piRNAs is expected due to the mechanism of ping-pong piRNA biogenesis. Particularly for piRNAs isolated from WT *Hydra*, the relatively higher density of sense mapping piRNAs to gene transcripts as compared to non-coding RNA transcripts and uncharacterized transcripts could indicate primary processing of gene transcripts.

### Hydra piRNAs isolated from epithelial stem cells are enriched for TE sequences

To gain insight into the function of the PIWI-piRNA pathway in *Hydra* ESCs, we first isolated and analyzed piRNAs from epithelial *Hydra*, which do not contain any cell types of the interstitial lineage. To obtain epithelial *Hydra*, we used an established colchicine treatment protocol to remove ISCs {Campbell:1976wa}, which leaves ESCs as the only PIWI-expressing cell type. To confirm ISC depletion in our hands, we performed colchicine treatments using a transgenic line expressing GFP under the control of the promoter of *cnnos1*, a gene expressed in ISCs (Hemmrich et al. 2012). Animals were treated with 0.4% colchicine for 8 hours followed by a 10-day recovery period. Immunoblot analysis confirmed the loss of GFP-positive ISCs (**Fig. S1E**). Given that stress could lead to increased expression of TEs (Horváth et al. 2017), we assayed TE abundance in colchicine-treated animals by qPCR. We found that neither *hywi* expression or TE expression is increased in colchicine-treated animals, therefore TE-derived piRNAs are not artificially abundant in the epithelial *Hydra* used in this study (**Fig. S1F**). Epithelial *Hydra* obtained from colchicine treatment lack the interstitial lineage but can be cultured if fed by hand and show no deficits in asexual reproduction or regeneration (Marcum and Campbell 1978). In summary, the epithelial lineages are able to return to normal function after the colchicine treatment, thereby providing a suitable mechanism for studying PIWI-piRNA pathway function in the ESCs.

To isolate the piRNAs expressed in epithelial cells, we performed Hywi and Hyli immunoprecipitations from lysates prepared from colchicine-treated *cnnos1*∷GFP *Hydra* (**Fig. 1B**). The isolated piRNAs were sequenced and the base distribution across the length of the piRNAs was analyzed. Similar to piRNAs isolated from whole animals, 90% of Hywi-bound piRNAs isolated from epithelial animals had a uridine at the 5’ end and 79% of Hyli-bound piRNAs had an adenine at the 10th base from the 5’ end (**Fig. 1E**). In addition, epithelial piRNAs showed the 10 base pair overlap consistent with ping-pong piRNA biogenesis (**Fig. 1F**). The strongest 10 base pair overlap was found when comparing sense Hyli piRNAs and antisense Hywi piRNAs as expected. However, all possible combinations showed some degree of 10 base pair overlap, which suggested that the ping-pong mechanism can occur with any combination of PIWI proteins including homotypic ping-pong, which has also been documented in *Drosophila*, mice, and mollusks (Aravin et al. 2008; Zhang et al. 2012; Jehn et al. 2018). Overall, this demonstrates that piRNAs expressed in the ESCs are processed by the ping-pong biogenesis pathway (Brennecke et al. 2007) (Iwasaki et al. 2015).

To identify PIWI-piRNA pathway RNA targets in ESCs, we mapped epithelial piRNAs to a *Hydra vulgaris* AEP stranded transcriptome (**Fig. 1B**) (Siebert et al. 2019). We considered both possible mapping orientations for piRNAs, sense or antisense with respect to the transcript. piRNAs mapping in an antisense orientation could indicate binding of PIWI-piRNA complexes to an RNA target, which could ultimately lead to cleavage of that target by the “slicer” activity of the PIWI domain (Nishida et al. 2007; Gunawardane et al. 2007; Saito et al. 2006). piRNAs mapping in a sense orientation could indicate that the RNA target has been directly processed into piRNAs by PIWI proteins either by primary or secondary (i.e. ping-pong) piRNA biogenesis (Iwasaki et al. 2015; Brennecke et al. 2007). Therefore, for antisense mapping we allowed 3 mismatches because cleavage of RNA targets can occur with imperfect base-pairing (Zhang et al. 2015), but for sense mapping, we did not allow any mismatches under the assumption that piRNAs derived directly from an RNA target should have the identical sequence. Using this mapping strategy, we found that piRNAs isolated from epithelial animals largely have the same characteristics as piRNAs isolated from whole animals (**Fig. 2**; **Table S1**). We found that for both antisense and sense mapping, both Hywi-bound and Hyli-bound piRNAs map to TE transcripts at a significantly higher density than to non-TE transcripts (**Fig. 2**). This was likely a reflection of TE RNAs being processed into piRNAs by the ping-pong mechanism, which appears to occur between all possible combinations of PIWI protein pairs (**Fig. 1D,F**). For antisense mapping, most non-TE transcripts had a low mapping density. However, for both the Hywi-bound and Hyli-bound antisense mapping, there were groups of gene transcripts with high mapping density that could indicate the existence of gene targets (**Fig. 2**). Sense mapping of Hywi-bound piRNAs to gene transcripts may indicate primary processing of these transcripts (**Fig. 2**). A higher density of Hyli-bound sense mapping was observed for piRNAs isolated from whole animals than for piRNAs isolated from ESCs (**Fig 2**). This could indicate that Hyli-directed primary processing of gene transcripts is more prevalent in the interstitial lineage (**Fig. 2A**) as compared to the epithelial lineages (**Fig. 2B**). In summary, these data suggest that TEs are a major target of the PIWI-piRNA pathway in ESCs. In addition, this mapping strategy has identified putative non-TE targets that will be further addressed below.

To further test the lineage-specific functions of the PIWI-piRNA pathway, we analyzed total *Hydra vulgaris* AEP small RNAs that were isolated and sequenced from the ectodermal, endodermal, and interstitial lineages in our previous study (Juliano et al. 2014). This allowed us to determine the characteristics of piRNAs isolated from each lineage separately. We identified the piRNAs in the small RNA data sets by cross-referencing with the piRNAs sequenced from whole *Hydra vulgaris* AEP animals in this study (**Fig. S2A**). Using this strategy, we produced a list of unique piRNA species and annotated the lineage(s) of origin for each sequence (**Fig. S2B**). We next mapped the epithelial piRNAs to the *Hydra* transcriptome and, similar to our results from epithelial animals, we found an enrichment for piRNAs mapping to TE transcripts as compared to other transcript categories (**Fig. S2C**). In addition, this strategy allowed us to identify and map piRNAs specific to the interstitial lineage and we also found an enrichment for piRNAs mapping to TE transcripts as compared to other transcript categories (**Fig. S2D**). An enrichment for transposon sequences in piRNAs expressed specifically in ISCs is expected because ISCs have germline potential.

### RNAi knockdown of hywi in epithelial stem cells leads to the upregulation of both TE and non-TE transcripts

Transcripts that are targeted by the PIWI-piRNA pathway should be upregulated in response to down regulation of PIWI genes. To identify such transcripts, we used our previously validated *hywi* RNAi-1 transgenic line, which constitutively expresses a *hywi* hairpin under the control of an *actin* promoter that is not active in ISCs (Juliano et al. 2014). The hairpin construct also includes DsRed2, which is used to track transgenic tissue. The hairpin construct is integrated only in the interstitial lineage and is maintained through asexual propagation. In the asexually propagated *hywi* RNAi-1 transgenic line, the hairpin is not expressed in the PIWI-positive ISCs and therefore does not have a negative impact on the viability of the line. Integration of the hairpin construct in the interstitial lineage allows for germline propagation. After sexual reproduction, the transgene is now integrated into the genome of all cells in the offspring and the actin promoter drives expression in all epithelial cells and differentiated interstitial cells; the ESCs are the only *piwi*-positive cells that have transgene expression. Therefore, we can analyze the effect of knocking down *hywi* in the ESCs in the F1 generation. Using this strategy, we previously found that *hywi* knockdown in ESCs leads to death within 12 days of hatching (Juliano et al. 2014). In this study we performed RNA-seq and differential gene expression (DGE) analysis to compare gene expression between wild type offspring (did not inherit the RNAi transgene) and *hywi* RNAi offspring (did inherit the transgene) 4 days after hatching. We identified 441 transcripts from our *Hydra* transcriptome that were upregulated in *hywi* RNAi animals as compared to wild type siblings (**Fig. 3A**; **Table S1, S2**).

**Figure 3.**
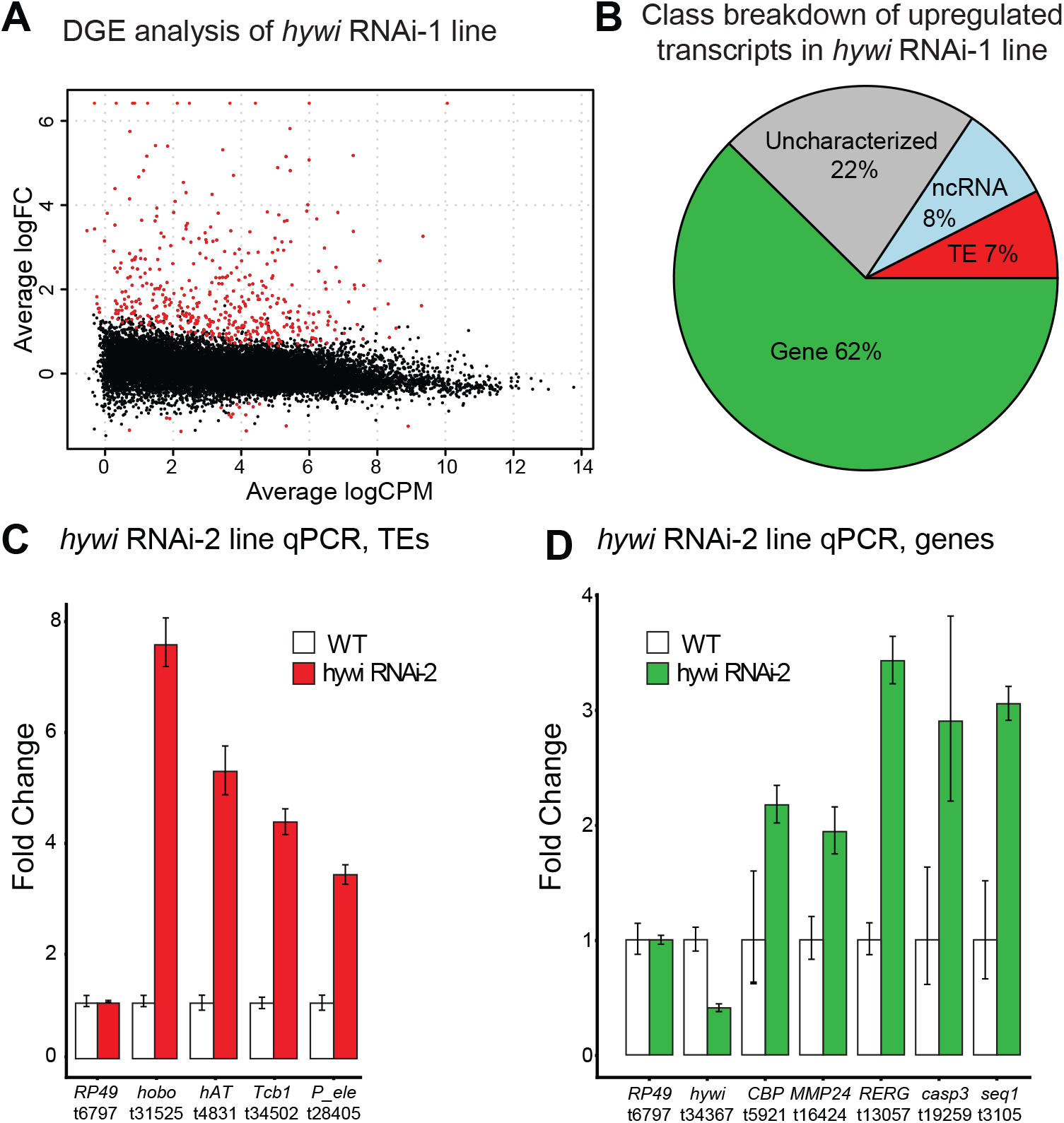
Both TE and non-TE transcripts are upregulated in response to epithelial *hywi* RNAi. (**A**) Differential gene expression (DGE) analysis was performed to compare transcript expression levels between *hywi* RNAi animals and wild type siblings; knockdown animals carry the *hywi* RNA-1 transgene (Juliano et al. 2014). Volcano plot showing 458 differentially expressed transcripts (red); 441 transcripts are upregulated and 17 transcripts are downregulated (adjusted p-value < 0.05). (**B**) Transcript class breakdown of the 441 transcripts upregulated in response to *hywi* knockdown. (**C,D**) A second RNAi transgenic line (*hywi* RNAi-2) was used to validate the DGE results (Juliano et al. 2014). qPCR was used to test the levels of select (**C**) TE transcripts and (**D**) gene transcripts identified by the DGE performed on the *hywi* RNAi-1 transgenic line (**A**). These results indicate that transcripts upregulated in the hywi RNAi-1 transgenic line are also upregulated in the hywi RNAi-2 transgenic line, demonstrating the specific effect of *hywi* knockdown.

Given that epithelial piRNAs were enriched for TE sequences, we predicted that TEs would be upregulated upon RNAi knockdown of *hywi* in ESCs. However, among the 441 transcripts upregulated in response to *hywi* knockdown only 7% (33 out of 441) were annotated as TEs (**Fig. 3B**). To gain insights into the identity of these transcripts that were upregulated in response to *hywi* knockdown (either due to direct or indirect effects), we performed GO-term analysis. We found seven terms (biological process) significantly enriched, including “defense response” and “innate immune response” (adjusted p-value < 0.05; Supplementary Analysis 2). This result could suggest that TE de-repression triggers an innate immune response in the affected cells, a phenomenon which has been observed in other organisms (Simon et al. 2019; Cecco et al. 2019).

To test the reproducibility of the DGE results, we used a second *hywi* RNAi line (*hywi* RNAi-2), which targets a different portion of the *hywi* gene. We generated this transgenic line using the same strategy as the *hywi* RNAi-1 line (Juliano et al. 2014). RNA was extracted from both wild type and *hywi* RNAi hatchlings produced from the *hywi* RNAi-2 parental line. qPCR analysis of these samples was used to test the expression levels of select transcripts identified by the DGE analysis (**Fig. 3A**). These data confirmed that many of the same transcripts were upregulated in *hywi* RNAi animals produced from both the *hywi* RNA-1 and *hywi* RNA-2 lines (**Fig. 3C, D**). This indicates that the results were due to RNAi knockdown of *hywi* and not off target effects of the *hywi* hairpins.

Our DGE analysis raises the possibility that the PIWI-piRNA pathway has many non-TE targets in ESCs, but it is also possible that these are upregulated due to secondary effects of TE upregulation. To help discern between these two scenarios, we asked how many of the upregulated transcripts have a high density of piRNAs mapping to them, which could indicate direct targeting by the PIWI-piRNA pathway. To identify transcripts with the highest density of mapped piRNAs, we ordered transcripts from the highest to lowest density of mapped epithelial Hywi piRNAs, and defined transcripts in the top 20% as “high mapping” (**Table S2**). Of the 441 upregulated transcripts, we found the following percentages to be “high mapping” in each category: 1) 93.9% (31 out of 33) for TE transcripts, 2) 13.9% (5 out of 36) for non-coding transcripts, 3) 25.4% (34 out of 134) for uncharacterized transcripts, and 4) 15.1% (36 out of 238) for gene transcripts. Therefore, we found that TE transcripts were overrepresented in the “high mapping” category. In addition, we identified 36 genes that may be direct targets of the PIWI-piRNA pathway in ESCs because they were both in the “high mapping” category and were upregulated in response to *hywi* knockdown. Of these 36 genes, 27 may be targeted by primary processing because the mapped piRNAs were almost entirely in the sense orientation (**Table S3**).

We predicted that direct targets of the PIWI-piRNA pathway would have lower expression in stem cells and higher expression in differentiated cells. We therefore analyzed the homeostatic expression patterns of the 36 putative gene targets in the epithelial cells by interrogating single-cell expression data (Siebert et al. 2019). The data used in these visualizations included genes that were found to be expressed in at least 1% of the ectodermal or endodermal epithelial cells (Siebert et al. 2019). We found epithelial expression for 24 of the 36 putative gene targets: 1) nineteen gene transcripts were expressed in both the endodermal and ectodermal epithelial lineages, 2) one transcript was expressed only in the ectodermal epithelial lineage, and 3) four transcripts were expressed only in the endodermal epithelial lineage. No epithelial expression was found for 12 of the gene transcripts, which could indicate low expression in a homeostatic animal. This could be due to transcript degradation by the PIWI-piRNA pathway or an overall low level of transcription. For the 24 gene transcripts for which we had expression data, we generated plots to visualize expression profiles in epithelial cells along the oral-aboral axis (Supplementary Analysis 5). We inspected these plots for expression patterns that could suggest targeting by the PIWI-piRNA pathway: higher expression in the PIWI-negative cells at the extremities as compared to the PIWI-positive cells in the body column. Of the 24 transcripts, the expression pattern of t14391 was the most suggestive of Hywi-mediated repression, with higher expression at the extremities in both the ectoderm and endoderm (**Fig. S3**). Swissprot Blast analysis of this transcript suggested similarity to *PARP-12*. Interestingly, PARP family genes are involved in stress response, including DNA damage repair, making this an interesting putative target for future study (Bai 2015).

### PIWI-piRNA complexes cleave TE transcripts in epithelial stem cells

Our analysis of piRNA sequences isolated from ESCs strongly suggests that Hywi and Hyli repress TE RNAs by processing these RNAs into piRNAs using the ping pong mechanism. To further test this, we looked for evidence of cleaved transposable element RNAs using degradome sequencing, a strategy used to isolate and sequence cleaved RNAs by selecting for uncapped RNAs with a poly(A) tail (Addo-Quaye et al. 2008; German et al. 2008). To determine the RNA targets cleaved by PIWI proteins in *Hydra*, we generated degradome sequencing libraries from both untreated and colchicine-treated *cnnos1*∷GFP *Hydra*. Cleaved RNA targets of the PIWI-piRNA pathway display the ping-pong signature, which in *Hydra* typically manifests as a 10-bp overlap between the 5’ end of antisense-oriented Hywi-bound piRNAs with the 5’ end of sense-oriented Hyli-bound piRNAs and degradome fragments (**Fig. 4A**). We considered genes that have 10 or more of each of these species mapped and aligned in a “ping-pong” signature to be RNA targets of the PIWI-piRNA pathway. We identified 2,118 such targets in wild type animals and 251 such targets in epithelial animals (**Table S4**). In libraries produced from both whole animals and epithelial animals, TEs comprised the highest percentage of transcripts displaying this signature (**Fig. 4B**). The second highest category was uncharacterized transcripts, which may include some TE transcripts that we were unable to annotate. Of the 441 transcripts upregulated in response to *hywi* knockdown, 47 were identified as targets in the wild type degradome library (**Table S4**). The majority of these (39 out of 47) were either TEs (22) or uncharacterized (17). Three gene transcripts were upregulated in response to *hywi* RNAi, had a high number of piRNAs mapping, and were also identified as targets in the degradome analysis (**Table S3**). In summary, the degradome results indicated that a major function of the PIWI-piRNA pathway in *Hydra* ESCs is to cleave TE transcripts, but also revealed some possible non-TE targets for future study.

**Figure 4.**
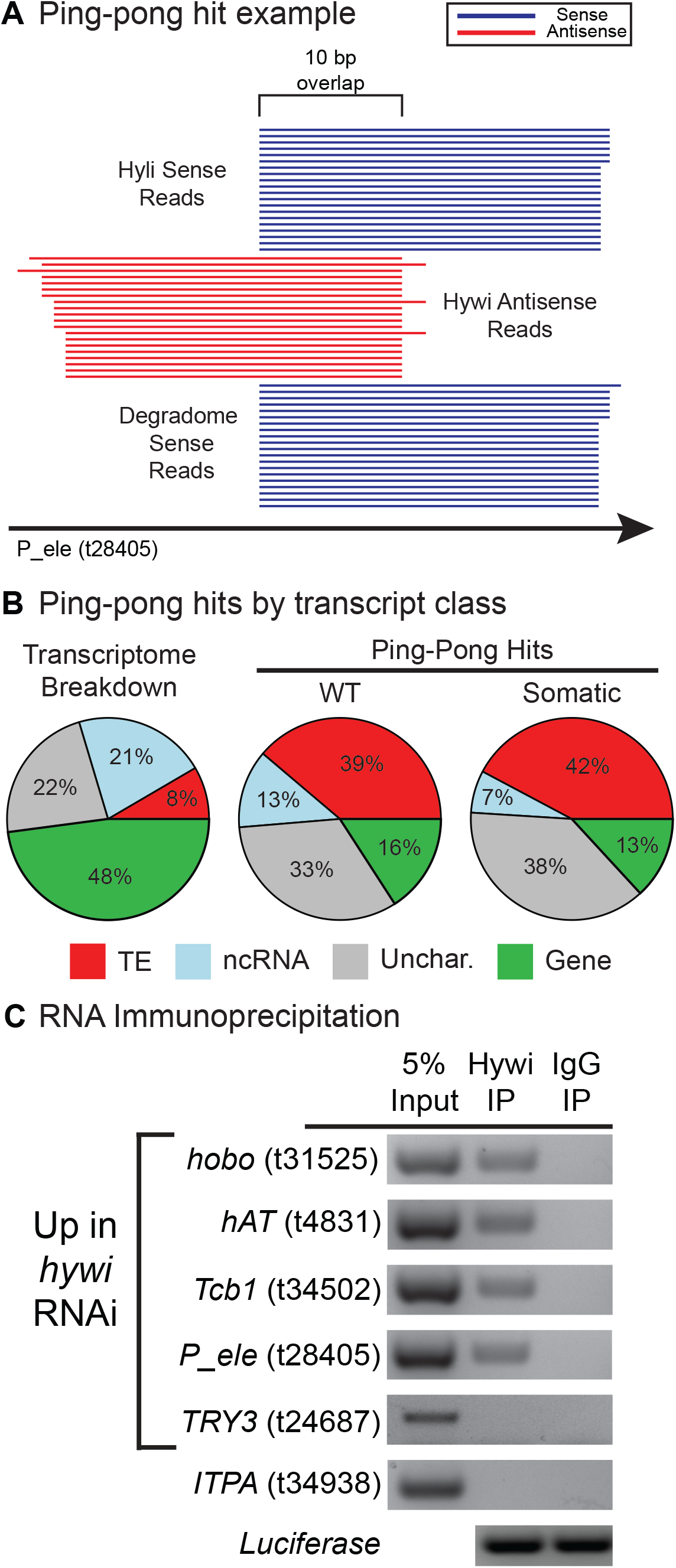
TE transcripts are cleaved by PIWI-piRNA complexes in *Hydra* epithelial stem cells. (**A**) Hywi-bound piRNAs, Hyli-bound piRNAs, and degradome reads were aligned to the transcriptome to identify likely direct targets of the PIWI-piRNA pathway. One example from the data is shown (t28405). The following pattern indicates cleavage: Antisense-oriented Hywi-bound piRNAs align with a 10-nucleotide 5’ overlap with both sense-oriented Hyli-bound piRNAs and sense-oriented degradome reads. For a transcript to be considered a target of the PIWI-piRNA pathway, we require a minimum of 10 reads of each species to map in such an arrangement. (**B**) The degradome sequencing from wild type (WT) *Hydra* identified 2118 transcripts as targets of the PIWI-piRNA pathway and the transcript class distribution of these targets is shown. The degradome sequencing from epithelial *Hydra* identified 251 transcripts (**Table S4**) as targets of the PIWI-piRNA pathway and the class distribution of these targets is shown. In contrast to the transcriptome as a whole, TE transcripts and uncharacterized transcripts comprise the majority of targets in both whole animals and epithelial animals. (**C**) RNA immunoprecipitation (RIP) was used to identify transcripts that directly interact with Hywi-piRNA complexes in somatic stem cells (for complete results see **Fig S4**). Following RIP, luciferase mRNA was added to the washed beads to act as a positive control for RNA extraction and cDNA synthesis. RT-PCR was used to test for the presence of target transcripts in complexes immunoprecipitated with a Hywi antibody. The results for four TE transcripts are shown, which were amplified in three biological replicates (**Fig. S4**). The following transcripts did not associate with Hywi protein: 1) The *TRY3* transcript is upregulated in response to *hywi* RNAi but has a low piRNA mapping density and 2) the *ITPA* transcript did not change in response to *hywi* RNAi and has a low piRNA mapping density.

Finally, we tested whether Hywi-piRNA complexes physically associate with the somatic RNA targets identified by piRNA mapping, DGE analysis of *hywi* RNAi animals, and/or degradome sequencing. To this end, we performed RNA-immunoprecipitation (RIP) to isolate Hywi-piRNA-RNA ternary complexes from epithelial *Hydra* (Ilyin et al. 2017). We performed RT-PCR to test for the presence of the following transcripts in our immunoprecipitated complexes: 1) eight TE transcripts that were upregulated in response to *hywi* knockdown, had a high number of piRNAs mapping, and were identified as targets in the degradome sequencing; 2) seven gene transcripts that were upregulated in response to *hywi* knockdown and had a high number of piRNAs mapping (one of these was identified as a target in the degradome sequencing) (**Table S3**); 3) three gene transcripts that were upregulated in response to *hywi* knockdown, but did not have a high number of piRNAs mapping; and 4) six gene transcripts that were not affected by *hywi* knockdown and did not have a high number of piRNAs mapping to act as negative controls. We could reproducibly pull down four of the eight TE transcripts in all three biological replicates; the remaining four could be pulled down in at least one replicate. These data demonstrated a physical interaction between Hywi-piRNA complexes and TE RNAs (**Fig. 4C**; **Fig. S3**). By contrast, we could not reproducibly pull down transcripts from the remaining categories listed above (**Fig. S4**). Three of the seven gene transcripts from the second category (t27033, t8142, and t5862) were pulled down in one or two of the three biological replicates (**Fig. S3**; **Table S3**). These three genes did not appear in the degradome sequencing, but rather may be a product of primary processing, a ping-pong independent mechanism which produces only sense oriented piRNAs, as indicated by a large number of sense mapping Hywi-bound piRNAs (**Table S1**) (Homolka et al. 2015; Pandey et al. 2017).

## Conclusion

TE expression is upregulated in aging metazoan somatic tissues and in aging yeast, suggesting that TE de-repression is a conserved feature of cellular aging (Maxwell et al. 2011; Cecco et al. 2013; Li et al. 2013; Van Meter et al. 2014; Patterson et al. 2015; Wood et al. 2016). Some studies show that repressing TEs can extend lifespan, thus understanding the mechanisms by which long-lived organisms repress TEs is of interest (Wood et al. 2016; Jones et al. 2016; Wang et al. 2011; Cecco et al. 2019; Simon et al. 2019). Here we show that the PIWI-piRNA pathway represses TEs in the somatic stem cells of the long-lived *Hydra*. This, together with our previous finding that loss of *hywi* function in ESCs leads to loss of epithelial function and death (Juliano et al. 2014), leads us to propose that the PIWI-piRNA pathway contributes to the longevity of *Hydra* somatic stem cells. Furthermore, we demonstrate direct targeting of TE RNAs by the pathway which is accompanied by the upregulation of genes involved in immune response. This suggests a model in which TE upregulation triggers an immune response in *Hydra*, similar to recent reports in mammals (Cecco et al. 2019; Simon et al. 2019; Capone et al. 2018; Benayoun et al. 2018).

Although the PIWI-piRNA pathway is best studied in the germline, our data in combination with several studies demonstrates a wider role in somatic tissues (Ross et al. 2014). For example, recent studies in mollusks and arthropods found evidence for widespread somatic ping-pong as a transposon repression mechanism (Jehn et al. 2018; Lewis et al. 2018). *Drosophila piwi* is required to repress TE expression in the fat body and in intestinal stem cells. In these somatic tissues, *piwi* expression has been linked to somatic longevity (Jones et al. 2016; Sousa-Victor et al. 2017). For example, in aging flies, TEs are upregulated in intestinal stem cells and overexpression of *piwi* prevents the age-associated decline of intestinal stem cell function (Sousa-Victor et al. 2017). It is however not clear why *piwi* fails to be effective in the somatic tissues of aging *Drosophila*, whereas *hywi* and *hyli* appear to indefinitely protect somatic cells in *Hydra*. PIWI genes are expressed in planarian neoblasts and HeLa cells, where they could also contribute to the maintenance of somatic immortality (Reddien et al. 2005; Palakodeti et al. 2008; Lu et al. 2010). In conclusion, we find that the PIWI-piRNA pathway represses TE expression through a ping-pong mechanism in the somatic tissue of a cnidarian, the sister group to bilaterians, suggesting that this is an ancient TE repression mechanism that may contribute somatic longevity.

## Materials and Methods

### Code Availability

Analysis code, scripts and supplemental analysis files that are needed to reproduce the analyses are available in a git repository: https://github.com/cejuliano/bteefy_piwi_transposon

1_piRNA_and_Degradome_Counts.Rmd Generation of piRNA and degradome count files, transcript classification
2_Differential_Gene_Expression_GO_Analysis.Rmd hywi knockdown - Differential Gene Expression analysis, GO-term enrichment
3_Ping_Pong_Analysis.Rmd Ping-pong hit count files, ping-pong overlap frequency, ping-pong hit classification
4_Lineage_Sorted_piRNA_Mapping.Rmd Lineage-sorting and counting of piRNAs
5_Single_Cell_Data_Exploration.Rmd Assessment of homeostatic epithelial expression of putative hywi targets

### Data Availability

Raw reads from piRNA sequencing, degradome sequencing, and lineage-sorted mRNA sequencing have been submitted to the GEO repository GSE135440.

### Transgenic strains used in this study

Interstitial stem cell depletion was performed on cnnos1∷GFP *Hydra* (Hemmrich et al. 2012). Differential Gene Expression (DGE) analysis was performed on our previously published *hywi* RNAi-1 animals (targeting bases 379-899 of hywi) (Juliano et al. 2014). Validation was done by creating a second transgenic line using our previously published *hywi* RNAi-2 transgene (targeting bases 1557-2093 of *hywi*) (Juliano et al. 2014). *Hydra* were cultured at 18°C and fed *Artemia nauplii* 1-3 times per week.

### Colchicine-treatment to remove the interstitial lineage

Colchicine treatment was performed as previously described (Campbell 1976). In brief, cnnos1∷GFP *Hydra* (Hemmrich et al. 2012) were exposed to 0.4% colchicine dissolved in *Hydra* medium (HM) (1 mM CaCl2, 0.1 mM MgCl2, 0.03 mM KNO3, 0.5 mM NaHCO3, 0.08 mM MgSO4) for 8 hours. After exposure, animals were rinsed with HM and transferred to fresh HM. Replicates were kept in separate dishes and HM was changed every day for 10 days after colchicine exposure while animals recovered. Ten animals were used per RNA immunoprecipitation experiment. To create degradome libraries, RNA was extracted from 10 *Hydra* per replicate using Trizol (ThermoFisher Scientific) according to the manufacturer’s instructions. Interstitial cell depletion was assayed using qPCR to detect levels of *Cnnos1* and *GFP* transcripts using cDNA synthesized from 1 μg of Trizol-extracted RNA. Interstitial cell depletion was also tested using an immunoblot to detect levels of GFP from three animals (see below for immunoblot details).

### Immunoprecipitation of piRNAs with PIWI antibodies

Immunoprecipitation (IP) of PIWI proteins for piRNA isolation and sequencing was done using either 225 untreated *Hydra vulgaris* AEP or 225 colchicine-treated cnnos1∷GFP *Hydra*. *Hydra* were centrifuged at low speed, HM was removed, and animals were resuspended in 1 mL MCB buffer (50 mM HEPES, pH 7.5, 150 mM potassium acetate, 2 mM magnesium acetate, 10% glycerol, 0.1% Triton X-100, 0.1% NP-40, 1 mM DTT). The following was added fresh to the MCB buffer before use: 1) 1 tablet/10 mL Pierce™ Protease Inhibitor Mini Tablets, EDTA-free (ThermoFisher Scientific) and 2) 1 U/*μ*L RNaseOUT™ Recombinant Ribonuclease Inhibitor (ThermoFisher Scientific). *Hydra* polyps in MCB buffer were homogenized on ice using a Dounce homogenizer. The resulting lysate was centrifuged at 20,000xg for 10 mins at 4°C and the supernatant was collected. Lysate was pre-incubated with 200 *μ*L of rehydrated Protein A Sepharose CL-4B beads (GE Healthcare, Life Sciences) for 15 minutes at 4°C with rocking; to rehydrate beads, 100 mg of beads were washed and resuspended in 300 *μ*L of MCB buffer. To remove beads, lysate was centrifuged at 20,000xg for 10 mins at 4°C and supernatant was collected. For each IP, 12 *μ*g of purified Hywi or Hyli antibody (Juliano et al. 2014) was added to 500 *μ*g of cleared protein lysate as measured by a NanoDrop ND-1000 Spectrophotometer 280 nM reading. The volume was then brought up to 1.2 mL per tube with MCB buffer. Next, the samples were incubated for 40 minutes with rocking at 4°C. Following antibody incubation, 60 *μ*L of rehydrated beads were added to the lysate and rocked at 4°C for an additional 40 minutes. Beads were collected by centrifugation at 500xg for 5 mins at 4°C and then washed 5 times with 1 mL of MCB buffer. After the washes, the bead volume was brought up to 100 *μ*L with MCB buffer and 10 *μ*L was removed for immunoblot blot analysis (see below for immunoblot details). To isolate piRNAs, 1 mL of Trizol was added to the remaining beads and RNA extraction was performed according to the manufacturer’s instructions. The RNA pellet was resuspended in 20 *μ*L of DEPC-treated water (Ambion). For size analysis, 8 *μ*L was removed for 5’-end labeling with [ɣ-32P] ATP using polynucleotide kinase. Labeled RNAs were run on a TBE-Urea gel. The remaining 12 *μ*L of RNA was used to prepare a sequencing library (see below for details on library preparation).

### Immunoblotting

To detect GFP protein in cnnos1∷GFP animals, 10 *Hydra* were homogenized in SDS-PAGE sample buffer for immunoblot analysis. Anti-GFP antibody was used at a 1:1000 dilution (Sigma, Catalog #11814460001 Roche). All antibodies were diluted in blocking solution (3% w/v powdered milk dissolved in 0.1% TBS-Tween). For detection of Hywi and Hyli protein, 8 *μ*g of purified anti-Hywi or anti-Hyli antibody in 50 mL of blocking solution was used for immunoblots. Anti-GFP was detected using goat anti-mouse IgG (H+L) Highly Cross-Adsorbed Secondary Antibody, Alexa Fluor Plus 800 (ThermoFisher Scientific, A32730). Anti-Hywi was detected using goat anti-Rabbit IgG (H+L) Superclonal™ Secondary Antibody, Alexa Fluor 680 (Thermofisher Scientific, A27042). Anti-Hyli was detected using goat anti-guinea pig IgG (H+L) Cross Adsorbed Secondary Antibody, DyLight 800 conjugate (ThermoFisher Scientific, SA5-10100). All secondary antibodies were diluted 1:10,000 in blocking solution. Immunoblots were imaged using the LI-COR Odyssey platform.

### piRNA and degradome sequencing library preparation

Sequencing libraries were prepared from immunoprecipitated piRNAs using the NEXTflex Small RNA-Seq Kit v3 (PerkinElmer Cat# NOVA-5132-05). The manufacturer’s protocol was followed, but with modifications to 3’ and 5’ adapter ligations: 3’ adapter ligation was performed at 16°C overnight and 5’ adapter ligation was performed at 20°C for two hours. Following adapter ligation, libraries were amplified for 18 cycles and were selected for a size of 108-180 base pairs using BluePippin (Sage Science). For degradome sequencing, RNA was isolated using Trizol from 20 untreated and colchicine-treated cnnos1∷GFP *Hydra*. Starting with 2 *μ*g of total RNA, poly(A) enrichment was performed to capture mRNAs, which was followed by 5’ adapter ligation. Ligation of the 5’ adaptor was done without removing the 5’ m7G cap so that only cleaved mRNAs were available for ligation. The RNA was then chemically fragmented and 3’ adapters were ligated. For ligations, 25% of the standard adapter mass was used. Libraries were prepared following the NEXTflex Small RNA-Seq Kit v3 protocol. Libraries were amplified using 14 cycles. Samples were sheared for one minute at 94°C and size selected for 125-200 base pairs using BluePippin. Degradome and piRNA libraries were pooled and sequenced using a HiSeq 4000 single end 50 bp run.

### Analysis of sequenced piRNAs and degradome reads

Adapters were trimmed from piRNA and degradome sequencing reads using cutadapt (Martin 2011). After trimming, piRNAs and degradome reads were mapped to a previously published *Hydra vulgaris* AEP transcriptome (Siebert et al. 2019) using RSEM v1.2.31 and bowtie v1.1.2 (Langmead et al. 2009). Mapping parameters were as follows: the command “rsem-calculate-expression” and the parameters “--forward-prob 0 --bowtie-n 3” was used for antisense mapping and the command “--forward-prob 1 --bowtie-n 0” was used for sense mapping. Degradome reads were mapped using the same parameters as sense piRNA reads. Boxplots were generated in R using custom code (see supplementary analysis file “1_piRNA_and_Degradome_Counts.Rmd”). Ping-pong analysis was performed using custom scripts available in the accompanying git repository. Overlap plots were generated in R using custom code (see supplementary analysis file “3_Ping_Pong_Analysis.Rmd”). Mapping results were visually inspected using Integrative Genomics Viewer (IGV) (Robinson et al. 2011).

### Transcript annotation

**Transposable Element (TE)**: TEs were annotated in our transcriptome by using the Repbase, Swissprot, PFAM, and nr databases. First, BLAST against repeat class “Transposable element” available for *Hydra vulgaris* in Repbase (Bao et al. 2015) (https://www.girinst.org/repbase/) was used to identify transcripts with TE identity (e-value cutoff of 1e-5). Next, BLAST against the Swissport database was used to add Uniprot protein descriptions to transcripts using the Uniprot Retrieve ID/mapping tool (https://www.uniprot.org/uploadlists/, cutoff e-value of 1e-5). Transcripts with Uniprot protein descriptions containing the character strings “transpos”, “jerky”, or “mobile element” were classified as TEs. Finally, previously published PFAM annotations of our *Hydra* transcriptome (Siebert et al. 2019) were used to predict additional TEs. Transcripts predicted to encode domains containing “transposase”, “THAP”, or “_tnp_” in the domain descriptor were also classified as TEs. **Gene**: Transcripts with the following characteristics were placed in the “gene” category: 1) received a Swissprot or nr annotation and 2) not classified as TE by the method described above. **Uncharacterized**: Transcripts with the following characteristics were placed in the “uncharacterized” category: 1) lack of Swissprot/nr/Pfam annotation, 2) predicted ORF of 100 amino acids or greater, and 3) not classified as a TE by the approach described above. This category contains taxonomically restricted genes. **Non-coding (ncRNA)**: Transcripts with the following characteristics were placed in the “ncRNA” category: 1) lack of Swissprot hit, nr hit, or PFAM domain, 2) ORF less than 100 amino acids, and 3) not classified as a TE by the method described above. Annotation results described above are summarized in the supplementary file “Table_S1”

### GO-term enrichment analysis

GO-term enrichment analysis was performed on the 441 transcripts upregulated in response to *hywi* RNAi-1 using GOATOOLS v0.6.10 (https://github.com/tanghaibao/goatools) (Klopfenstein et al. 2018). GO terms were considered enriched if the bonferroni-corrected p-value was <0.05.

### RNA immunoprecipitation (RIP) and RT-PCR

For each RIP experiment, 10 colchicine-treated cnnos1∷GFP *Hydra* were used to prepare a lysate and perform an immunoprecipitation as described above for piRNA isolation. Prior to immunoprecipitation, 5% of the total cleared lysate volume was removed and saved as the “input” sample. After immunoprecipitation, 1 ng of Luciferase Control RNA (Promega L4561) was added to the washed beads as a positive control for subsequent RNA extraction and cDNA synthesis. RNA was extracted from the beads using Trizol. cDNA was synthesized using M-MLV Reverse Transcriptase, RNase H Minus, Point Mutant (Promega, M3681) and random hexamers (ThermoFisher Scientific, N8080127). Three biological replicates were performed for each transcript tested (**Fig. S3**). PCR was performed using primers spanning 75-150 base pairs (**Table S5**) for each transcript using GoTaq® Green Master Mix (Promega, M7122) for 35 cycles.

### qRT-PCR

To test the effect of colchicine treatment on TE expression, RNA was extracted from 10 colchicine-treated cnnos1∷GFP *Hydra* using Trizol. To perform qPCR on *hywi* RNAi-2 hatchlings, RNA was extracted from 10 hatchlings using Trizol. For all qPCR, cDNA was synthesized using M-MLV Reverse Transcriptase, RNase H Minus, Point Mutant (Promega, M3681) and random hexamers (ThermoFisher Scientific, N8080127). qRT-PCR was performed using SsoAdvanced™ Universal SYBR® Green Supermix (BioRad,1725271) with primers spanning 75-150 base pairs for each transcript assayed (Table S5). *RP49* was used for normalization. Fold change was calculated using the ΔΔ Ct method.

### Differential Gene Expression Analysis

For differential gene expression (DGE) analysis, offspring from the *hywi* RNAi-1 transgenic line were collected 4 days post hatching. Offspring that did not inherit the transgene were considered “wild type.” Three wild type and three *hywi* RNAi replicates were collected consisting of 10 animals each and RNA was extracted using Trizol. Total RNA libraries were prepared using the Tru-Seq stranded RNA Kit (Illumina RS-122-2201). Libraries were sequenced on one lane of an Illumina HiSeq 2000 using a single-end 50 bp sequencing strategy. Illumina TruSeq3 Adapters were trimmed using trimmomatic (2:30:10 LEADING:3 TRAILING:3 SLIDINGWINDOW:4:15 MINLEN:36). RSEM v1.2.31 (Li and Dewey 2011), bowtie v1.1.2 (Langmead et al. 2009), and the transcriptome reference were used to estimate expression levels. Differential gene expression analysis was performed using edgeR v 3.20.9 (Robinson et al. 2010). After expression normalization, replicates were contrasted for variance using function plotMDS. One replicate from each treatment was identified as an outlier and excluded from downstream analysis. Analysis code is available in file “2_Differential_Gene_Expression_GO_Analysis” in the accompanying git repository.

### Small RNA and piRNA Filtering

Small RNA libraries used for sorting piRNAs by lineage were generated in our previous study (Juliano et al. 2014); NCBI BioProject PRJNA213706.

### Single Cell Data Exploration

Single-cell sequencing data for epithelial cells from homeostatic *Hydra* were interrogated for expression of putative PIWI targets (Siebert et al. 2019). URD spline objects for endoderm and ectoderm are available from Dryad: https://doi.org/10.5061/dryad.v5r6077 (file Hydra_URD_analysis_objects). These objects contain expression data for genes that are expressed in at least 1% of the ectodermal or endodermal epithelial cells. Analysis code is available in file “5_Single_Cell_Data_Exploration” in the git repository.

## Supporting information

Supplementary Analysis

## Acknowledgements

The sequencing was carried by the DNA Technologies and Expression Analysis Cores at the UC Davis Genome Center, supported by NIH Shared Instrumentation Grant 1S10OD010786-01. We thank Lutz Froenicke and Siranoosh Ashtari for their assistance and expert advice in RNA library preparation and sequencing, Matt Settles and Nikhil Joshi of the UC Davis Bioinformatics Core for their help in building code for piRNA analysis, Rob Steele and Prashanth Rangan for their critical review and editing during manuscript preparation. BBT was supported by an NIH Training Grant (5 T32 GM 7377-39). This study was supported by UC Davis start-up funds (CEJ) and by the NIH (CEJ, 1K01AG044435-01A1).

