## Supplementary Material for "PIWI-piRNA pathway-mediated transposable element repression in *Hydra* somatic stem cells"

**Teefy et al.: Supplementary material – supplementary figures (S1-S4) and tables/table legends**

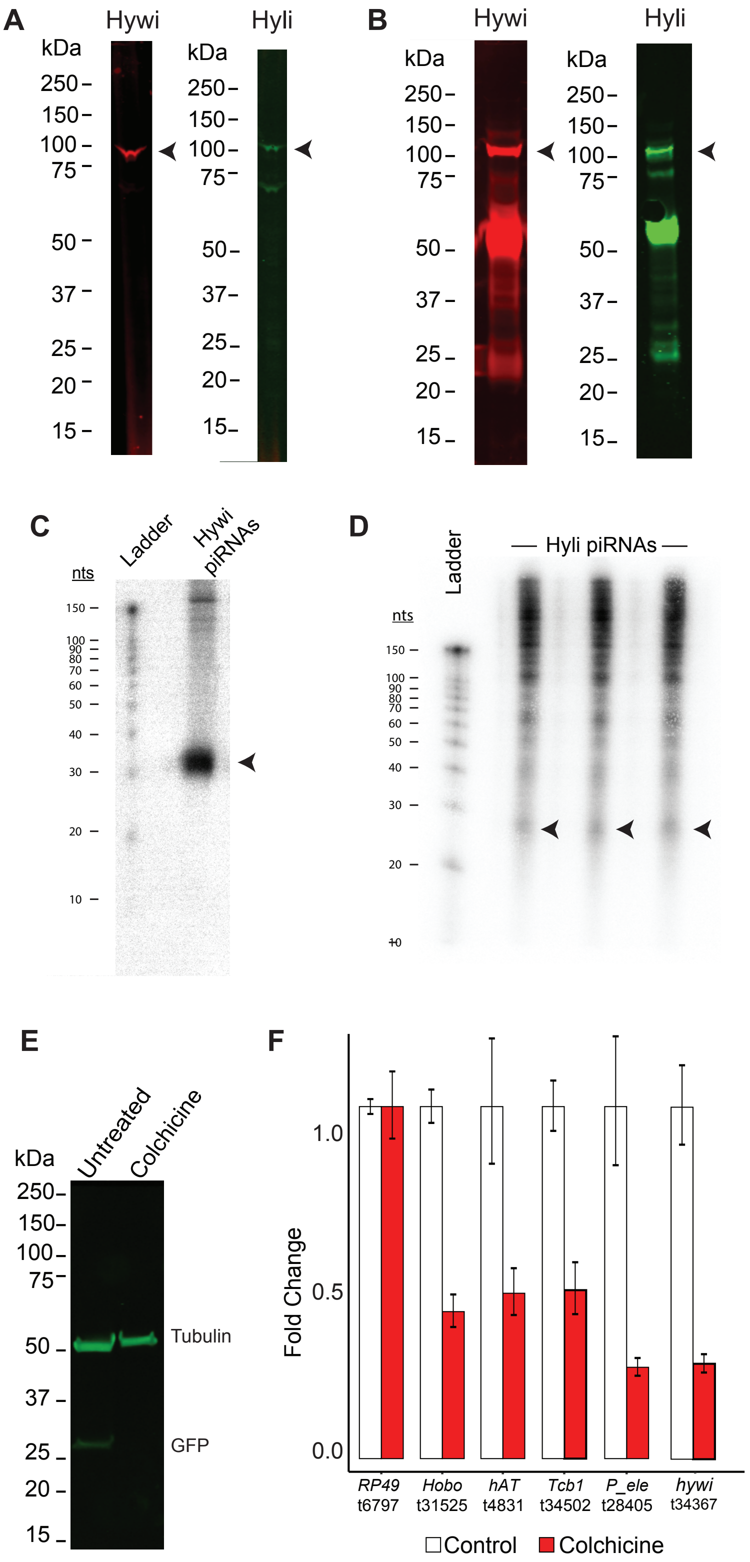

**Figure S1. Purification of PIWI-piRNA complexes from *Hydra vulgaris* AEP and colchicine treatment to remove interstitial cells. (A)** *Hydra vulgaris* strain AEP whole animal lysates were analyzed via immunoblot with Hywi and Hyli antibodies generated in a previous study against *Hydra vulgaris* 105 PIWI sequences (Juliano et al. 2014). A 100k Da band (arrowheads) is detected for both the Hywi and Hyli immunoblots demonstrating that the antibodies cross-react in the *Hydra vulgaris* AEP strain. *Hydra vulgaris* AEP is referred to as "*Hydra*" throughout the manuscript. **(B)** Hywi and Hyli immunoprecipitations were performed from whole *Hydra* lysates and Hywi and Hyli proteins were detected by immunoblot (arrowheads). Bands at ~50 kDa are the antibody heavy chain and bands at ~25 kDa are the antibody light chain. **(C,D)** piRNAs were extracted from immunoprecipitated PIWI-piRNA complexes and end-labeled with $^{32}P$, separated on a TBE-Urea gel, and visualized with a Phosphor screen. (**C**) Hywi- and (**D**) Hyli-bound piRNAs (arrowheads) run at the expected size (~30 nts for Hywi and < ~30 nts for Hyli) (Juliano et al. 2014). Hyli immunoprecipitations were run in triplicate to collect sufficient piRNAs for sequencing. **(E)** Lysates from untreated and colchicine-treated *cnnos1::GFP Hydra* (Hemmrich et al. 2012) were analyzed via immunoblot with a GFP antibody to test for the loss of GFP-positive interstitial cells. Colchicine treatment eliminated the 27 kDa GFP band, indicating the successful elimination of interstitial cells. **(F)** TE expression levels were assayed following colchicine treatment by qPCR to test if the treatment increased TE expression. Colchicine treatment appeared to decrease TE expression which is likely due to TE expression in the interstitial lineage. Similarly, we observed lower expression of *hywi* due to loss of ISCs. *RP49* is not expected to change and is used as an internal control.

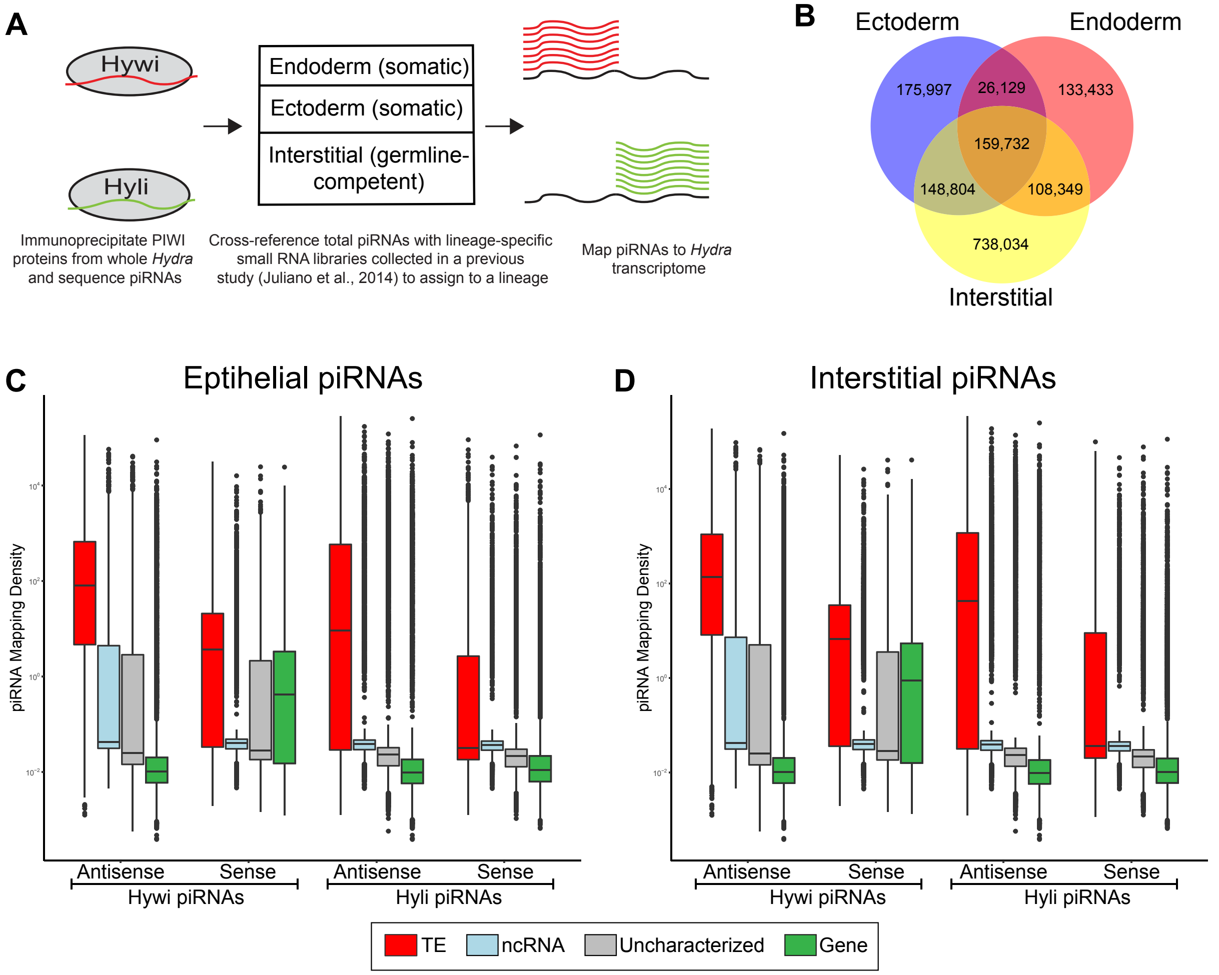


**Figure S2. piRNAs specific to each lineage map predominantly to TE transcripts (A)** Experimental design: piRNA sequences identified from whole *Hydra vulgaris* strain AEP immunoprecipitations (this s-tudy) were --cross-referenced with lineage-specific total small RNA libraries generated in our previous study; lineage-specific small RNAs were isolated by FACS of each lineage from transgenic *Hydra vulgaris* AEP (Juliano et al. 2014). **(B)** This allowed us to classify the lineage of origin for all unique piRNAs appearing in both data sets. The highest number of unique piRNAs is found in the interstitial lineage. **(C,D)** Lineage-specific piRNAs were then mapped to the *Hydra* transcriptome (Siebert et al. 2019); ectodermal- and endodermal-specific piRNAs (i.e. epithelial piRNAs) were combined for the mapping. The X-axis shows results subdivided by: 1) Hywi- or Hyli-bound piRNAs, 2) Mapped in a sense or antisense orientation, and 3) Transcript class (see legend). In all cases, TEs have a significantly higher mapping density than other transcript classes as determined using Tukey's range test ($p < 0.001$; Supplementary Analysis 4). Both interstitial and epithelial piRNAs show similar mapping patterns suggesting similar functions for the pathway in all lineages.

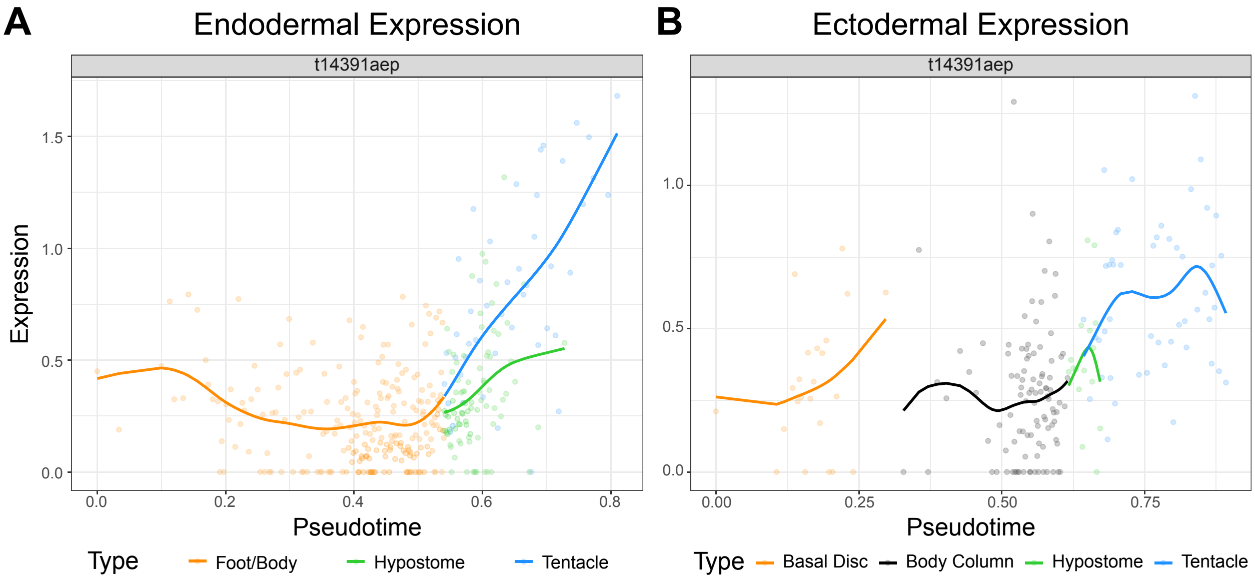


**Figure S3. Epithelial expression patterns of a putative PIWI-piRNA pathway target.** Thirty-six gene transcripts were identified as putative epithelial targets of the PIWI-piRNA pathway based on both high levels of piRNA mapping and upregulation in response to *hywi* knockdown (**Table S3**). The homeostatic epithelial expression patterns of these genes were explored using epithelial differentiation trajectories built from single cell sequencing data (Siebert et al. 2019). For 24 genes we obtained expression in at least one of the epithelial layers in homeostatic *Hydra*. We hypothesized that targets of the PIWI-piRNA would have low expression in the body column, where *hywi* is expressed, and relatively higher expression in the basal disk, hypostome, and tentacles. (**A,B**) In the differentiation trajectories, each dot represents a single cell, the X-axis shows the position of the cell along the oral-aboral axis, and the Y-axis indicates the expression level of the gene. One of the thirty-six putative target transcripts, t14391 (similarity to *PARP-12*), was found to have lower expression in the body column as compared to the extremities in both the ectoderm and endoderm. This transcript shows the expected expression pattern, which is observed in both the (A) Endoderm and (B) Ectoderm.

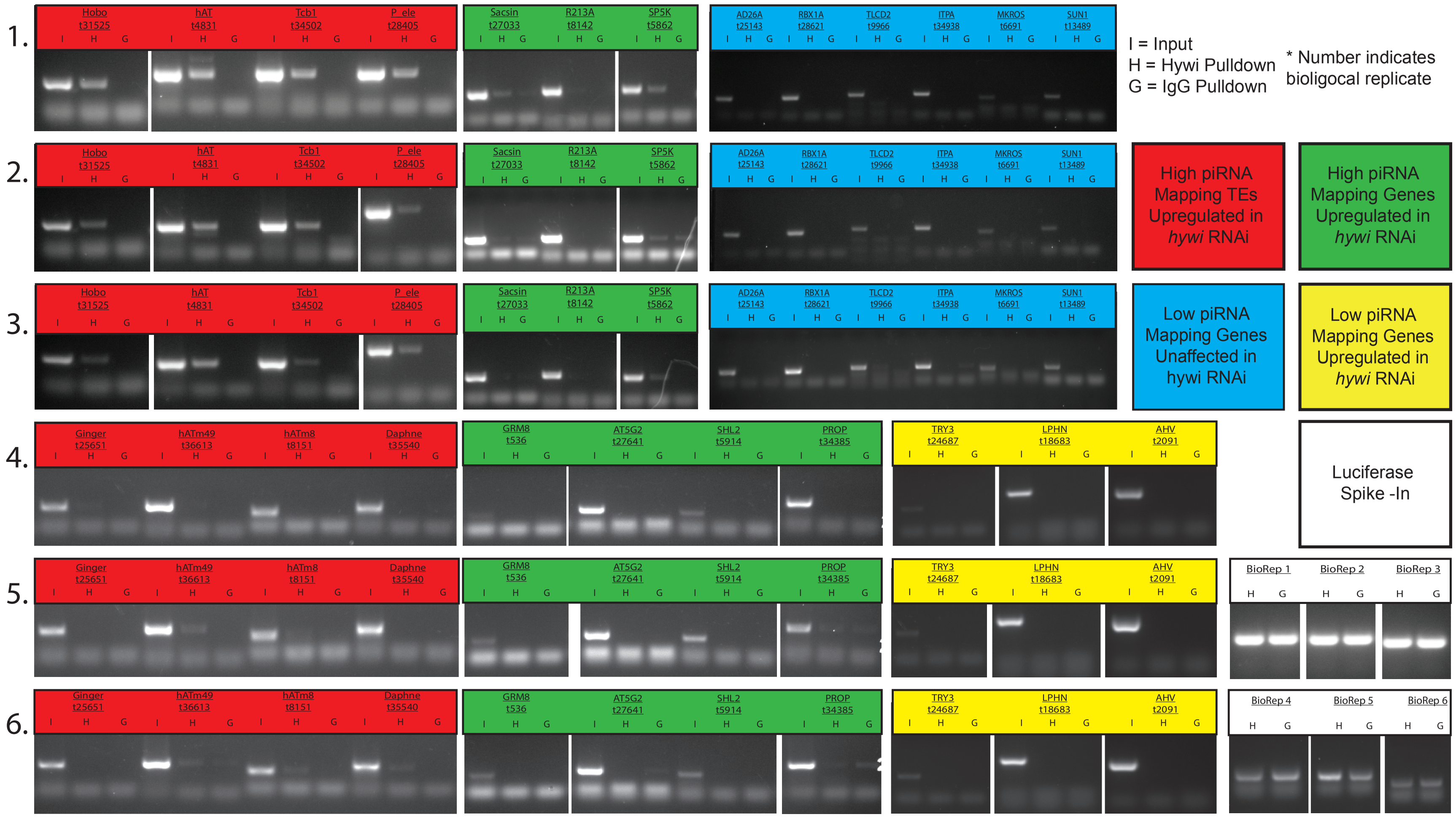
1.
2.
3.
4.
5.
6.
Hobo t31525
hAT t4831
Tcb1 t34502
P_ele t28405
Sacsin t27033
R213A t8142
SP5K t5862
AD26A t25143
RBX1A t28621
TLCD2 t9966
ITPA t34938
MKROS t6691
SUN1 t13489
Ginger t25651
hATm49 t36613
hATm8 t8151
Daphne t35540
GRM8 t536
AT5G2 t27641
SHL2 t5914
PROP t34385
TRY3 t24687
LPHN t18683
AHV t2091
BioRep 1
BioRep 2
BioRep 3
BioRep 4
BioRep 5
BioRep 6
I H G
I = Input
H = Hywi Pulldown
G = IgG Pulldown
* Number indicates bioligocal replicate
High piRNA Mapping TEs Upregulated in hywi RNAi
High piRNA Mapping Genes Upregulated in hywi RNAi
Low piRNA Mapping Genes Unaffected in hywi RNAi
Low piRNA Mapping Genes Upregulated in hywi RNAi
Luciferase Spike -In

**Figure S4. Transcripts with a high number of piRNAs mapping are more likely to immunoprecipitate with Hywi protein.** Six immunoprecipitation experiments were performed in total (numbered 1-6) to test 26 transcripts. Each transcript was assayed in triplicate. RT-PCR was used to test for the presence of target transcripts in the immunoprecipitated complexes. The following transcript categories were assayed by RT-PCR after Hywi immunoprecipitation: 1) high piRNA mapping TE transcripts upregulated in response to *hywi* knockdown (red), 2) high piRNA mapping gene transcripts upregulated in response to *hywi* knockdown (green), 3) low piRNA mapping gene transcripts upregulated in response to *hywi* knockdown (yellow), 4) low piRNA mapping gene transcripts unaffected by *hywi* knockdown (blue), and 5) Luciferase spike-in was added to the washed beads to act as a positive control for RNA extraction and cDNA synthesis. (white). The following samples were subjected to RT-PCR for each transcript assayed: Input (I), Hywi pulldown (H), and IgG pulldown (G). All TE transcripts assayed (red) amplified in at least one biological replicate and four amplified in all three replicates (see also **Fig. 4**). Out of the nine gene-encoding transcripts tested in the high mapping and responsive to knockdown category (green), two transcripts (*sacsin* and *R213A*) amplified in one biological replicate and *SP5K* amplified in two biological replicates. Low mapping gene transcripts (yellow and blue) did not amplify in any replicate. Successful amplification of the luciferase spike-in control indicated successful RNA extraction and cDNA synthesis following Hywi immunoprecipitation. For transcript IDs see **Table S5**.

**Table S1. Complete matrix of all data collected in this study – separate excel file**. Each row holds data for a single transcript in the transcriptome. See “Read Me” tab for abbreviations and explanation of columns.

**Table S2. Transcripts that are differentially expressed in response to *hywi* RNAi – separate excel file.** Bolded transcripts are classified as “high mapping” (transcripts within the top 20% of somatic Hywi piRNA density). Italicized transcripts have been classified as “high sense mapping” (transcripts within the top 5% of somatic Hywi sense-mapped piRNA density, column H). Underlined transcripts have been classified as “high antisense mapping” status (transcripts within the top 5% of somatic Hywi antisense-mapped piRNA density, column G). “P-adjusted” (column D) is the Bonferroni-corrected p-value measuring the probability of differential expression between the WT and KD groups TE: Transposable Element, ncRNA: noncoding RNA, Unchar: Uncharacterized Transcript, FC: Fold Change, S: Sense, AS: Antisense.

#### Table S3

| Transcript ID | Accession | Swissprot Hit | nr_Hit | Ping-Pong Hit | Primary Processing | Pulldown Hit Rate |
|---|---|---|---|---|---|---|
| t11116 | GHHG01011087.1 | ACT_HYDVU | PREDICTED: actin, non-muscle 6.2 [Hydra vulgaris] >gi\|113296\|sp\|P17126.1\|ACT_HYDVU RecName: Full=Actin, non-muscle 6.2 >gi\|159254\|gb\|AAA29205.1\| actin [Hydra vulgaris] | N | N | 0 |
| t14391 | GHHG01014355.1 | PAR12_HUMAN | PREDICTED: uncharacterized protein LOC101235770 [Hydra vulgaris] | N | Y | NA |
| t16179 | GHHG01016137.1 | R213A_DANRE | PREDICTED: E3 ubiquitin-protein ligase RNF213-like [Hydra vulgaris] | N | Y | NA |
| t18192 | GHHG01018145.1 | KLC4_HUMAN | PREDICTED: kinesin light chain 1-like [Hydra vulgaris] | Y | Y | NA |
| t20662 | GHHG01020603.1 | DUS7_MOUSE | PREDICTED: dual specificity protein phosphatase 6-like [Hydra vulgaris] | N | Y | NA |
| t26586 | GHHG01026513.1 | NA | hypothetical protein g.15990 [Graphocephala atropunctata] | Y | N | NA |
| t26740 | GHHG01026667.1 | RDR1_ORYSJ | PREDICTED: probable RNA-dependent RNA polymerase SHL2 [Hydra vulgaris] | N | Y | NA |
| t2683 | GHHG01002675.1 | NA | PREDICTED: ARL14 effector protein-like [Hydra vulgaris] | N | N | NA |
| t27033 | GHHG01026960.1 | SACS_HUMAN | PREDICTED: sacsin-like [Hydra vulgaris] | N | Y | 1 |
| t27094 | GHHG01027021.1 | NA | PREDICTED: uncharacterized protein LOC100210004, partial [Hydra vulgaris] | N | Y | NA |
| t27641 | GHHG01027567.1 | AT5G2_HUMAN | PREDICTED: ATP synthase F(0) complex subunit C1, mitochondrial [Larimichthys crocea] | N | N | 0 |
| t27839 | GHHG01027765.1 | RN213_HUMAN | PREDICTED: E3 ubiquitin-protein ligase RNF213-like [Hydra vulgaris] | N | Y | NA |
| t29612 | GHHG01029537.1 | NA | PREDICTED: zonadhesin-like, partial [Hydra vulgaris] | N | Y | NA |
| t29621 | GHHG01029546.1 | NA | hypothetical protein AC249_AIPGENE16468 [Exaiptasia pallida] | N | N | NA |
| t29674 | GHHG01029598.1 | R213A_DANRE | PREDICTED: E3 ubiquitin-protein ligase RNF213, partial [Hydra vulgaris] | N | Y | NA |
| t30176 | GHHG01030100.1 | R213A_DANRE | PREDICTED: E3 ubiquitin-protein ligase RNF213-like, partial [Hydra vulgaris] | N | Y | NA |
| t31117 | GHHG01031038.1 | NA | PREDICTED: reticulocyte-binding protein 2 homolog a-like [Hydra vulgaris] | N | Y | NA |
| t3141 | GHHG01003133.1 | KCP4_PINMG | PREDICTED: BPTI/Kunitz domain-containing protein 4-like [Hydra vulgaris] | N | Y | NA |
| t33022 | GHHG01032937.1 | R213A_DANRE | PREDICTED: uncharacterized protein LOC100211975 [Hydra vulgaris] | N | Y | NA |
| t33227 | GHHG01033141.1 | NA | PREDICTED: reticulocyte-binding protein 2 homolog a-like [Hydra vulgaris] | N | N | NA |
| t33764 | GHHG01033674.1 | NA | PREDICTED: zonadhesin-like, partial [Hydra vulgaris] | N | Y | NA |
| t34380 | GHHG01034289.1 | NA | PREDICTED: uncharacterized protein LOC105846005 [Hydra vulgaris] | N | Y | NA |
| t34385 | GHHG01034294.1 | PROP_MOUSE | PREDICTED: SCO-spondin-like, partial [Hydra vulgaris] | N | N | 0 |
| t34847 | GHHG01034756.1 | Y165_RICPR | PREDICTED: uncharacterized protein LOC105846304 [Hydra vulgaris] | N | Y | NA |
| t35271 | GHHG01035178.1 | NA | PREDICTED: zonadhesin-like, partial [Hydra vulgaris] | N | Y | NA |
| t35573 | GHHG01035477.1 | SHL2_ORYSJ | PREDICTED: probable ATP-dependent RNA helicase DDX58, partial [Hydra vulgaris] | N | Y | NA |

| | | | | | | |
|---|---|---|---|---|---|---|
| t37740 | GHHG01037636.1 | SAL_SILAS | PREDICTED: rhamnose-binding lectin-like [Hydra vulgaris] | N | Y | NA |
| t536 | GHHG01000533.1 | GRM8_RAT | PREDICTED: metabotropic glutamate receptor 8-like [Hydra vulgaris] | Y | Y | 0 |
| t5520 | GHHG01005511.1 | NA | PREDICTED: zonadhesin-like, partial [Hydra vulgaris] | N | Y | NA |
| t5862 | GHHG01005852.1 | SP5K_BACSU | PREDICTED: uncharacterized protein LOC101234364, partial [Hydra vulgaris] | N | Y | 2 |
| t5914 | GHHG01005904.1 | SHL2_ORYSJ | PREDICTED: probable RNA-dependent RNA polymerase 1 [Hydra vulgaris] | N | Y | NA |
| t7577 | GHHG01007562.1 | ZNFX1_MOUSE | PREDICTED: NFX1-type zinc finger-containing protein 1-like [Hydra vulgaris] | N | N | NA |
| t7958 | GHHG01007941.1 | NA | PREDICTED: uncharacterized protein LOC100212432, partial [Hydra vulgaris] | N | Y | NA |
| t7960 | GHHG01007943.1 | NA | PREDICTED: uncharacterized protein LOC100212432, partial [Hydra vulgaris] | N | N | NA |
| t8142 | GHHG01008125.1 | R213A_DANRE | PREDICTED: E3 ubiquitin-protein ligase RNF213-like [Hydra vulgaris] | N | Y | 1 |
| t8490 | GHHG01008469.1 | IF4G3_MOUSE | PREDICTED: eukaryotic translation initiation factor 4 gamma 3-like [Hydra vulgaris] | N | Y | NA |

**Table S3. Gene transcripts that are upregulated in response to *hywi* RNAi and have a high number of piRNAs mapping.** Transcripts marked as “Y” in the “Ping-Pong Hit” Column have the following mapping properties: Antisense-oriented Hywi-bound piRNAs align with a 10-nucleotide 5’ overlap with both sense-oriented Hyli-bound piRNAs and sense-oriented degradome reads; for a transcript to be marked as “Y” we require a minimum of 10 reads of each species to map in such an arrangement. Transcripts marked as “Y” in the “primary processing” column have largely sense mapping piRNAs (found within the top 5% of transcripts with respect to number of sense mapped piRNAs). For transcripts that were tested by RNA immunoprecipitation, the number of positive results out of three replicates is shown in the column labeled “pulldown hit rate”.

**Table S4. Degradome analysis of WT and colchicine-treated *Hydra* – separate excel file.** Transcripts listed as “TRUE” in the table were identified as positive “ping-pong hits” in the WT and/or colchicine-treated *Hydra*. The ping-pong signature in *Hydra* is a 10-bp overlap between the 5’ end of an antisense-oriented Hywi-bound piRNA with the 5’ end of a sense-oriented Hyli-bound piRNA and a degradome fragment. We considered transcripts that have 10 or more of each of these species mapped and aligned with this signature to be “ping-pong hits.” The last column (DE) indicates if the transcript is differentially expressed in response to *hywi* RNAi. TE: Transposable Elements, ncRNA: noncoding RNA, Unchar: Uncharacterized Transcript, Colch: Colchicine, DE: Differential Expression

### Table S5

| Transcript ID | Accession | Target Name | Forward Primer | Reverse Primer | Application | Figures |
|---|---|---|---|---|---|---|
| t6797aep | GHHG01006784.1 | RP49 | GCCAAACTGGAGAAAACCTAAAG | TGACGTGTCTTAGCATTACTTCC | qPCR | 3C;3D;S1F |
| t31525aep | GHHG01031444.1 | Hobo | TTTTCGATCTGTCGAACACG | TGCAGCTTGTAGCCGTGTAG | qPCR/RT-PCR | 3C;4D; S4;S1F |
| t4831aep | GHHG01004823.1 | hAT | TCAAGTGGCCACACTACTGC | TCGCCAATGTTGAGTTTCAG | qPCR/RT-PCR | 3C;4D; S4;S1F |
| t34502aep | GHHG01034411.1 | Tcb1 | GCCTCGACGATTCTCTTCAC | GGGTGAGCTGTCTTTGAAGC | qPCR/RT-PCR | 3C;4D; S4;S1F |
| t28405aep | GHHG01028330.1 | P_ele | CAGATGGGGCATCTACCAAC | GTGAGGTGCATCAGCAAAAA | qPCR/RT-PCR | 3C;4D; S4;S1F |
| t34367aep | GHHG01034276.1 | Hywi | CCACAACCTCCTGTTGGAGT | TGAGCAGTTTGCTGAGGTTG | qPCR | 3D |
| t5921aep | GHHG01005911.1 | CBP | GATCTGCAACCGAACCAACT | CTGCATTGCTCAATTCTCCA | qPCR | 3D |
| t16424aep | GHHG01016382.1 | MMP24 | CATGACGACCCCTACAGGTT | AGTCCTTCGTTGGTGTGAGG | qPCR | 3D |
| t13057aep | GHHG01013026.1 | RERG | CGGAAGGAACGAAGCAGTTA | AGCACAATTGGAGCTTGCTT | qPCR | 3D |
| t19259aep | GHHG01019206.1 | Casp3 | AGATGGCTCGGAAGTAGACG | CAGGATTTTACGCACCAACC | qPCR | 3D |
| t3105aep | GHHG01003097.1 | Seq1 | ATGGAAATCGCTTCAAATGC | ATGGTCTGATGGGTGCTCTC | qPCR | 3D |
| t27033aep | GHHG01026960.1 | Sacsin | TTACTCGACGCACTTTGGTG | GCACCTGCATCTTCTCCATT | RT-PCR | S4 |
| t8142aep | GHHG01008125.1 | R213A | GGTCGTCCTTGGGTTACAGA | CCAAACAATGACCAGGCTTT | RT-PCR | S4 |
| t5862aep | GHHG01005852.1 | SP5K | GTTGCACGATTACTGGCAGA | TTAGCAGATGCGGCTAGGTT | RT-PCR | S4 |
| t25143aep | GHHG01025073.1 | AD26A | CACGCGAAGAACAAGTTTCA | TTGGTGCCACAGAACTACCA | RT-PCR | S4 |
| t28621aep | GHHG01028546.1 | RBX1A | ATTGTGGGCATGGGATATTG | CACTGGTAGCTGATGCTTGG | RT-PCR | S4 |
| t9966aep | GHHG01009941.1 | TLCD2 | ATGCCTATGGATGTGGAAGC | CACAAACGTCGCATCATACC | RT-PCR | S4 |
| t34938aep | GHHG01034847.1 | ITPA | GGTTGGGAAGACAAGAGTGC | TCCCATCCAAACGAAGTAGG | RT-PCR | 4D;S4 |
| t6691aep | GHHG01006678.1 | MKROS | TGTTTTTCCCACTGCTCAGA | CAGATCTTTTTCGTGCAGCTT | RT-PCR | S4 |
| t13489aep | GHHG01013455.1 | SUN1 | ACAAATGTTGTTGGCTGCAA | TTTGGCTCCAAACCAAAAAG | RT-PCR | S4 |
| t24687aep | GHHG01024617.1 | TRY3 | TCTCCAGCAAGCAAAAATGA | TATCACCATGGCAACCAGAA | RT-PCR | 4D;S4 |
| t18683aep | GHHG01018634.1 | LPHN | CGGAAGAACCTCGTCTCGTA | GATGCTTCAACAACGCAAGA | RT-PCR | S4 |
| t2091aep | GHHG01002084.1 | AHV | AAACCGCTGTTTGGATAGCA | CTCGCGATGTGGCTTTATTT | RT-PCR | S4 |
| NA | NA | Luciferase | GCTGGGCGTTAATCAGAGAG | CGCTTCCGGATTGTTTACAT | RT-PCR | 4D;S4 |
| t25651aep | GHHG01025579.1 | Ginger | TGCACTTGCTTAATGCCTTG | ATTTCGCAATTCCACTCCTG | RT-PCR | S4 |
| t36613aep | GHHG01036512.1 | hATm49 | GTCATCTTCACGAGCACGAA | AGCAAGTTTTGAAGCGCAAT | RT-PCR | S4 |
| t8151aep | GHHG01008134.1 | hATm8 | AGCTTCATGTCCTCTTGGTGA | CGCATGGTAAACATTTTTGG | RT-PCR | S4 |
| t35540aep | GHHG01035445.1 | Daphne | GGCTCCAGGATTTGATGAGA | CATCCGGAAATATCCCAGAA | RT-PCR | S4 |
| t34385aep | GHHG01034294.1 | PROP | CGGGTACTGGTTTGTTGAGG | TTGTGCTGGGCCTAGTTCTC | RT-PCR | S4 |

| | | | | | | |
|---|---|---|---|---|---|---|
| t536aep | GHHG01000533.1 | GRM8 | CATTGGTCCGAACGAAAACT | TGCACCGTAGCTCACCATAA | RT-PCR | S4 |
| t27641aep | GHHG01027567.1 | AT5G2 | GCAGCACCAGCTCCAATAA | CTTGCAAGGACTCCAATGGT | RT-PCR | S4 |
| t5914aep | GHHG01005904.1 | SHL2 | GCTGCGTTGATAAACCCTCT | CTGCAAGATGCAGCAAAAGA | RT-PCR | S4 |

**Table S5. Primer pairs used in the study for qPCR and RT-PCR**.
