## Supplementary Analysis for "PIWI-piRNA pathway-mediated transposable element repression in *Hydra* somatic stem cells"

### 1 piRNA and Degradome Count Generation

*Bryan Teefy*

*08/09/2019*

#### Load Required Libraries

```
library(dplyr)
library(reshape2)
library(ggplot2)
library(ggpubr)
```

#### Classifying Transcripts in the *Hydra* Transcriptome

To determine the category of transcript to which piRNAs and degradome reads align, transcripts were classified as TEs, ncRNAs, uncharacterized transcripts, and genes.

The *Hydra* transcriptome was BLASTed against the *Hydra* Repbase, Swissprot, and nr databases with an e-value of 1e-5.

HMMER suite 3.1b2 (February 2015, http://www.hmmer.org/) and Pfam v31.0 database were used to identify protein domains in the transcriptome using an e-value of 1e-6.

Uniprot protein descriptions were added to any transcripts that had a match in the Swissprot database using Uniprot's Retrieve ID/mapping tool (https://www.uniprot.org/uploadlists/).

Open reading frames were identified using Transdecoder.

Results are summarized in Table S1.

#### Load Transcriptome Annotation Matrix

```
Transcript_Characterization <- read.table("objects/Transcriptome_Annotation_Matrix.txt",
    sep = "\t", check.names = FALSE, header = TRUE)
```

##### Transposon Annotation

Transcripts that met the following criteria were classified as TEs:

Transcripts with significant similarities to entries in the Repbase database.

Transcripts with Swissprot protein descriptions or nr sequence descriptions containing the strings "transpos", "J/jerky", and "mobile element".

Transcripts with Pfam domain descriptions predicted to encode domains containing "transposase", "THAP", "DDE_Tnp", "_Tnp" or "*tnp*".

#### non-coding RNA (ncRNA) Annotation

We considered sequences non-coding RNAs if they were lacking TE annotation, Swissprot hit, nr hit, known PFAM domain, and an ORF equal to or greater than 100 amino acids. ORFs were predicted using Transdecoder using command TransDecoder.LongOrfs -S -t.

#### Taxonomically Restricted Genes (TRGs)/Uncharacterized Genes

Uncharacterized Genes were defined as transcripts predicted to contain an ORF equal to or greater than 100 amino acids without a Swissprot Hit, nr hit, known domain, or TE annotation. Nr hits termed "uncharacterized protein" were also considered in this category.

#### Gene Annotation

Genes were defined as transcripts with a Swissprot hit, nr hit, or domain annotation, and that were not classified as TEs by our annotation.

```
#Classify transcripts

Transcript_Characterization$Transcript_Class <- ifelse((

  !is.na(Transcript_Characterization$Repbase_Hit) | grepl(("transpos"), Transcript_Characterization$Uni
    grepl(("mobile element"), Transcript_Characterization$Uniprot_Description, ignore.case = TRUE) |
    grepl(("jerky"), Transcript_Characterization$Uniprot_Description, ignore.case = TRUE) |
    grepl(("transpos"), Transcript_Characterization$nr_Hit, ignore.case = TRUE) |
    grepl(("mobile element"), Transcript_Characterization$nr_Hit, ignore.case = TRUE) |
    grepl(("jerky"), Transcript_Characterization$nr_Hit, ignore.case = TRUE) |
    grepl(("Transposase"), Transcript_Characterization$PFAM_Annotation, ignore.case = TRUE) |
    grepl(("THAP"), Transcript_Characterization$PFAM_Annotation, ignore.case = TRUE) |
    grepl(("_Tnp_"), Transcript_Characterization$PFAM_Annotation, ignore.case = TRUE) |
    grepl(("DDE_Tnp"), Transcript_Characterization$PFAM_Annotation, ignore.case = TRUE) |
    grepl(("_Tnp"), Transcript_Characterization$PFAM_Annotation, ignore.case = TRUE)),"TE",

    ifelse(is.na(Transcript_Characterization$ORF) & is.na(Transcript_Characterization$Uniprot_Descripti

    ifelse((!is.na(Transcript_Characterization$ORF) & is.na(Transcript_Characterization$Uniprot_Descrip

#Visualize Transcriptome Breakdown

transcriptome_annotation_whole <- c(sum(Transcript_Characterization$Transcript_Class == "TE"),sum(Trans

lbls <- c("TEs", "ncRNAs", "Unchar", "Genes")
colors = c("red", "blue", "gray", "green")
pct <- round(transcriptome_annotation_whole/sum(transcriptome_annotation_whole)*100)
lbls <- paste(lbls, pct) # add percents to labels
lbls <- paste(lbls,"%",sep="") # ad % to labels
pie(transcriptome_annotation_whole,labels = lbls, col=colors,
    main="Transcriptome Transcript Composition")
```

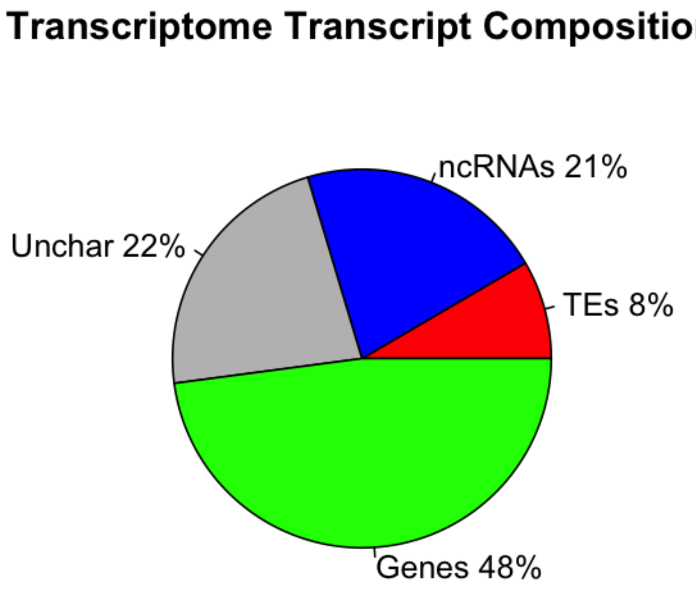


#### Generating piRNA and Degradome counts

Since piRNAs are complementary to RNA transcript targets (antisense to the target) or derived directly from targets (sense to the target), we used piRNA mapping to identify these targets in the *Hydra* transcriptome.

Adapters were trimmed from the WT and Colchicine raw reads using the script trimbiooadapter.sh.

piRNAs were mapped to the *Hydra* transcriptome using RSEM functions rsem-calculate-expression and rsem-generate-data-matrix was used to generate the count matrix. Three mismatches were allowed in the antisense orientation and no mismatches were allowed in the sense orientation. Degradome reads were mapped in the sense orientation with no mismatches since degradome reads are transcript fragments.

piRNA_Deg_rsem_mapping.sh was used for piRNA and Degradome Mapping.

Results are summarized in the table, "piRNA_Counts_Matrix.txt" and "Degradome_Counts_Matrix.txt", which can be found in the GEO repository (GSE135440).

#### Load and Merge the piRNA and Degradome Count Files

```
piRNA_counts <- read.table("objects/piRNA_Counts_Matrix.txt", sep = "\t", header = T)

Deg_counts <- read.table("objects/Degradome_Counts_Matrix.txt", sep = "\t", header = T)

piRNA_Deg_counts <- merge(piRNA_counts, Deg_counts, by = "ID")
```

```r
# Merge with Count Files

piRNA_Deg_counts <- merge(Transcript_Characterization, piRNA_Deg_counts, by = "ID")
```

#### Generate Normalized piRNA Counts

PIWI targets should have a high density of piRNA counts. We normalize piRNA counts by transcript length to determine piRNA count density.

To determine if the piRNA count density values were significantly different between classes of transcripts, we performed Tukey's Honest Significant Difference test to compare mean piRNA count density between each transcript type (i.e. TE, ncRNA, Unchar., Gene) for each piRNA class (i.e. Hywi Antisense-mapped, Hyli Sense-mapped).

```r
# Determine count density by dividing counts by transcrpt length in kilobases

norm <- (piRNA_Deg_counts$Length/1000)

piRNA_Deg_counts$WT_Hyli_AS_kb <- piRNA_Deg_counts$WT_Hyli_AS/norm
piRNA_Deg_counts$WT_Hyli_S_kb <- piRNA_Deg_counts$WT_Hyli_S/norm
piRNA_Deg_counts$WT_Hywi_AS_kb <- piRNA_Deg_counts$WT_Hywi_AS/norm
piRNA_Deg_counts$WT_Hywi_S_kb <- piRNA_Deg_counts$WT_Hywi_S/norm
piRNA_Deg_counts$WT_Deg_S_kb <- piRNA_Deg_counts$WT_Deg_S/norm
piRNA_Deg_counts$Colch_Hyli_AS_kb <- piRNA_Deg_counts$Colch_Hyli_AS/norm
piRNA_Deg_counts$Colch_Hyli_S_kb <- piRNA_Deg_counts$Colch_Hyli_S/norm
piRNA_Deg_counts$Colch_Hywi_AS_kb <- piRNA_Deg_counts$Colch_Hywi_AS/norm
piRNA_Deg_counts$Colch_Hywi_S_kb <- piRNA_Deg_counts$Colch_Hywi_S/norm
piRNA_Deg_counts$Colch_Deg_S_kb <- piRNA_Deg_counts$Colch_Deg_S/norm

# Perform Tukey's Honest Significant Difference test

# Group normalized mapping counts
Normalized_Mapping_Counts_Matrix <- piRNA_Deg_counts[, c(22:25, 27:30, 11)]
Normalized_Mapping_Counts_Matrix_Formatted <- melt(Normalized_Mapping_Counts_Matrix,
    id.var = "Transcript_Class")

# Subset count density based on piRNA origin
WT_Hyli_AS_Stats <- subset(Normalized_Mapping_Counts_Matrix_Formatted, variable ==
    "WT_Hyli_AS_kb")
WT_Hyli_S_Stats <- subset(Normalized_Mapping_Counts_Matrix_Formatted, variable ==
    "WT_Hyli_S_kb")
WT_Hywi_AS_Stats <- subset(Normalized_Mapping_Counts_Matrix_Formatted, variable ==
    "WT_Hywi_AS_kb")
WT_Hywi_S_Stats <- subset(Normalized_Mapping_Counts_Matrix_Formatted, variable ==
    "WT_Hywi_S_kb")

Colch_Hyli_AS_Stats <- subset(Normalized_Mapping_Counts_Matrix_Formatted, variable ==
    "Colch_Hyli_AS_kb")
Colch_Hyli_S_Stats <- subset(Normalized_Mapping_Counts_Matrix_Formatted, variable ==
    "Colch_Hyli_S_kb")
Colch_Hywi_AS_Stats <- subset(Normalized_Mapping_Counts_Matrix_Formatted, variable ==
    "Colch_Hywi_AS_kb")
```

```r
Colch_Hywi_S_Stats <- subset(Normalized_Mapping_Counts_Matrix_Formatted, variable ==
    "Colch_Hywi_S_kb")

# Develop Tukey Test Function (ANOVA post hoc test)

Tukey_Test <- function(x) {
    res.aov <- aov(value ~ Transcript_Class, data = x)
    return(TukeyHSD(res.aov))
}

# Run Tukey Test

Tukey_Test(WT_Hyli_AS_Stats)
```

```
##   Tukey multiple comparisons of means
##     95% family-wise confidence level
##
## Fit: aov(formula = value ~ Transcript_Class, data = x)
##
## $Transcript_Class
##                    diff         lwr        upr     p adj
## ncRNA-Gene     380.1172   127.98864   632.2457 0.0006214
## TE-Gene       2003.2537  1641.98790  2364.5194 0.0000000
## Unchar-Gene    264.9371    17.73571   512.1385 0.0300726
## TE-ncRNA      1623.1365  1229.34925  2016.9237 0.0000000
## Unchar-ncRNA  -115.1801  -407.86415   177.5040 0.7429976
## Unchar-TE    -1738.3166 -2128.96747 -1347.6657 0.0000000
```

```r
Tukey_Test(WT_Hyli_S_Stats)
```

```
##   Tukey multiple comparisons of means
##     95% family-wise confidence level
##
## Fit: aov(formula = value ~ Transcript_Class, data = x)
##
## $Transcript_Class
##                    diff         lwr        upr     p adj
## ncRNA-Gene     611.2690   293.53321   929.0049 0.0000046
## TE-Gene       2817.3087  2362.03660  3272.5808 0.0000000
## Unchar-Gene    991.6493   680.12272  1303.1759 0.0000000
## TE-ncRNA      2206.0396  1709.78354  2702.2957 0.0000000
## Unchar-ncRNA   380.3803    11.53577   749.2247 0.0402490
## Unchar-TE    -1825.6594 -2317.96301 -1333.3558 0.0000000
```

```r
Tukey_Test(WT_Hywi_AS_Stats)
```

```
##   Tukey multiple comparisons of means
##     95% family-wise confidence level
##
## Fit: aov(formula = value ~ Transcript_Class, data = x)
##
## $Transcript_Class
##                    diff         lwr        upr     p adj
## ncRNA-Gene     438.5738   287.65972   589.4878 0.0000000
## TE-Gene       2380.8399  2164.60063  2597.0791 0.0000000
```

```
## Unchar-Gene    655.0570   507.09215   803.0219 0.0000000
## TE-ncRNA      1942.2661  1706.56080  2177.9714 0.0000000
## Unchar-ncRNA   216.4832    41.29426   391.6722 0.0081737
## Unchar-TE    -1725.7829 -1959.61086 -1491.9548 0.0000000
```

```
Tukey_Test(WT_Hywi_S_Stats)
```

```
##   Tukey multiple comparisons of means
##     95% family-wise confidence level
##
## Fit: aov(formula = value ~ Transcript_Class, data = x)
##
## $Transcript_Class
##                   diff         lwr        upr     p adj
## ncRNA-Gene     25.60785  -21.999540   73.21524 0.5107346
## TE-Gene       429.48593  361.271034  497.70082 0.0000000
## Unchar-Gene    48.24260    1.565567   94.91964 0.0396104
## TE-ncRNA      403.87808  329.522416  478.23374 0.0000000
## Unchar-ncRNA   22.63475  -32.630411   77.89992 0.7186042
## Unchar-TE    -381.24332 -455.006771 -307.47987 0.0000000
```

```
Tukey_Test(Colch_Hyli_AS_Stats)
```

```
##   Tukey multiple comparisons of means
##     95% family-wise confidence level
##
## Fit: aov(formula = value ~ Transcript_Class, data = x)
##
## $Transcript_Class
##                    diff         lwr       upr     p adj
## ncRNA-Gene     215.40525    68.89178  361.9187 0.0009129
## TE-Gene       1238.53260  1028.59878 1448.4664 0.0000000
## Unchar-Gene    169.34452    25.69424  312.9948 0.0131238
## TE-ncRNA      1023.12734   794.29510 1251.9596 0.0000000
## Unchar-ncRNA   -46.06073  -216.14128  124.0198 0.8987613
## Unchar-TE    -1069.18807 -1296.19777 -842.1784 0.0000000
```

```
Tukey_Test(Colch_Hyli_S_Stats)
```

```
##   Tukey multiple comparisons of means
##     95% family-wise confidence level
##
## Fit: aov(formula = value ~ Transcript_Class, data = x)
##
## $Transcript_Class
##                   diff        lwr       upr     p adj
## ncRNA-Gene     16.62708  -86.87629  120.1305 0.9763053
## TE-Gene       762.39902  614.09282  910.7052 0.0000000
## Unchar-Gene   213.11326  111.63257  314.5939 0.0000004
## TE-ncRNA      745.77194  584.11507  907.4288 0.0000000
## Unchar-ncRNA  196.48617   76.33401  316.6383 0.0001558
## Unchar-TE    -549.28577 -709.65511 -388.9164 0.0000000
```

```
Tukey_Test(Colch_Hywi_AS_Stats)
```

```
##   Tukey multiple comparisons of means
##     95% family-wise confidence level
```

```
##
## Fit: aov(formula = value ~ Transcript_Class, data = x)
##
## $Transcript_Class
##                      diff         lwr        upr     p adj
## ncRNA-Gene      312.40992    54.84956   569.9703 0.0099067
## TE-Gene        3260.65993  2891.61109  3629.7088 0.0000000
## Unchar-Gene     357.13604   104.60898   609.6631 0.0015942
## TE-ncRNA       2948.25001  2545.97905  3350.5210 0.0000000
## Unchar-ncRNA     44.72612  -254.26350   343.7157 0.9807061
## Unchar-TE     -2903.52388 -3302.59092 -2504.4568 0.0000000
```

```r
Tukey_Test(Colch_Hywi_S_Stats)
```

```
##   Tukey multiple comparisons of means
##     95% family-wise confidence level
##
## Fit: aov(formula = value ~ Transcript_Class, data = x)
##
## $Transcript_Class
##                      diff       lwr        upr     p adj
## ncRNA-Gene     -60.003234 -161.3535   41.34708 0.4248261
## TE-Gene        323.611889  178.3907  468.83306 0.0000001
## Unchar-Gene    -54.290610 -153.6603   45.07910 0.4970390
## TE-ncRNA       383.615123  225.3210  541.90923 0.0000000
## Unchar-ncRNA     5.712624 -111.9402  123.36540 0.9993069
## Unchar-TE     -377.902499 -534.9359 -220.86913 0.0000000
```

### Visualizing piRNA Mapping

Since the range of observed count density values was large, we used a log scale to visualize piRNA count density. For boxplot visualization, we added a pseudocount to the raw piRNA counts to remove any 0 count density values that would return infinite values on a log scale. The pseudocount we chose was 0.01 since that was the lowest fractional count used in our counting strategy. We explored piRNA count density for 1) Whole animals and 2) Epithelial Animals.

```r
# Set pseudocount

pseudocount <- 0.01

# Add pseudocount to raw piRNA counts then generate piRNA count density values

boxplot_matrix <- piRNA_Deg_counts[, c(12:19, 3, 11)]
boxplot_matrix[, c(1:8)] <- boxplot_matrix[, c(1:8)] + pseudocount
boxplot_matrix[, c(1:8)] <- boxplot_matrix[, c(1:8)]/(boxplot_matrix$Length/1000)

# Plot whole animal piRNA count density values

wt_matrix_data <- boxplot_matrix[, c(10, 1:4)]
wt_matrix_data_formatted <- melt(wt_matrix_data, id.var = "Transcript_Class")
colnames(wt_matrix_data_formatted) <- c("Transcript_Class", "piRNA_Origin", "piRNA_Mapping_Density")
wt_matrix_data_formatted$Transcript_Class <- factor(wt_matrix_data_formatted$Transcript_Class,
    levels = c("TE", "ncRNA", "Unchar", "Gene"))
```

```
WT_level_order <- c("WT_Hywi_AS", "WT_Hywi_S", "WT_Hyli_AS", "WT_Hyli_S")

WT_boxplot <- ggplot(data = wt_matrix_data_formatted, aes(x = factor(piRNA_Origin,
    level = WT_level_order), y = piRNA_Mapping_Density), log = "y") + geom_boxplot(aes(fill = Transcript
    scale_y_log10(breaks = scales::trans_breaks("log10", function(x) 10^x), labels = scales::trans_forma
        scales::math_format(10^.x)))

WT_boxplot + scale_fill_manual(values = c("red", "light blue", "grey", "green")) +
    theme(panel.grid.major = element_blank(), panel.grid.minor = element_blank(),
        panel.background = element_blank(), axis.line = element_line(colour = "black")) +
    theme(legend.text = element_text(size = rel(1))) + ggtitle("Whole Animal piRNA Mapping Density") +
    theme(plot.title = element_text(hjust = 0.5)) + xlab("piRNA Origin") + ylab("piRNA Mapping Density"
```

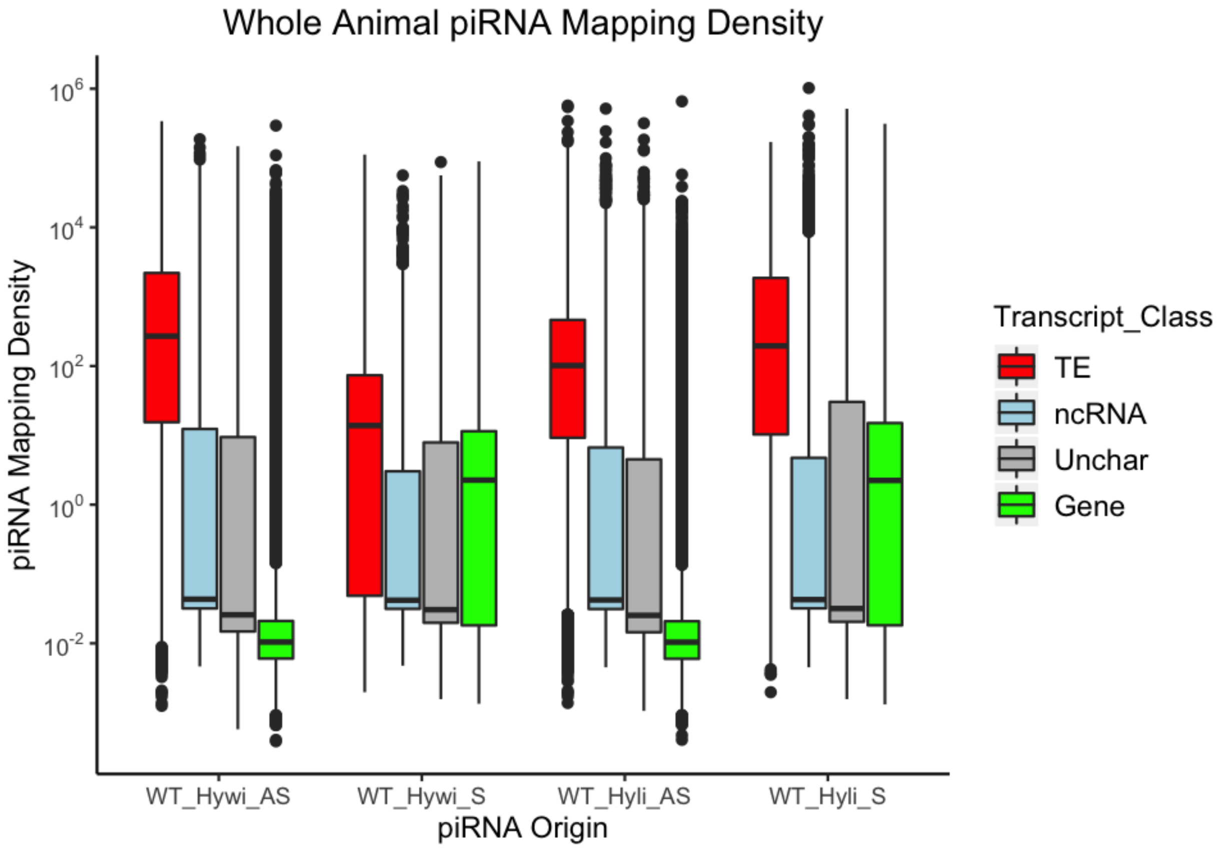


```
# Plot epithelial animal piRNA count density values

colch_matrix_data <- boxplot_matrix[, c(10, 5:8)]
colch_matrix_data_formatted <- melt(colch_matrix_data, id.var = "Transcript_Class")
colnames(colch_matrix_data_formatted) <- c("Transcript_Class", "piRNA_Origin", "piRNA_Mapping_Density")
colch_matrix_data_formatted$Transcript_Class <- factor(colch_matrix_data_formatted$Transcript_Class,
    levels = c("TE", "ncRNA", "Unchar", "Gene"))

colch_level_order <- c("Colch_Hywi_AS", "Colch_Hywi_S", "Colch_Hyli_AS", "Colch_Hyli_S")

colch_boxplot <- ggplot(data = colch_matrix_data_formatted, aes(x = factor(piRNA_Origin,
    level = colch_level_order), y = piRNA_Mapping_Density), log = "y") + geom_boxplot(aes(fill = Transc
    scale_y_log10(breaks = scales::trans_breaks("log10", function(x) 10^x), labels = scales::trans_forma
```

```
        scales::math_format(10^.x)))

colch_boxplot + scale_fill_manual(values = c("red", "light blue", "grey", "green")) +
    theme(panel.grid.major = element_blank(), panel.grid.minor = element_blank(),
        panel.background = element_blank(), axis.line = element_line(colour = "black")) +
    theme(legend.text = element_text(size = rel(1))) + ggtitle("Somatic piRNA Mapping Density") +
    theme(plot.title = element_text(hjust = 0.5)) + xlab("piRNA Origin") + ylab("piRNA Mapping Density")
```

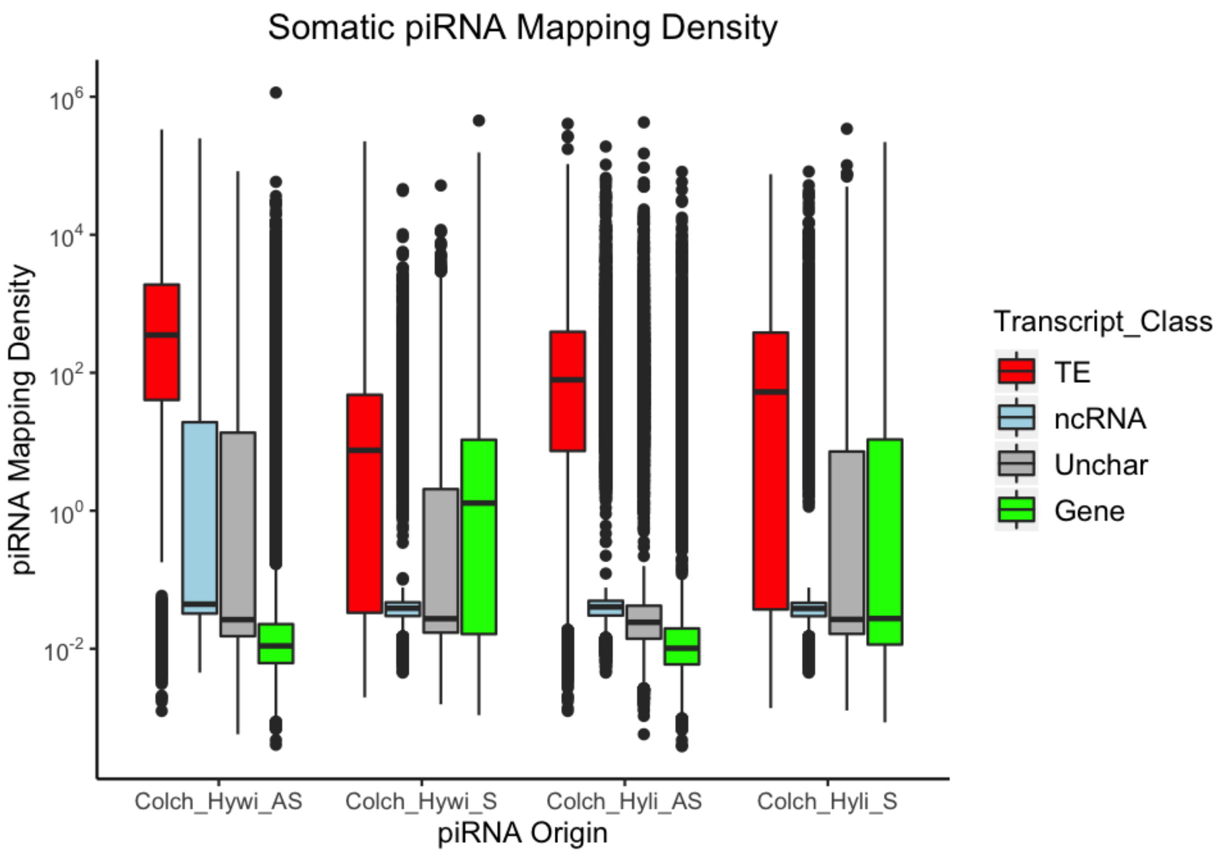


```
# write table that summarizes results write.table(piRNA_Deg_counts, file =
# 'Annotated_piRNA_Degradome_Count_Matrix.txt')
```

##### Software versions

This document was computed on Fri Aug 09 19:21:20 2019 with the following R package versions.

```
R version 3.5.3 (2019-03-11)
Platform: x86_64-apple-darwin15.6.0 (64-bit)
Running under: macOS Mojave 10.14.5

Matrix products: default
BLAS: /Library/Frameworks/R.framework/Versions/3.5/Resources/lib/libRblas.0.dylib
LAPACK: /Library/Frameworks/R.framework/Versions/3.5/Resources/lib/libRlapack.dylib

locale:
[1] en_US.UTF-8/en_US.UTF-8/en_US.UTF-8/C/en_US.UTF-8/en_US.UTF-8
```

```
attached base packages:
[1] stats     graphics  grDevices utils     datasets  methods   base

other attached packages:
[1] ggpubr_0.2     magrittr_1.5   ggplot2_3.2.0  reshape2_1.4.3
[5] dplyr_0.8.3    knitr_1.22

loaded via a namespace (and not attached):
 [1] Rcpp_1.0.1       munsell_0.5.0    tidyselect_0.2.5 colorspace_1.4-1
 [5] R6_2.4.0         rlang_0.4.0      stringr_1.4.0    plyr_1.8.4
 [9] tools_3.5.3      grid_3.5.3       gtable_0.3.0     xfun_0.5
[13] withr_2.1.2      htmltools_0.3.6  lazyeval_0.2.2   yaml_2.2.0
[17] assertthat_0.2.1 digest_0.6.20    tibble_2.1.3     crayon_1.3.4
[21] purrr_0.3.2      formatR_1.7      glue_1.3.1       evaluate_0.13
[25] rmarkdown_1.12   stringi_1.4.3    compiler_3.5.3   pillar_1.4.2
[29] scales_1.0.0     pkgconfig_2.0.2
```

### 2 Differential Gene Expression Analysis and GO Enrichment

*Stefan Siebert and Bryan Teefy*

*08/09/2019*

##### Differential Gene Expression

We performed differential gene expression analysis between wildtype and *hywi* knockdown animals. As a first step we generated expression estimates using RSEM/bowtie and used reads that were filtered for adapters (TruSeq3) using trimmomatic (LEADING:3 TRAILING:3 SLIDINGWINDOW:4:15 MINLEN:36). We made use of the RSEM functions rsem-calculate-expression (–forward-prob 0) and rsem-generate-data-matrix to generate an expression matrix (scripts: DGE_expression.sh, DGE_count_matrix.sh). The raw count matrix is available at GEO (GSE135440).

##### Load the expression data / GO annotations

```
# Load the Differential Gene Expression Count Matrix

counts <- read.table("objects/Differential_Gene_Expression_Count_Matrix.txt", sep = "\t",
    check.names = FALSE, header = TRUE, row.names = 1)

# Load normalized piRNA and degradome reads and transcript characterization from
# RMD1

piRNA_Deg_counts <- read.table("objects/Annotated_piRNA_Degradome_Count_Matrix.txt",
    sep = "\t", check.names = FALSE, header = TRUE)
```

##### Load Required Packages

```
library(edgeR)
library(knitr)
library(xtable)
library(ggplot2)
```

#### Data exploration

We explore the data to get insights into the paired nature of the triplicate samples from wildtype and *hywi* knockdown tissue.

```
# We first calculate normalized counts for future use and to include them in the
# master dataframe.

# set gene IDs as rownames
rownames(counts) <- counts[, 1]
```

```r
# raw counts from all treatments
k <- counts[, c(2:7)]

# calculate normalization factors
nf_k <- calcNormFactors(k)

# calculate library sizes
ls_k <- colSums(k)

# effective library size using normalization factors
lse_k <- ls_k * nf_k

# normalization multiplier to use on counts
nm_k <- 1e+06/lse_k

# normalize counts using normalization multiplier
k <- k * nm_k

# round normalized counts
k <- round(k, digits = 0)

# restore ID column
k$ID <- rownames(k)

# reorder columns
k <- k[, c(7, 1:6)]

# combine rounded raw and normalized counts
k <- merge(counts[, c(1:7)], k, by = "ID")

# rename columns

colnames(k) <- c("ID", "WT1", "WT2", "WT3", "KD1", "KD2", "KD3", "nWT1", "nWT2",
    "nWT3", "nKD1", "nKD2", "nKD3")
```

```r
# explore replication

# define treatment groups for DGE
TR_k <- factor(c("k", "k", "k", "w", "w", "w"))

# set wild type as reference
TR_k <- relevel(TR_k, ref = "w")

# create experimental design data frame defining treatment type for each sample
d_k <- data.frame(Sample = colnames(counts[, c(5, 6, 7, 2, 3, 4)]), TR_k)

# generate DGEList object containing raw counts, the treatment type for each
# column, and the gene IDs
y <- DGEList(counts = counts[, c(5, 6, 7, 2, 3, 4)], group = d_k$TR_k, genes = counts[,
    1])

# label each column with the appropriate sample name
colnames(y) <- d_k$Sample
```

```r
# calculate library size for each sample and store within DGElist
y$samples$lib.size <- colSums(y$counts)

# exclude transcripts that do not have at least two samples with more than one
# count per million
keep <- rowSums(cpm(y) > 1) >= 2
y <- y[keep, , keep.lib.sizes = FALSE]

# number of transcripts remaining after count filtering
dim(y)
```

```
## [1] 16435     6
```

```r
# calculate normalization factors for each sample
y <- calcNormFactors(y)

# review library sizes and normalization factors
y$samples
```

```
##     group lib.size norm.factors
## KD1     k 27236412    1.0711124
## KD2     k 26955915    0.9927897
## KD3     k 27081381    1.0126300
## WT1     w 22165507    1.0181741
## WT2     w 27471090    0.9836625
## WT3     w 25597005    0.9272326
```

```r
# create a matrix defining the control and treatment groups for the analysis
# based on what is described by the TR object
design_k <- model.matrix(~TR_k)

# Estimate dispersions
y <- estimateDisp(y, design_k)

# generate MDS Plot

plotMDS(y, method = "bcv", cex = 0.7)
```

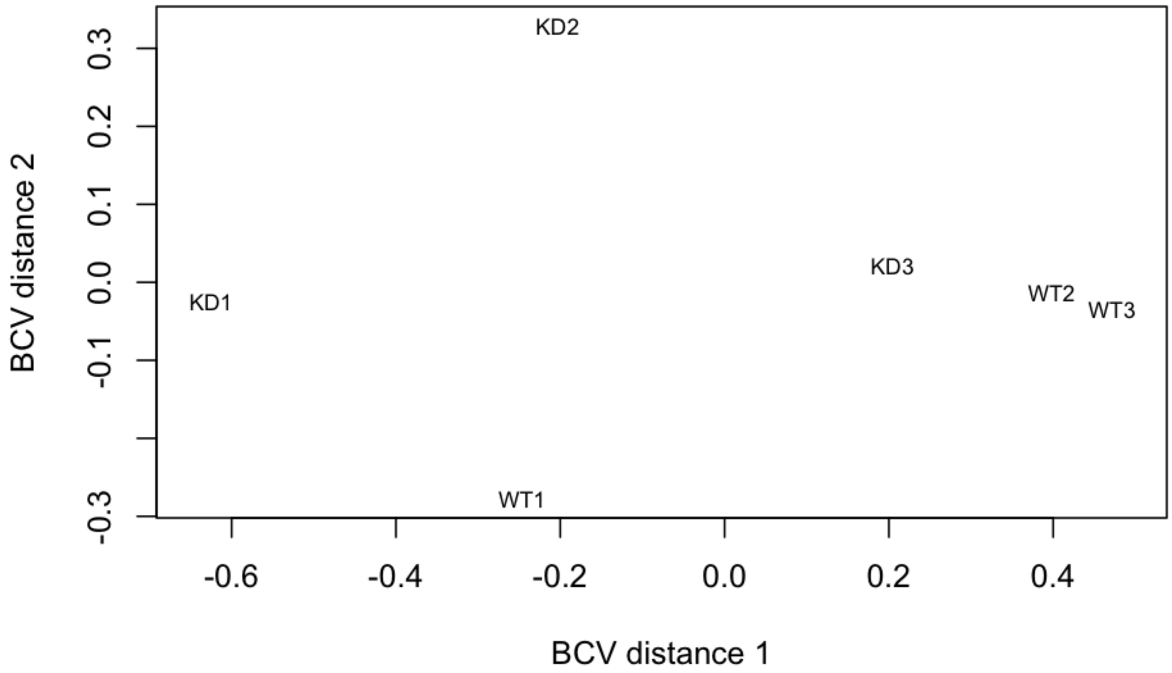


```r
# we exclude outlier replicates WT1 and KD3 from downstream analyses

# define treatment groups for DGE
TR_k <- factor(c("k", "k", "w", "w"))

# set wild type as reference
TR_k <- relevel(TR_k, ref = "w")

# create experimental design data frame defining treatment type for each sample
d_k <- data.frame(Sample = colnames(counts[, c(5, 6, 3, 4)]), TR_k)

# generate DGElist object containing raw counts, the treatment type for each
# column, and the gene annotations
y <- DGEList(counts = counts[, c(5, 6, 3, 4)], group = d_k$TR_k, genes = counts[,
    1])

# label each column with the appropriate sample name
colnames(y) <- d_k$Sample

# calculate library size for each sample and store within DGElist
y$samples$lib.size <- colSums(y$counts)

# exclude transcripts that do not have at least two samples with more than one
# count per million
keep <- rowSums(cpm(y) > 1) >= 2
y <- y[keep, , keep.lib.sizes = FALSE]
```

```r
# number of transcripts remaining after count filtering
dim(y)
```

```
## [1] 15924     4
```

```r
# calculate normalization factors for each sample
y <- calcNormFactors(y)

# review library sizes and normalization factors
y$samples
```

```
##     group lib.size norm.factors
## KD1     k 27217655    1.0849445
## KD2     k 26945252    0.9960237
## WT2     w 27460768    0.9922666
## WT3     w 25588479    0.9325978
```

```r
# create a matrix defining the control and treatment groups for the analysis
# based on what is described by the TR object
design_k <- model.matrix(~TR_k)

# estimate dispersions
y <- estimateDisp(y, design_k)

# generate MDS Plot
plotMDS(y, method = "bcv", cex = 0.7)
```

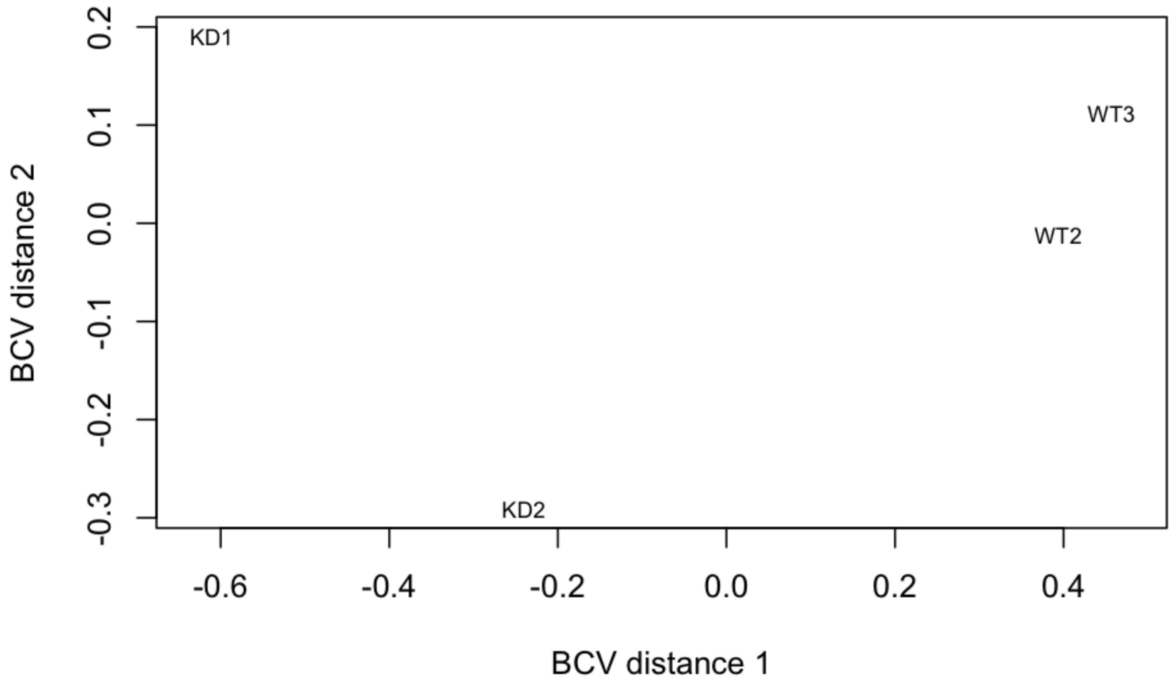

#### Differential Gene Expression (DGE) analysis - wildtype vs *hywi* knockdown.

We perform DGE analysis after excluding outlier replicates WT1 and KD3.

```
# DGE Analysis

# set p-value for cutoff
p.value = 0.05

# fit data to a negative binomial geralized log-linear model
fit_k <- glmFit(y, design_k)

# conduct a statistical test for differential gene expression based on the fit
# from
lrt_k <- glmLRT(fit_k)

# create a list of values that describes the status of each gene in the DGE test
de_k <- decideTestsDGE(lrt_k, adjust.method = "BH", p.value)

# overview of number of differentially expressed genes
summary(de_k)
```

```
##        TR_kk
## Down      17
## NotSig 15466
## Up       441
```

```
# build results table
D_k <- lrt_k$table

# restore ID column
D_k$ID <- rownames(D_k)

# add column of adjusted p values
D_k <- cbind(D_k, p.adjust(D_k$PValue, method = "BH"))
names(D_k)[names(D_k) == "p.adjust(D_k$PValue, method = \"BH\")"] = "k_Padj"

# create table summarizing rounded raw counts, normalized counts and DGE results
res_k <- merge(k, D_k, by = "ID", all = TRUE)

# call transcripts upregulated, downregulated, or unaffected

res_k$DE <- ifelse(res_k$k_Padj <= 0.05 & res_k$logFC >= 0 & !is.na(res_k$logFC &
    res_k$k_Padj), "Up", ifelse(res_k$k_Padj <= 0.05 & res_k$logFC <= 0 & !is.na(res_k$logFC &
    res_k$k_Padj), "Down", "None"))

# merge with the master dataframe

piRNA_Deg_counts <- merge(piRNA_Deg_counts, res_k, by = "ID")

# generate volcano plot

detags <- rownames(y)[as.logical(de_k)]
```

```r
plotSmear(lrt_k, de.tags = detags, cex = 0.2, cex.lab = 1, cex.axis = 1)
```

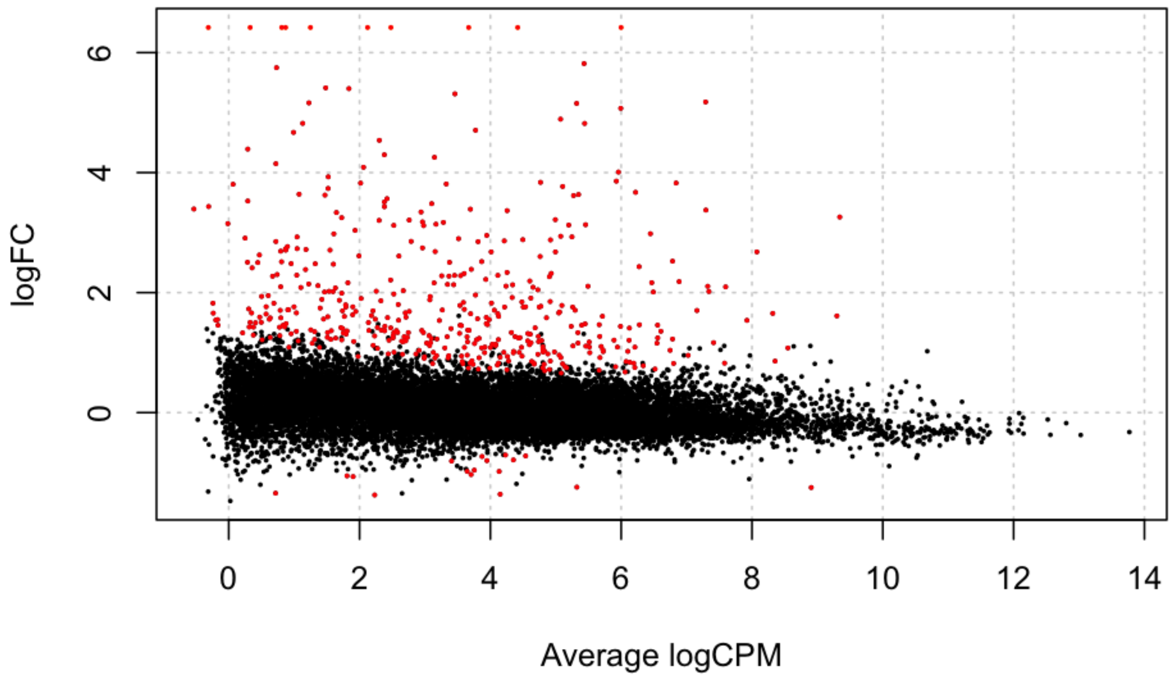


```r
# generate plot of upregulated transcripts expression fold change

Upreg_Types <- c(sum(piRNA_Deg_counts$DE == "Up" & piRNA_Deg_counts$Transcript_Class ==
    "TE"), sum(piRNA_Deg_counts$DE == "Up" & piRNA_Deg_counts$Transcript_Class ==
    "ncRNA"), sum(piRNA_Deg_counts$DE == "Up" & piRNA_Deg_counts$Transcript_Class ==
    "Unchar"), sum(piRNA_Deg_counts$DE == "Up" & piRNA_Deg_counts$Transcript_Class ==
    "Gene"))

lbls <- c("TEs", "ncRNAs", "Unchar", "Genes")
colors = c("red", "blue", "gray", "green")
pct <- round(Upreg_Types/sum(Upreg_Types) * 100)
lbls <- paste(lbls, pct)  # add percents to labels
lbls <- paste(lbls, "%", sep = "")  # ad % to labels
pie(Upreg_Types, labels = lbls, col = colors, main = "hywi RNAi Upregulated Transcript Composition")
```

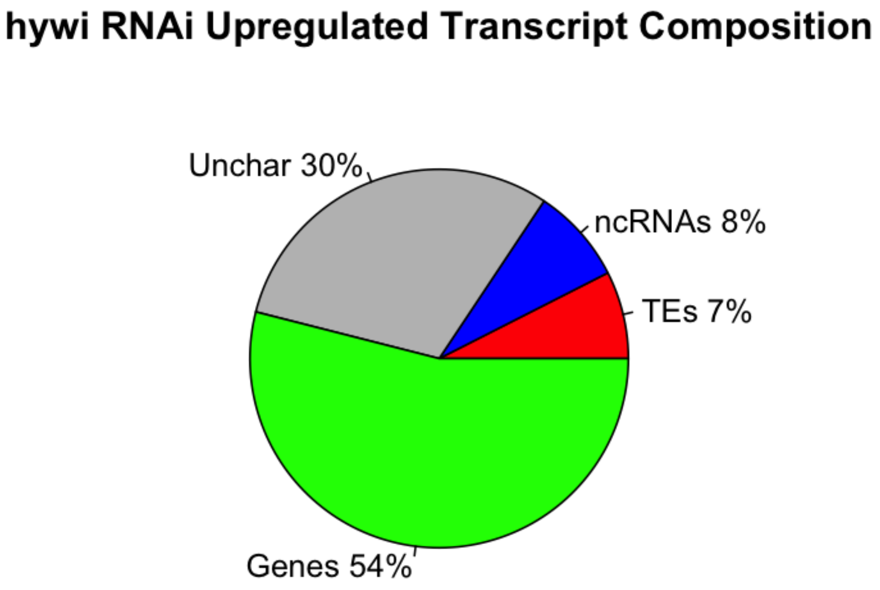


##Somatic Hywi piRNA Mapping Ordering

To infer direct targets of Hywi, *hywi* RNAi upregulated transcripts can be described as high and low Hywi piRNA mapping transcripts. "High-mapping"" transcripts are likely to be direct Hywi targets while "low-mapping" transcripts are unlikely to be direct Hywi targets. High Mapping transcripts were defined as those transcripts that were in the top 20% of read counts per kilobase million after combining Colch Hywi Sense and Colch Hywi Antisense reads.

To infer which transcripts were most likely to be involved in the ping-pong cycle in somatic stem cells, transcripts were ordered by Colch Hywi Antisense reads per kilobase million. Transcripts that fall within the top 5% in this category were considered to be putative ping-pong transcripts.

To infer which transcripts may be "primary-processed" by Hywi in somatic stem cells, transcripts were ordered by Colch Hywi Sense reads per kilobase million. Transcripts that fall within the top 5% in this category were considered to be putative primary-processed transcripts.

```
# generate RPM (Read counts Per kilobase Million) data for Colch Hywi
# Sense/Antisense piRNAs (i.e. summed Colch Hywi Sense/Antisense piRNA read
# counts mapped per kb of transcript per million piRNA read counts)

piRNA_Deg_counts$Somatic_Hywi_Perc <- (piRNA_Deg_counts$Colch_Hywi_AS_kb + piRNA_Deg_counts$Colch_Hywi_[illegible]
    piRNA_Deg_counts$Colch_Hywi_S_kb)/1e+06)

piRNA_Deg_counts$Somatic_Hywi_Perc <- ecdf(piRNA_Deg_counts$Somatic_Hywi_Perc)(piRNA_Deg_counts$Somatic_[illegible]

# repeat for only Colch Hywi Antisense piRNAs

piRNA_Deg_counts$Somatic_Hywi_Perc_AS <- (piRNA_Deg_counts$Colch_Hywi_AS_kb)/(sum(piRNA_Deg_counts$Colc[illegible]
    piRNA_Deg_counts$Colch_Hywi_S_kb)/1e+06)
```

```
piRNA_Deg_counts$Somatic_Hywi_Perc_AS <- ecdf(piRNA_Deg_counts$Somatic_Hywi_Perc_AS)(piRNA_Deg_counts$So

# repeat for only Colch Hywi Sense piRNAs

piRNA_Deg_counts$Somatic_Hywi_Perc_S <- (piRNA_Deg_counts$Colch_Hywi_S_kb)/(sum(piRNA_Deg_counts$Colch_H
    piRNA_Deg_counts$Colch_Hywi_S_kb)/1e+06)

piRNA_Deg_counts$Somatic_Hywi_Perc_S <- ecdf(piRNA_Deg_counts$Somatic_Hywi_Perc_S)(piRNA_Deg_counts$Soma

# write table that summarizes data write.table(piRNA_Deg_counts, file =
# 'Annotated_piRNA_Degradome_DGE_Count_Matrix.txt')
```

#### GO-Term Enrichment Analysis

GO-term enrichment analysis was performed on upregulated transcripts against the entire transcriptome to investigate a functional response to somatic *hywi* knockdown.

GO-term enrichment analysis was performed using goatools v0.6.10 using the script: goatools_GO_enrichment.pl

```
# load GO annotation results

GO_table <- read.table("objects/GO_upreg_trans_full_ref.txt", header = T, sep = "\t")

# take enriched biological processes

GO_table_sub <- subset(GO_table, p_bonferroni <= 0.05 & NS == "BP")

# include GO accession number, GO term, ratio in study, ratio in population,
# bonferroni corrected p-value, and transcript IDs

GO_table_sub <- GO_table_sub[, c(1, 4:6, 10, 14)]
colnames(GO_table_sub) <- c("GO", "GO_Term", "Study", "Pop", "p_val", "ID")

# print table

kable(GO_table_sub, format = "markdown", padding = 100)
```

| GO | GO_Term | Study | Pop | p_val | ID |
|---|---|---|---|---|---|
| . . . GO: 0006952 | defense response | 26/441 | 753/38747 | 0.0142 | t11117aep, t12198aep, t16424aep, t17178aep, t17750aep, t21013aep, t21682aep, t22133aep, t24687aep, t32280aep, t33020aep, t34385aep, t34424aep, t34475aep, t35573aep, t35608aep, t35837aep, t38672aep, t38673aep, t5914aep, t6387aep, t7326aep, t7388aep, t8582aep, t8645aep, t8946aep |
| . . . .GO: 0051899 | membrane depolarization | 7/441 | 46/38747 | 0.0170 | t17803aep, t18338aep, t24938aep, t25020aep, t526aep, t527aep, t9936aep |

| GO | GO_Term | Study | Pop | p_val | ID |
|---|---|---|---|---|---|
| .......GO: 2000051 | negative regulation of non-canonical Wnt signaling pathway | 4/441 | 8/38747 | 0.0221 | t29674aep, t30176aep, t33022aep, t8142aep |
| ....GO: 0048440 | carpel development | 3/441 | 3/38747 | 0.0290 | t35573aep, t38672aep, t38673aep |
| ...GO: 0045087 | innate immune response | 14/441 | 258/38747 | 0.0370 | t11117aep, t17178aep, t22133aep, t24687aep, t34385aep, t34424aep, t35573aep, t35608aep, t35837aep, t38672aep, t38673aep, t6387aep, t7326aep, t8946aep |
| .......GO: 0031050 | dsRNA processing | 5/441 | 19/38747 | 0.0377 | t30770aep, t35488aep, t38672aep, t38673aep, t5914aep |
| ......GO: 0042108 | positive regulation of cytokine biosynthetic process | 5/441 | 20/38747 | 0.0498 | t21682aep, t22133aep, t24687aep, t34424aep, t8645aep |

**Software versions**

This document was computed on Fri Aug 09 19:21:50 2019 with the following R package versions.

```
R version 3.5.3 (2019-03-11)
Platform: x86_64-apple-darwin15.6.0 (64-bit)
Running under: macOS Mojave 10.14.5

Matrix products: default
BLAS: /Library/Frameworks/R.framework/Versions/3.5/Resources/lib/libRblas.0.dylib
LAPACK: /Library/Frameworks/R.framework/Versions/3.5/Resources/lib/libRlapack.dylib

locale:
[1] en_US.UTF-8/en_US.UTF-8/en_US.UTF-8/C/en_US.UTF-8/en_US.UTF-8

attached base packages:
[1] stats     graphics  grDevices utils     datasets  methods   base

other attached packages:
[1] ggplot2_3.2.0 xtable_1.8-3  edgeR_3.22.5  limma_3.38.3  knitr_1.22

loaded via a namespace (and not attached):
 [1] Rcpp_1.0.1        magrittr_1.5      splines_3.5.3     tidyselect_0.2.5
 [5] munsell_0.5.0     colorspace_1.4-1 lattice_0.20-38  R6_2.4.0
 [9] rlang_0.4.0       highr_0.7         dplyr_0.8.3       stringr_1.4.0
[13] tools_3.5.3       grid_3.5.3        gtable_0.3.0      xfun_0.5
[17] withr_2.1.2       htmltools_0.3.6   assertthat_0.2.1 yaml_2.2.0
[21] lazyeval_0.2.2    digest_0.6.20     tibble_2.1.3      crayon_1.3.4
[25] purrr_0.3.2       formatR_1.7       glue_1.3.1        evaluate_0.13
[29] rmarkdown_1.12    stringi_1.4.3     compiler_3.5.3    pillar_1.4.2
[33] scales_1.0.0      locfit_1.5-9.1    pkgconfig_2.0.2
```

### 3 Ping Pong Analysis

*Bryan Teefy*

*08/09/2019*

#### Ping-Pong Hits

PIWI targets are often degraded with a distinctive "ping-pong signature" consisting of a 10 bp overlap between the 5' ends of antisense-mapped piRNAs and sense-mapped piRNAs/degradome reads. Transcripts that had a ping-pong signature comprised of at least 10 of the contributing species (antisense piRNA, sense piRNA and degradome read) in the correct orientation were deemed "ping-pong hits".

To identify transcripts with "ping-pong" hits, we used the following approach:

piRNA and Degradome BAM files generated from the RSEM mapping strategy in RMD 1 were converted to BED files using the script: rsem_bam2bed.sh.

The script rsem_grouping.sh retains only those transcripts that have at least 10 piRNA/degradome reads beginning at a particular position such that the depth criteria for calling a ping-pong hit can be met. The script outputs transcript ID, start position of a piRNA/degradome read, and 10 bps from the start of the piRNA/degradome read. Adding 10 bps from the start position allows for the subsequent overlap_BT.perl script to find reads that overlap by the specified 10 bp length.
rsem_grouping.sh uses: "group_reads_sense.perl" and "group_reads_antisense.perl"

The script overlap_rsem.sh generates a matrix containing ping-pong hit coordinates.
overlap_rsem.sh makes uses: "overlap_BT.perl"

overlap_rsem.sh output files were converted to .txt files, merged in R, and used to generate TRUE/FALSE statements in reference to whether a transcript has a Whole Animal or a Epithelial Animal ping-pong hit.

The resultant table is: "rsem_ping_pong_hits.txt"

#### Load the Necessary Files

```
#Load normalized piRNA and degradome counts, expression data, DGE data from RMD2.

Read_Counts_Master_DF <- read.table("objects/Annotated_piRNA_Degradome_DGE_Count_Matrix.txt", sep = "\t

#Load binary matrix of ping-pong hits per transcript.

Ping_Pong_Matrix <- read.table("objects/rsem_ping_pong_hits.txt", sep = "\t")
```

#### Load Necessary Packages

```
###load packages###

library(ggplot2)

library(reshape2)
```

### Visualizing Ping-Pong Hits

To visualize ping-pong hit distribution, we generated pie charts consisting of all transcripts that have at least one ping-pong hit and plot the output by transcript class for 1) Whole Animals and 2) Epithelial Animals.

```
Read_Counts_Master_DF <- merge(Read_Counts_Master_DF, Ping_Pong_Matrix, by = "ID")

#retrieve hits

WT_ping_hits <- c(sum(Read_Counts_Master_DF$Whole_Ping_Pong & Read_Counts_Master_DF$Transcript_Class ==

Colch_ping_hits <- c(sum(Read_Counts_Master_DF$Epi_Ping_Pong & Read_Counts_Master_DF$Transcript_Class =

#create pie chart with percentages

#Whole Animal
lbls <- c("TEs", "ncRNAs", "Unchar", "Genes")
colors = c("red", "blue", "gray", "green")
pct <- round(WT_ping_hits/sum(WT_ping_hits)*100)
lbls <- paste(lbls, pct) # add percents to labels
lbls <- paste(lbls,"%",sep="") # ad % to labels
pie(WT_ping_hits,labels = lbls, col=colors,
    main="Whole Animal Ping-Pong Hits")
```

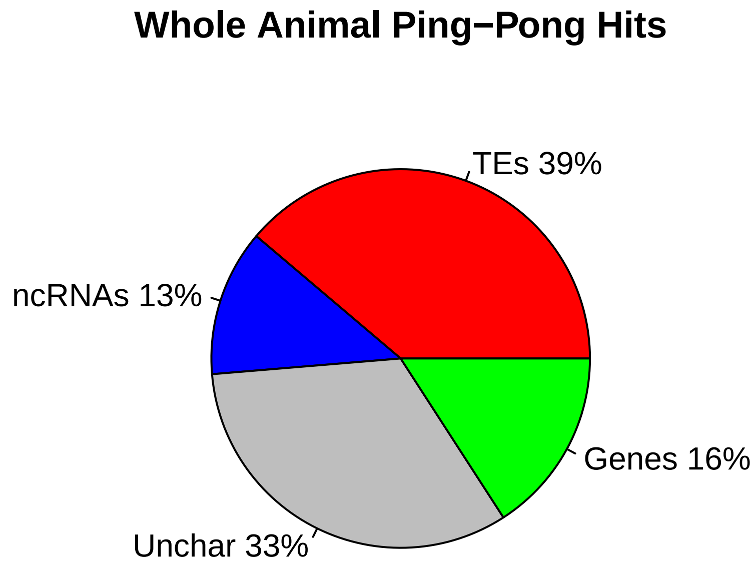


```
#Epithelial Animal
lbls <- c("TEs", "ncRNAs", "Unchar", "Genes")
colors = c("red", "blue", "gray", "green")
```

```r
pct <- round(Colch_ping_hits/sum(Colch_ping_hits)*100)
lbls <- paste(lbls, pct) # add percents to labels
lbls <- paste(lbls,"%",sep="") # ad % to labels
pie(Colch_ping_hits,labels = lbls, col=colors,
    main="Epithelial Animal Ping-Pong Hits")
```

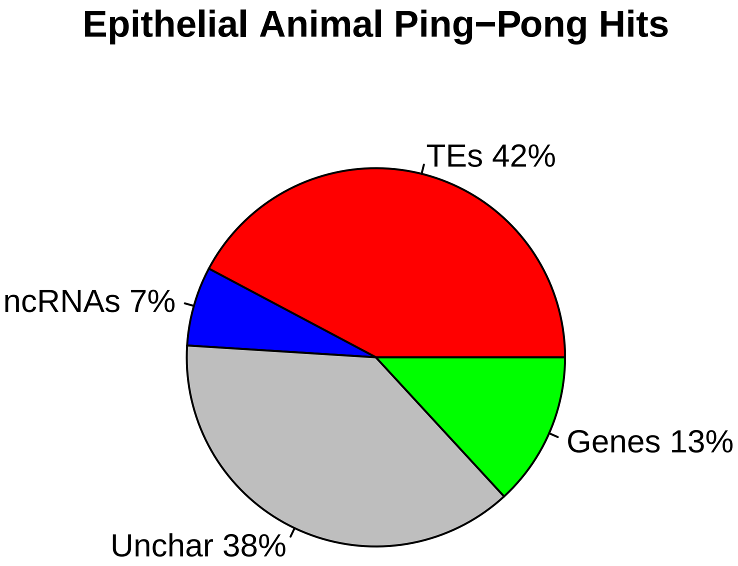


#### piRNA Positional Overlap Frequency Interrogation

Ping-pong processing of transcripts is known to occur in *Hydra* but whether this type of processing is active in somatic stem cells remains unknown. piRNAs from Whole Animals and Epithelial Animals were used to address this problem by generating piRNA overlap frequency plots. Ping-pong processing should consist of a preponderance of 10 bp overlaps between antisense- and sense-mapped piRNAs.

Positional information was extracted from piRNA BAM files and exported as a .txt file using the script: get_rsem_position.sh This script uses: "get_rep_piRNA_sense_BT.perl" and "get_rep_piRNA_antisense_BT.perl".

txt files were converted into BED files using the script: olap_bed_rsem_all.sh.

BED files containing piRNA overlap positions and overlap length between pairs of piRNAs (i.e. Hywi antisense/Hyli sense) were generated using the script windowbedall_rsem.sh.

The output of windowbedall_rsem.sh are BED files consisting of rows indexing a piRNA overlap event (ex. an overlap event between piRNA_1 and piRNA_2) within a 30 bp window.
The file contains has 11 columns:
1) Transcript on which piRNA_1 is mapped 2) Start position of piRNA_1 3) End position of piRNA_1 4) Sequence of piRNA_1 5) Copy number of piRNA_1

6) Transcript on which piRNA_2 is mapped 7) Start position of piRNA_2 8) End position of piRNA_2 9) Sequence of piRNA_2 10) Copy number of piRNA_2 11) piRNA overlap length

BED files are converted to txt files before importing into R.

```
#Load Matrices of piRNA overlap events

overlap_matrices <- list(read.table("objects/ping_pong/WT_HywiS_HywiAS_rsem.overlap.txt"),read.table("o

#Generate a function that provides the frequency of overlap events at a given distance between piRNA 5'

Generate_Freq <- function(x){
  x <- x[,c(5,10,11)]
  x.sub <- x[,1:2]
  x.min <- apply(x.sub, 1, min)
  x <- data.frame(frequency = x.min, Length = x[,3])
  x.table <- aggregate(frequency ~ Length, data = x, FUN = sum)
  x.table$Percentage <- (x.table$frequency/sum(x.table$frequency))*100
  x.table <- x.table[,c(1,3)]
  return(x.table)
}

Ping_Pong_List <- vector("list",length(overlap_matrices))

for (i in 1:length(overlap_matrices)){
  Ping_Pong_List[[i]] <- Generate_Freq(overlap_matrices[[i]])
  perc <- paste("Percentage_", i)
  colnames(Ping_Pong_List[[i]]) <- c("Length", perc)
}

Col_Names <- c("Length", "WT_HywiS_HywiAS","WT_HywiS_HyliAS", "WT_HyliS_HyliAS", "WT_HyliS_HywiAS", "Co

Ping_Pong_Overlap_Matrix <- Reduce(function(...) merge(..., by = "Length"), Ping_Pong_List)

colnames(Ping_Pong_Overlap_Matrix) <- paste(Col_Names, sep = "")

#Plot piRNA overlap frequency in whole animals

WT_Matrix <- Ping_Pong_Overlap_Matrix[,c(1:5)]

WTolapmelt <- melt(WT_Matrix, id.vars = "Length")

ggplot(data=WTolapmelt, aes(x=Length, y=value, group=variable, color= variable)) +
  geom_line()+
  geom_point()
```

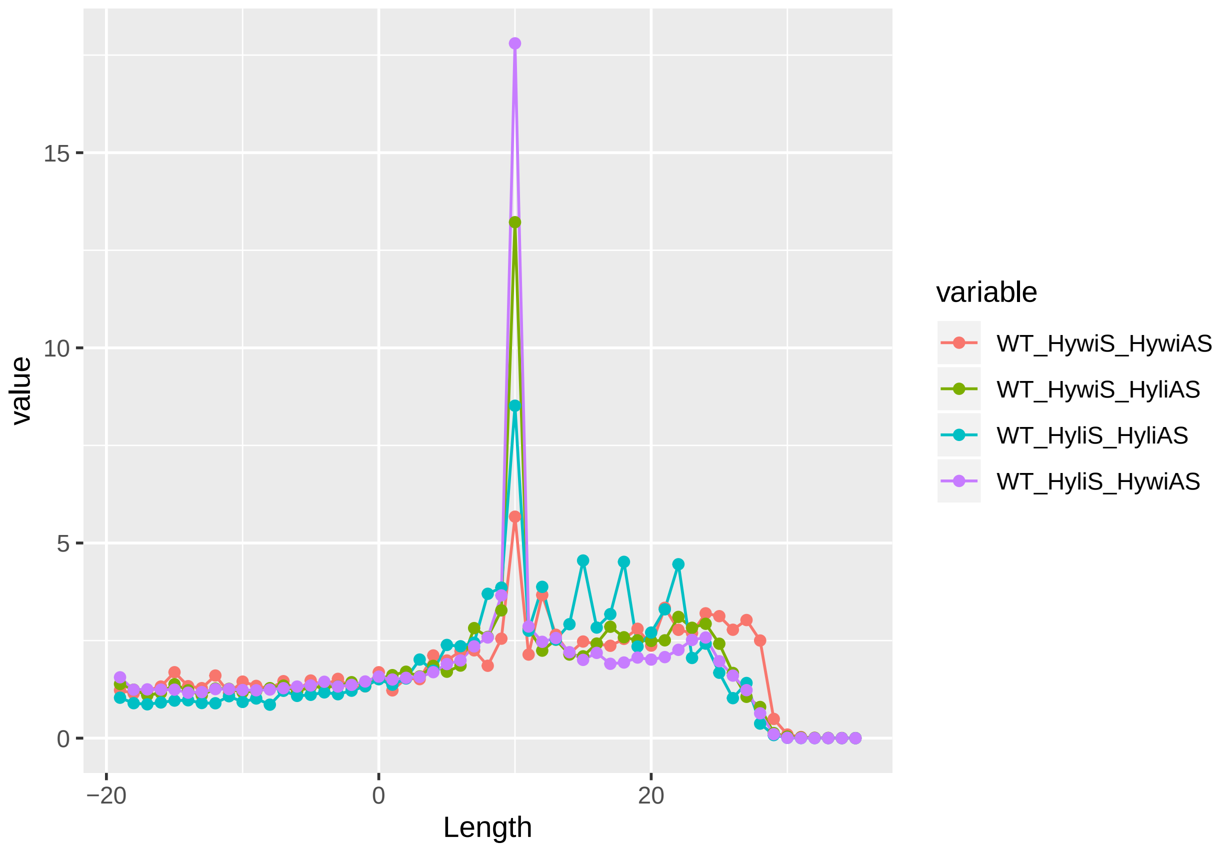


```
#Plot piRNA overlap frequency in epithelial animals

Colch_Matrix <- Ping_Pong_Overlap_Matrix[,c(1,6:9)]

Colcholapmelt <- melt(Colch_Matrix, id.vars = "Length")

ggplot(data=Colcholapmelt, aes(x=Length, y=value, group=variable, color= variable)) +
  geom_line()+
  geom_point()
```

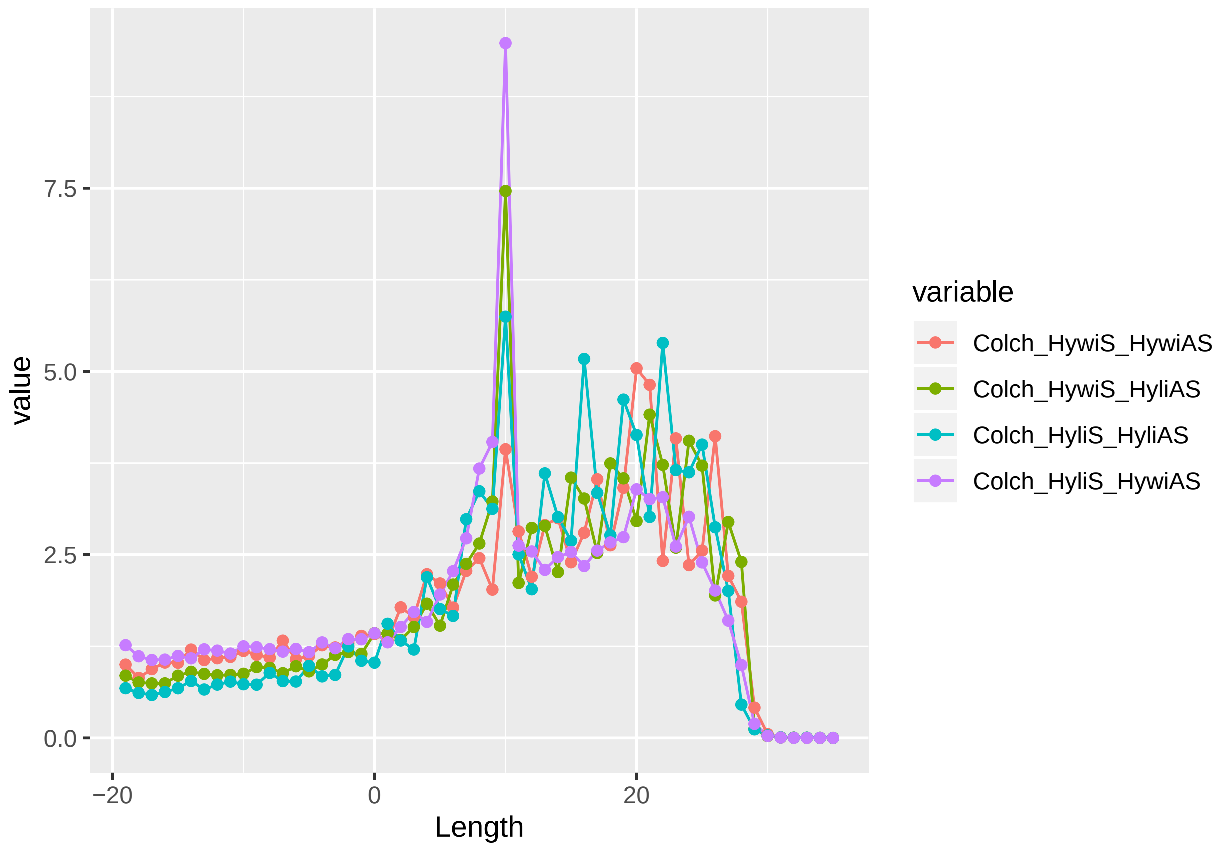


**Software versions**

This document was computed on Fri Aug 09 19:22:40 2019 with the following R package versions.

```
R version 3.5.3 (2019-03-11)
Platform: x86_64-apple-darwin15.6.0 (64-bit)
Running under: macOS Mojave 10.14.5

Matrix products: default
BLAS: /Library/Frameworks/R.framework/Versions/3.5/Resources/lib/libRblas.0.dylib
LAPACK: /Library/Frameworks/R.framework/Versions/3.5/Resources/lib/libRlapack.dylib

locale:
[1] en_US.UTF-8/en_US.UTF-8/en_US.UTF-8/C/en_US.UTF-8/en_US.UTF-8

attached base packages:
[1] stats     graphics  grDevices utils     datasets  methods   base

other attached packages:
[1] reshape2_1.4.3 ggplot2_3.2.0

loaded via a namespace (and not attached):
 [1] Rcpp_1.0.1       knitr_1.22       magrittr_1.5     tidyselect_0.2.5
 [5] munsell_0.5.0    colorspace_1.4-1 R6_2.4.0         rlang_0.4.0
```

```
 [9] plyr_1.8.4       stringr_1.4.0    dplyr_0.8.3      tools_3.5.3
[13] grid_3.5.3       gtable_0.3.0     xfun_0.5         withr_2.1.2
[17] htmltools_0.3.6  yaml_2.2.0       lazyeval_0.2.2   digest_0.6.20
[21] assertthat_0.2.1 tibble_2.1.3     crayon_1.3.4     purrr_0.3.2
[25] glue_1.3.1       evaluate_0.13    rmarkdown_1.12   labeling_0.3
[29] stringi_1.4.3    compiler_3.5.3   pillar_1.4.2     scales_1.0.0
[33] pkgconfig_2.0.2
```

### 4 Lineage-sorted piRNA Count Generation

*Bryan Teefy*

*08/09/2019*

#### Load Required Libraries

```r
library(dplyr)
library(reshape2)
library(ggplot2)
library(ggpubr)
library(VennDiagram)
```

#### Explore Lineage-sorted piRNA Diversity

To explore the diversity of piRNAs in different lineages, we identified the unique piRNAs in each lineage as well as the piRNAs species that were present in multiple lineages.

Trimmed piRNAs from Whole Animals were cross-referenced against lineage-sorted piRNA libraries (Juliano *et al.*, 2014) to retain lineage-specific piRNAs.

Lineage-specific piRNAs were saved in a R dataframe using the script, "Unique_piRNA_Generation.R" which uses the script, "run_lin_sorting.sh".

Unique and shared piRNAs were visualized using a Venn Diagram.

```r
# Load unique piRNA sequences sorted by lineage and protein origin

load("objects/Unique_Ecto_Hyli_piRNAs.Rda")

load("objects/Unique_Ecto_Hywi_piRNAs.Rda")

load("objects/Unique_Endo_Hyli_piRNAs.Rda")

load("objects/Unique_Endo_Hywi_piRNAs.Rda")

load("objects/Unique_Int_Hyli_piRNAs.Rda")

load("objects/Unique_Int_Hywi_piRNAs.Rda")

# Count the number of unique piRNAs per lineage

PIWI_Ecto <- merge(EctoHyli, EctoHywi, by = "seq", all = TRUE)
rm(EctoHyli, EctoHywi)

PIWI_Endo <- merge(EndoHyli, EndoHywi, by = "seq", all = TRUE)
rm(EndoHyli, EndoHywi)

PIWI_Int <- merge(IntHyli, IntHywi, by = "seq", all = TRUE)
rm(IntHyli, IntHywi)
```

```r
# Visualize shared and unique piRNA species by lineage

Venn <- list(Ectodermal_piRNAs = PIWI_Ecto$seq, Endodermal_piRNAs = PIWI_Endo$seq,
    Interstitial_piRNAs = PIWI_Int$seq)

venn.plot <- venn.diagram(Venn, filename = NULL, fill = c("blue", "red", "yellow"))
grid.draw(venn.plot)
```

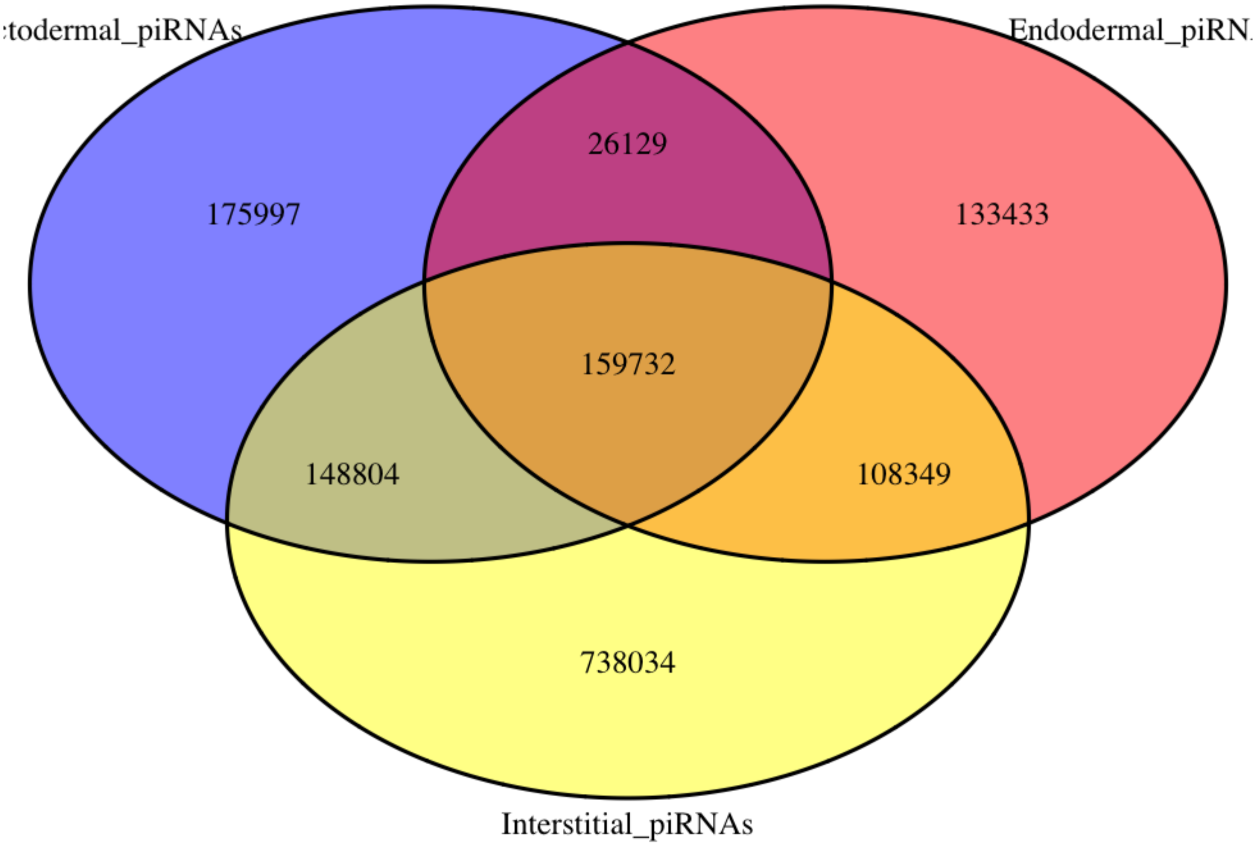


```r
rm(PIWI_Ecto, PIWI_Endo, PIWI_Int, Venn, venn.plot)
```

#### Generate Lineage-sorted piRNA Retaining Abundancies

To investigate piRNA targeting in each of the three cell lineages in *Hydra*, we cross-referenced lineage-specific small RNA libraries with the piRNA pulldown libraries from whole animals to generate lineage-specific piRNA libraries.

Trimmed piRNAs from Whole Animals were cross-referenced against lineage-sorted small RNA libraries (Juliano *et al.*, 2014) to retain only those piRNAs present in both libraries. Crucially, piRNA copy number from the piRNA pulldowns was maintained using the script, "small__RNA__piRNA__contrast.R" which uses the script, "run__contrast.sh".

The resultant lineage-sorted piRNA FASTA files were mapped to the *Hydra* transcriptome using RSEM functions rsem-calculate-expression and rsem-generate-data-matrix to generate a count matrix as in RMD 1.

Lineage-sorted piRNA mapping results are summarized in the table, "lin__rsem__piRNA__matrix.txt".

#### Load Transcriptome Annotation Matrix and Lineage-sorted piRNA Mapping Results

```
piRNA_Deg_counts <- read.table("objects/Annotated_piRNA_Degradome_Count_Matrix.txt",
    sep = "\t", check.names = FALSE, header = TRUE)

lin_matrix <- read.table("objects/lin_rsem_piRNA_matrix.txt", sep = "\t", header = T)
```

#### Generate Normalized piRNA Mapping Density Values

PIWI targets should have a high density of piRNA counts. We normalize piRNA counts by transcript length to determine piRNA count density.

To determine if the piRNA count density values were significantly different between classes of transcripts, we performed Tukey's Honest Significant Difference test to compare mean piRNA count density between each transcript type (i.e. TE, ncRNA, Unchar., Gene) for each piRNA class (i.e. Hywi Antisense-mapped, Hyli Sense-mapped, etc.).

```
# Merge lineage sorted piRNA counts with trancript length and transcript class
# data

lin_matrix <- merge(lin_matrix, piRNA_Deg_counts[, c(1, 3, 11)], by = "ID")

# To generate epithelial count values, take the mean of combined ectodermal and
# endodermal counts

lin_matrix$epi_Hyli_AS <- (lin_matrix$Ecto_Hyli_AS.isoforms.results + lin_matrix$Endo_Hyli_AS.isoforms.r

lin_matrix$epi_Hyli_S <- (lin_matrix$Ecto_Hyli_S.isoforms.results + lin_matrix$Endo_Hyli_S.isoforms.resu

lin_matrix$epi_Hywi_AS <- (lin_matrix$Ecto_Hywi_AS.isoforms.results + lin_matrix$Endo_Hywi_AS.isoforms.r

lin_matrix$epi_Hywi_S <- (lin_matrix$Ecto_Hywi_S.isoforms.results + lin_matrix$Endo_Hywi_S.isoforms.resu

# Calculate piRNA count density by dividing by counts by length in kilobases

norm <- (lin_matrix$Length/1000)

lin_matrix$epi_Hyli_AS_kb <- lin_matrix$epi_Hyli_AS/norm

lin_matrix$epi_Hyli_S_kb <- lin_matrix$epi_Hyli_S/norm

lin_matrix$epi_Hywi_AS_kb <- lin_matrix$epi_Hywi_AS/norm

lin_matrix$epi_Hywi_S_kb <- lin_matrix$epi_Hywi_S/norm

# Generate interstitial piRNA count density values

lin_matrix$int_Hyli_AS_kb <- lin_matrix$Int_Hyli_AS.isoforms.results/norm

lin_matrix$int_Hyli_S_kb <- lin_matrix$Int_Hyli_S.isoforms.results/norm
```

```r
lin_matrix$int_Hywi_AS_kb <- lin_matrix$Int_Hywi_AS.isoforms.results/norm

lin_matrix$int_Hywi_S_kb <- lin_matrix$Int_Hywi_S.isoforms.results/norm

# Perform Tukey's Honest Significant Difference test

# Group normalized mapping counts

Normalized_Mapping_Counts_Matrix <- lin_matrix[, c(20:27, 15)]

Feeder_Plots <- melt(Normalized_Mapping_Counts_Matrix, id.var = "Transcript_Class")

# Subset count density based on piRNA origin

epi_Hyli_AS_kb_Stats <- subset(Feeder_Plots, variable == "epi_Hyli_AS_kb")
epi_Hyli_S_kb_Stats <- subset(Feeder_Plots, variable == "epi_Hyli_S_kb")
epi_Hywi_AS_kb_Stats <- subset(Feeder_Plots, variable == "epi_Hywi_AS_kb")
epi_Hywi_S_kb_Stats <- subset(Feeder_Plots, variable == "epi_Hywi_S_kb")

int_Hyli_AS_kb_Stats <- subset(Feeder_Plots, variable == "int_Hyli_AS_kb")
int_Hyli_S_kb_Stats <- subset(Feeder_Plots, variable == "int_Hyli_S_kb")
int_Hywi_AS_kb_Stats <- subset(Feeder_Plots, variable == "int_Hywi_AS_kb")
int_Hywi_S_kb_Stats <- subset(Feeder_Plots, variable == "int_Hywi_S_kb")

# Develop Tukey Test Function

Tukey_Test <- function(x) {
    res.aov <- aov(value ~ Transcript_Class, data = x)
    return(TukeyHSD(res.aov))
}

# Run Tukey Test

Tukey_Test(epi_Hyli_AS_kb_Stats)
```

```
##   Tukey multiple comparisons of means
##     95% family-wise confidence level
##
## Fit: aov(formula = value ~ Transcript_Class, data = x)
##
## $Transcript_Class
##                     diff         lwr       upr     p adj
## ncRNA-Gene    142.23248   94.595365  189.8696 0.0000000
## TE-Gene       731.49377  663.236286  799.7512 0.0000000
## Unchar-Gene   199.60642  152.900244  246.3126 0.0000000
## TE-ncRNA      589.26129  514.859206  663.6634 0.0000000
## Unchar-ncRNA   57.37394    2.074276  112.6736 0.0385121
## Unchar-TE    -531.88734 -605.696846 -458.0778 0.0000000
```

```r
Tukey_Test(epi_Hyli_S_kb_Stats)
```

```
##   Tukey multiple comparisons of means
##     95% family-wise confidence level
##
```

```
## Fit: aov(formula = value ~ Transcript_Class, data = x)
##
## $Transcript_Class
##                     diff          lwr       upr     p adj
## ncRNA-Gene      7.227274   -6.3063246  20.76087 0.5170328
## TE-Gene       122.346413  102.9546143 141.73821 0.0000000
## Unchar-Gene    13.542257    0.2731351  26.81138 0.0433827
## TE-ncRNA      115.119140   93.9816730 136.25661 0.0000000
## Unchar-ncRNA    6.314984   -9.3955297  22.02550 0.7302595
## Unchar-TE    -108.804156 -129.7732718 -87.83504 0.0000000
```

```r
Tukey_Test(epi_Hywi_AS_kb_Stats)
```

```
##   Tukey multiple comparisons of means
##     95% family-wise confidence level
##
## Fit: aov(formula = value ~ Transcript_Class, data = x)
##
## $Transcript_Class
##                    diff        lwr       upr     p adj
## ncRNA-Gene    261.78351  147.84255  375.7245 0.0000000
## TE-Gene      1145.66905  982.40721 1308.9309 0.0000000
## Unchar-Gene   330.86298  219.14868  442.5773 0.0000000
## TE-ncRNA      883.88554  705.92672 1061.8444 0.0000000
## Unchar-ncRNA   69.07947  -63.18919  201.3481 0.5362372
## Unchar-TE    -814.80607 -991.34752 -638.2646 0.0000000
```

```r
Tukey_Test(epi_Hywi_S_kb_Stats)
```

```
##   Tukey multiple comparisons of means
##     95% family-wise confidence level
##
## Fit: aov(formula = value ~ Transcript_Class, data = x)
##
## $Transcript_Class
##                     diff        lwr        upr     p adj
## ncRNA-Gene     11.535023  -27.06414   50.13419 0.8690148
## TE-Gene       221.044393  165.73705  276.35173 0.0000000
## Unchar-Gene    21.036865  -16.80799   58.88172 0.4817158
## TE-ncRNA      209.509369  149.22321  269.79553 0.0000000
## Unchar-ncRNA    9.501842  -35.30610   54.30979 0.9479747
## Unchar-TE    -200.007528 -259.81353 -140.20152 0.0000000
```

```r
Tukey_Test(int_Hyli_AS_kb_Stats)
```

```
##   Tukey multiple comparisons of means
##     95% family-wise confidence level
##
## Fit: aov(formula = value ~ Transcript_Class, data = x)
##
## $Transcript_Class
##                    diff         lwr       upr     p adj
## ncRNA-Gene    232.96181  154.949002  310.9746 0.0000000
## TE-Gene      1196.48078 1084.699076 1308.2625 0.0000000
## Unchar-Gene   327.20555  250.717283  403.6938 0.0000000
## TE-ncRNA      963.51897  841.674577 1085.3634 0.0000000
```

```
## Unchar-ncRNA   94.24374    3.682369  184.8051 0.0376586
## Unchar-TE    -869.27523 -990.149185 -748.4013 0.0000000
```

```r
Tukey_Test(int_Hyli_S_kb_Stats)
```

```
##   Tukey multiple comparisons of means
##     95% family-wise confidence level
##
## Fit: aov(formula = value ~ Transcript_Class, data = x)
##
## $Transcript_Class
##                    diff          lwr        upr     p adj
## ncRNA-Gene     11.81402  -10.3373909   33.96543 0.5181634
## TE-Gene       199.98073  168.2407810  231.72068 0.0000000
## Unchar-Gene    22.15630    0.4377716   43.87482 0.0435122
## TE-ncRNA      188.16671  153.5695017  222.76392 0.0000000
## Unchar-ncRNA   10.34228  -15.3722483   36.05680 0.7298989
## Unchar-TE    -177.82443 -212.1460895 -143.50278 0.0000000
```

```r
Tukey_Test(int_Hywi_AS_kb_Stats)
```

```
##   Tukey multiple comparisons of means
##     95% family-wise confidence level
##
## Fit: aov(formula = value ~ Transcript_Class, data = x)
##
## $Transcript_Class
##                    diff         lwr       upr     p adj
## ncRNA-Gene     348.3751   216.46844  480.2817 0.0000000
## TE-Gene       1644.3352  1455.33100 1833.3393 0.0000000
## Unchar-Gene    464.4270   335.09811  593.7558 0.0000000
## TE-ncRNA      1295.9601  1089.94162 1501.9786 0.0000000
## Unchar-ncRNA   116.0519   -37.07222  269.1761 0.2084706
## Unchar-TE    -1179.9082 -1384.28585 -975.5305 0.0000000
```

```r
Tukey_Test(int_Hywi_S_kb_Stats)
```

```
##   Tukey multiple comparisons of means
##     95% family-wise confidence level
##
## Fit: aov(formula = value ~ Transcript_Class, data = x)
##
## $Transcript_Class
##                    diff         lwr        upr     p adj
## ncRNA-Gene     22.49505  -19.922810   64.91291 0.5230711
## TE-Gene       297.64593  236.866932  358.42494 0.0000000
## Unchar-Gene    34.23743   -7.351492   75.82635 0.1482836
## TE-ncRNA      275.15089  208.900501  341.40127 0.0000000
## Unchar-ncRNA   11.74238  -37.498503   60.98326 0.9281041
## Unchar-TE    -263.40850 -329.131232 -197.68578 0.0000000
```

### Visualizing piRNA Mapping

Since the range of observed count density values was large, we used a log scale to visualize piRNA count density. For boxplot visualization, we added a pseudocount to the raw piRNA counts to remove any 0 count density values that would return infinite values on a log scale. The pseudocount we chose was 0.01 since that was the lowest fractional count administered by our counting strategy. We explored piRNA count density for 1) Epithelial piRNAs and 2) Interstitial piRNAs.

```r
# Create pseudocount

pseudocount <- 0.01

# Add pseudocount to raw piRNA counts then generate piRNA count density values
# for epithelial piRNAs

boxplot_matrix <- lin_matrix[, c(2:15)]
boxplot_matrix[, c(1:12)] <- boxplot_matrix[, c(1:12)] + pseudocount

boxplot_matrix$epi_Hyli_AS <- (boxplot_matrix$Ecto_Hyli_AS.isoforms.results + boxplot_matrix$Endo_Hyli_

boxplot_matrix$epi_Hyli_S <- (boxplot_matrix$Ecto_Hyli_S.isoforms.results + boxplot_matrix$Endo_Hyli_S.

boxplot_matrix$epi_Hywi_AS <- (boxplot_matrix$Ecto_Hywi_AS.isoforms.results + boxplot_matrix$Endo_Hywi_

boxplot_matrix$epi_Hywi_S <- (boxplot_matrix$Ecto_Hywi_S.isoforms.results + boxplot_matrix$Endo_Hywi_S.

boxplot_matrix[, c(15:18)] <- boxplot_matrix[, c(15:18)]/(boxplot_matrix$Length/1000)

# Plot epithelial piRNA count density values

epi_matrix_feeder <- boxplot_matrix[, c(14, 15:18)]
epi_matrix_plotter <- melt(epi_matrix_feeder, id.var = "Transcript_Class")
colnames(epi_matrix_plotter) <- c("Transcript_Class", "piRNA_Origin", "piRNA_Mapping_Density")
epi_matrix_plotter$Transcript_Class <- factor(epi_matrix_plotter$Transcript_Class,
    levels = c("TE", "ncRNA", "Unchar", "Gene"))

epi_level_order <- c("epi_Hyli_AS", "epi_Hyli_S", "epi_Hywi_AS", "epi_Hywi_S")

epi_boxplot <- ggplot(data = epi_matrix_plotter, aes(x = factor(piRNA_Origin, level = epi_level_order),
    y = piRNA_Mapping_Density), log = "y") + geom_boxplot(aes(fill = Transcript_Class)) +
    scale_y_log10(breaks = scales::trans_breaks("log10", function(x) 10^x), labels = scales::trans_forma
        scales::math_format(10^.x)))

epi_boxplot + scale_fill_manual(values = c("red", "light blue", "grey", "green")) +
    theme(panel.grid.major = element_blank(), panel.grid.minor = element_blank(),
        panel.background = element_blank(), axis.line = element_line(colour = "black")) +
    theme(legend.text = element_text(size = rel(1))) + ggtitle("Eptihelial piRNA Mapping Density") +
    theme(plot.title = element_text(hjust = 0.5)) + xlab("piRNA Origin") + ylab("piRNA Mapping Density")
```

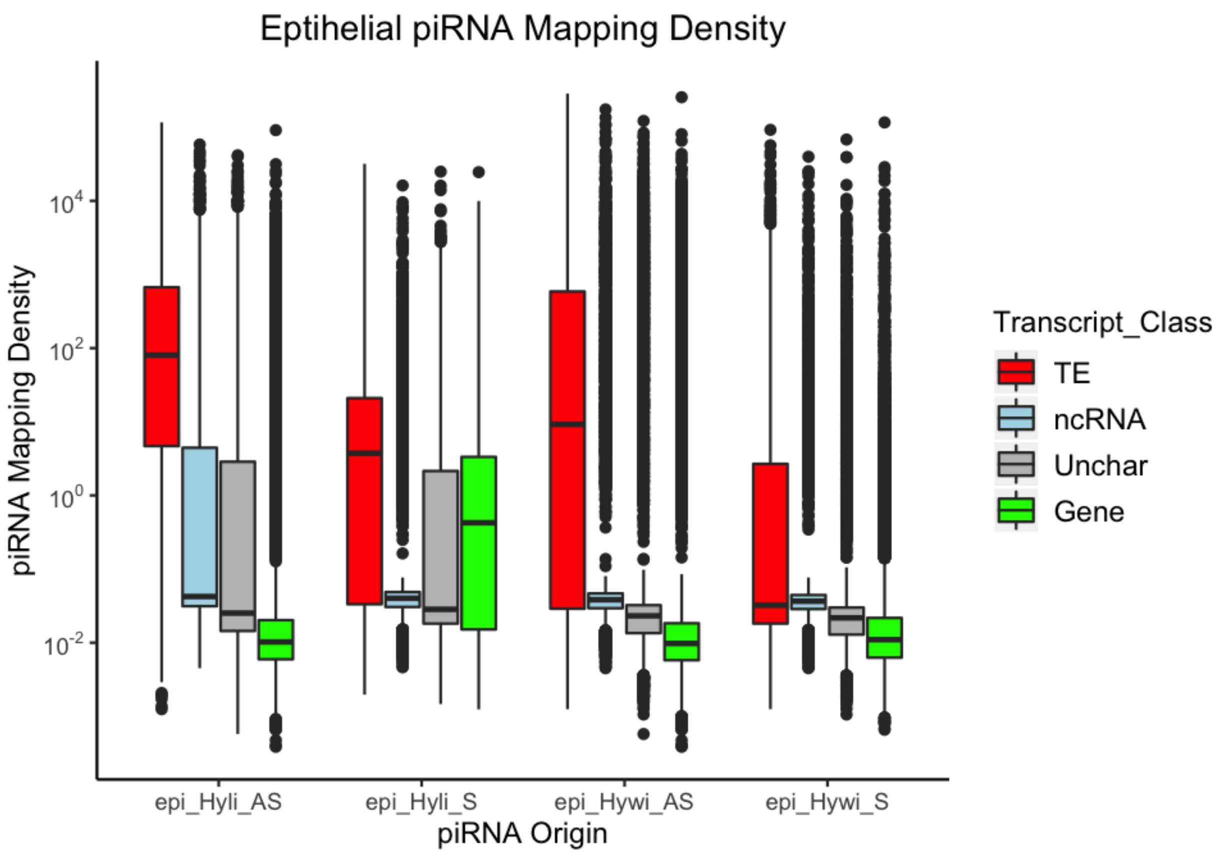


```
# Add pseudocount to raw piRNA counts then generate piRNA count density values
# for interstitial piRNAs

boplot_matrix$int_Hyli_AS <- (boxplot_matrix$Int_Hyli_AS.isoforms.results + boxplot_matrix$Int_Hyli_AS

boxplot_matrix$int_Hyli_S <- (boxplot_matrix$Int_Hyli_S.isoforms.results + boxplot_matrix$Int_Hyli_S.iso

boxplot_matrix$int_Hywi_AS <- (boxplot_matrix$Int_Hywi_AS.isoforms.results + boxplot_matrix$Int_Hywi_AS

boxplot_matrix$int_Hywi_S <- (boxplot_matrix$Int_Hywi_S.isoforms.results + boxplot_matrix$Int_Hywi_S.iso

boxplot_matrix[, c(19:22)] <- boxplot_matrix[, c(19:22)]/(boxplot_matrix$Length/1000)

# Plot interstitial piRNA count density values

int_matrix_feeder <- boxplot_matrix[, c(14, 19:22)]
int_matrix_plotter <- melt(int_matrix_feeder, id.var = "Transcript_Class")
colnames(int_matrix_plotter) <- c("Transcript_Class", "piRNA_Origin", "piRNA_Mapping_Density")
int_matrix_plotter$Transcript_Class <- factor(int_matrix_plotter$Transcript_Class,
    levels = c("TE", "ncRNA", "Unchar", "Gene"))

int_level_order <- c("int_Hyli_AS", "int_Hyli_S", "int_Hywi_AS", "int_Hywi_S")

int_boxplot <- ggplot(data = int_matrix_plotter, aes(x = factor(piRNA_Origin, level = int_level_order),
    y = piRNA_Mapping_Density), log = "y") + geom_boxplot(aes(fill = Transcript_Class)) +
    scale_y_log10(breaks = scales::trans_breaks("log10", function(x) 10^x), labels = scales::trans_forma
```

```
        scales::math_format(10^.x)))

int_boxplot + scale_fill_manual(values = c("red", "light blue", "grey", "green")) +
    theme(panel.grid.major = element_blank(), panel.grid.minor = element_blank(),
        panel.background = element_blank(), axis.line = element_line(colour = "black")) +
    theme(legend.text = element_text(size = rel(1))) + ggtitle("Interstitial Mapping Density") +
    theme(plot.title = element_text(hjust = 0.5)) + xlab("piRNA Origin") + ylab("piRNA Mapping Density")
```

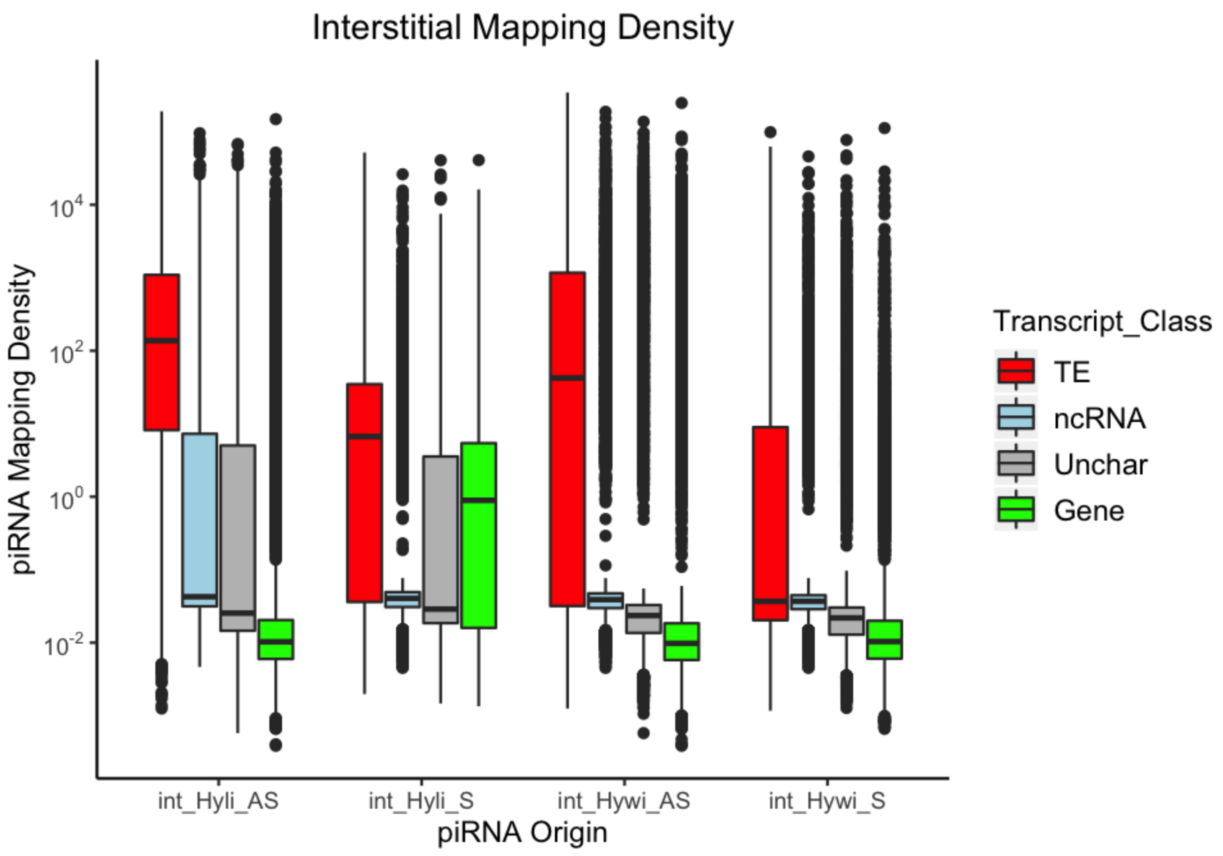


**Software versions**

This document was computed on Fri Aug 09 19:23:44 2019 with the following R package versions.

```
R version 3.5.3 (2019-03-11)
Platform: x86_64-apple-darwin15.6.0 (64-bit)
Running under: macOS Mojave 10.14.5

Matrix products: default
BLAS: /Library/Frameworks/R.framework/Versions/3.5/Resources/lib/libRblas.0.dylib
LAPACK: /Library/Frameworks/R.framework/Versions/3.5/Resources/lib/libRlapack.dylib

locale:
[1] en_US.UTF-8/en_US.UTF-8/en_US.UTF-8/C/en_US.UTF-8/en_US.UTF-8

attached base packages:
```

```
[1] grid      stats     graphics  grDevices utils     datasets  methods
[8] base

other attached packages:
[1] VennDiagram_1.6.20  futile.logger_1.4.3 ggpubr_0.2
[4] magrittr_1.5        ggplot2_3.2.0       reshape2_1.4.3
[7] dplyr_0.8.3         knitr_1.22

loaded via a namespace (and not attached):
 [1] Rcpp_1.0.1           munsell_0.5.0        tidyselect_0.2.5
 [4] colorspace_1.4-1     R6_2.4.0             rlang_0.4.0
 [7] stringr_1.4.0        plyr_1.8.4           tools_3.5.3
[10] gtable_0.3.0         xfun_0.5             lambda.r_1.2.3
[13] withr_2.1.2          htmltools_0.3.6      lazyeval_0.2.2
[16] yaml_2.2.0           assertthat_0.2.1     digest_0.6.20
[19] tibble_2.1.3         crayon_1.3.4         purrr_0.3.2
[22] formatR_1.7          futile.options_1.0.1 glue_1.3.1
[25] evaluate_0.13        rmarkdown_1.12       stringi_1.4.3
[28] compiler_3.5.3       pillar_1.4.2         scales_1.0.0
[31] pkgconfig_2.0.2
```

### 5 Single Cell Data Exploration

*Bryan Teefy*

*08/09/2019*

#### Load required libraries

```
library(URD)
library(Seurat)
```

#### Single Cell Data Exploration

We analyzed the homeostatic expression patterns of the 36 high mapping gene transcripts upregulated in response to hywi knockdown by interrogating single-cell expression data (Siebert, 2019). To interrogate expression, we used URD spline objects for ectoderm and endoderm. These objects contain expression data for genes that are expressed in at least 1% of the ectodermal or endodermal epithelial cells. We found epithelial expression for 24 of the 36 putative gene targets: 1) nineteen gene transcripts are expressed in both the endodermal and ectodermal epithelial lineages, 2) one transcript is expressed only in the ectodermal epithelial lineage, and 3) four transcripts are expressed only in the endodermal epithelial lineage. No epithelial expression was found for 12 of the gene transcripts, which could indicate low expression in a homeostatic animal. Higher expression at the extremities in both the ectoderm and endoderm and lower expression in body regions would be consistent with Hywi-mediated repression. Putative targets could show high expression. The gene, t14391, shows such a pattern.

#### Load the Necessary Files

```
# Load the list of searchable high mapping genes for each lineage

endo_mappers <- read.table("objects/high_mappers_endo.txt", header = T)

ecto_mappers <- read.table("objects/high_mappers_ecto.txt", header = T)

# Load the seurat object to load the searchable transcript IDs The seurat object
# (file Hydra_Seurat_Whole_Transcriptome) can be downloaded from Dryad:
# https://doi.org/10.5061/dryad.v5r6077. After download, place in the 'objects'
# folder

Hydra_Object <- readRDS("objects/Hydra_Seurat_Whole_Transcriptome.rds")

# Load single cell pseudotime objects (spline objects) and High Mapping Gene List
# The spline objects (file Hydra_URD_analysis_objects) can be downloaded from
# Dryad: https://doi.org/10.5061/dryad.v5r6077. After download, place in the
# 'objects' folder

endoderm.splines <- readRDS("objects/Splines-Endoderm.rds")
ectoderm.splines <- readRDS("objects/Splines-Ectoderm.rds")
```

#### Data Exploration

```r
# Establish hFind function to return the searchable transcript IDs

hFind <- function(x) {
    return(Hydra_Object@data@Dimnames[[1]][grep(x, Hydra_Object@data@Dimnames[[1]],
        ignore.case = T)])
}

# Run hFind function to convert to searchable transcript IDs

endo_mappers$ID <- sapply(endo_mappers$ID, FUN = hFind)

ecto_mappers$ID <- sapply(ecto_mappers$ID, FUN = hFind)
```

```r
# Generate Endodermal Plots

plotSmoothFitMultiCascade(endoderm.splines, genes = endo_mappers$ID[1:12], scaled = F,
    colors = c(`Foot/Body` = "#FF8C00", Tentacle = "#1E90FF", Hypostome = "#32CD32"),
    ncol = 1)
```

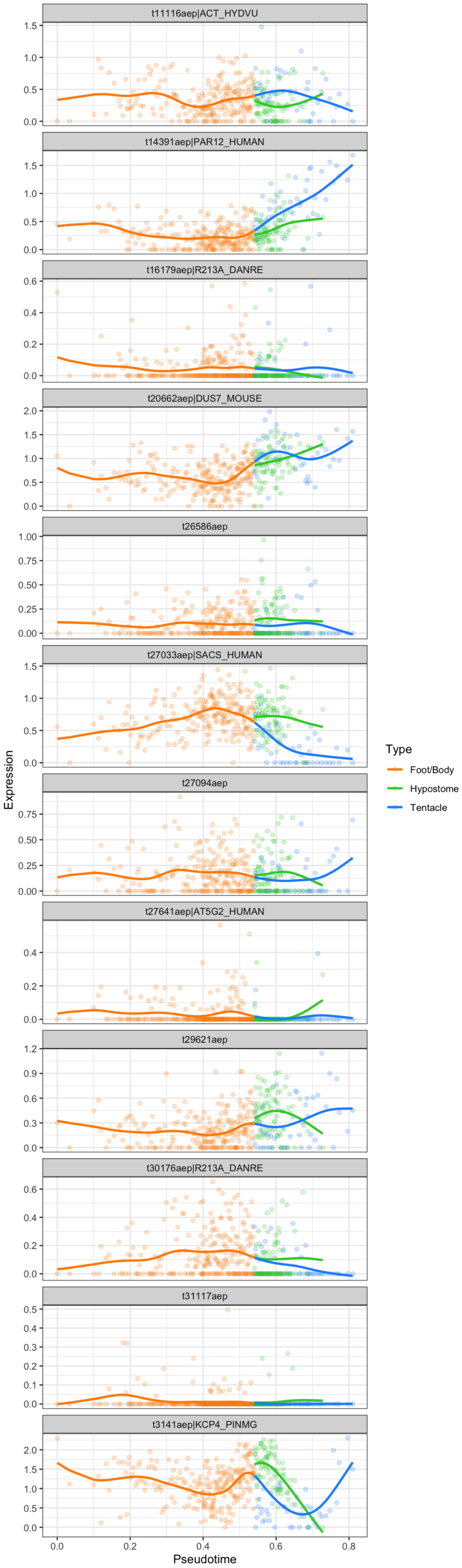
t11116aep|ACT_HYDVU
t14391aep|PAR12_HUMAN
t16179aep|R213A_DANRE
t20662aep|DUS7_MOUSE
t26586aep
t27033aep|SACS_HUMAN
t27094aep
t27641aep|AT5G2_HUMAN
t29621aep
t30176aep|R213A_DANRE
t31117aep
t3141aep|KCP4_PINMG
Expression
Pseudotime
Type
Foot/Body
Hypostome
Tentacle

```r
plotSmoothFitMultiCascade(endoderm.splines, genes = endo_mappers$ID[13:23], scaled = F,
    colors = c(`Foot/Body` = "#FF8C00", Tentacle = "#1E90FF", Hypostome = "#32CD32"),
    ncol = 1)
```

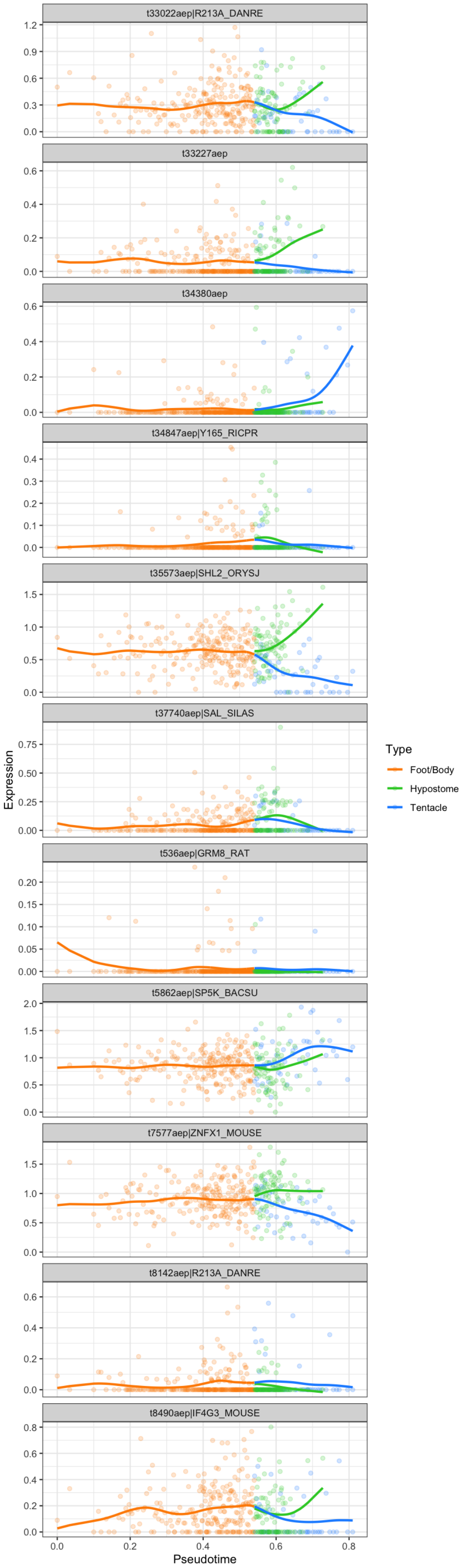
t33022aep|R213A_DANRE
t33227aep
t34380aep
t34847aep|Y165_RICPR
t35573aep|SHL2_ORYSJ
t37740aep|SAL_SILAS
t536aep|GRM8_RAT
t5862aep|SP5K_BACSU
t7577aep|ZNFX1_MOUSE
t8142aep|R213A_DANRE
t8490aep|IF4G3_MOUSE
Expression
Pseudotime
Type
Foot/Body
Hypostome
Tentacle

```r
# Generate Ectodermal Plots

plotSmoothFitMultiCascade(ectoderm.splines, genes = ecto_mappers$ID[1:9], scaled = F,
    colors = c(`Basal Disc` = "#FF8C00", `Body Column` = "#000000", Tentacle = "#1E90FF",
        Hypostome = "#32CD32"), ncol = 1)
```

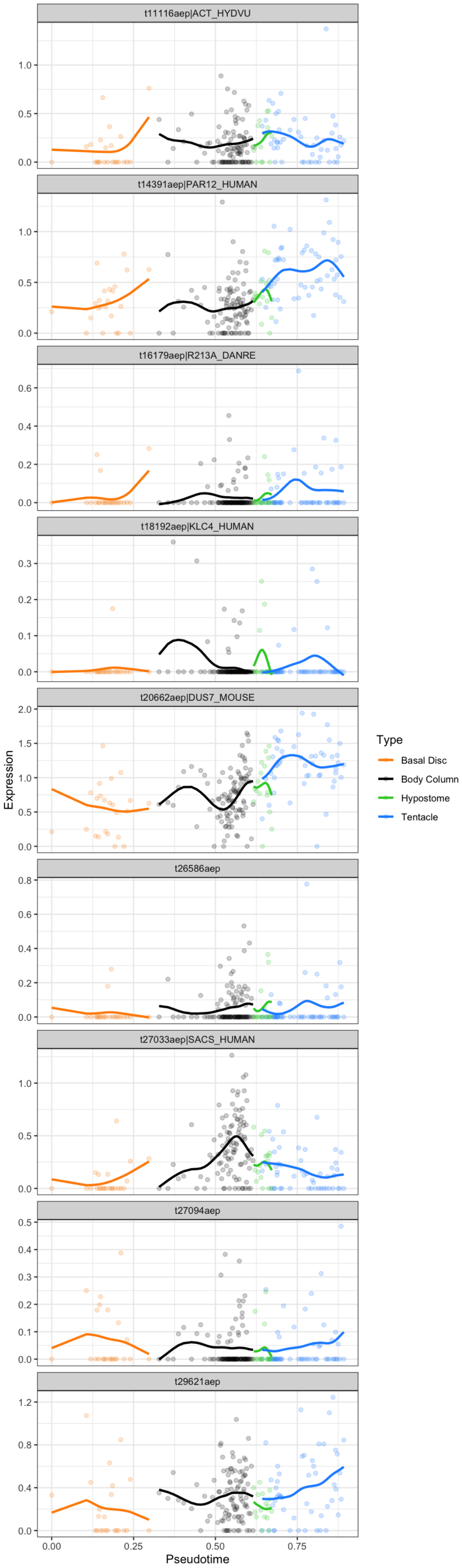

t11116aep|ACT_HYDVU
t14391aep|PAR12_HUMAN
t16179aep|R213A_DANRE
t18192aep|KLC4_HUMAN
t20662aep|DUS7_MOUSE
t26586aep
t27033aep|SACS_HUMAN
t27094aep
t29621aep
Expression
Pseudotime
Type
Basal Disc
Body Column
Hypostome
Tentacle

```r
plotSmoothFitMultiCascade(ectoderm.splines, genes = ecto_mappers$ID[10:20], scaled = F,
    colors = c(`Basal Disc` = "#FF8C00", `Body Column` = "#000000", Tentacle = "#1E90FF",
        Hypostome = "#32CD32"), ncol = 1)
```

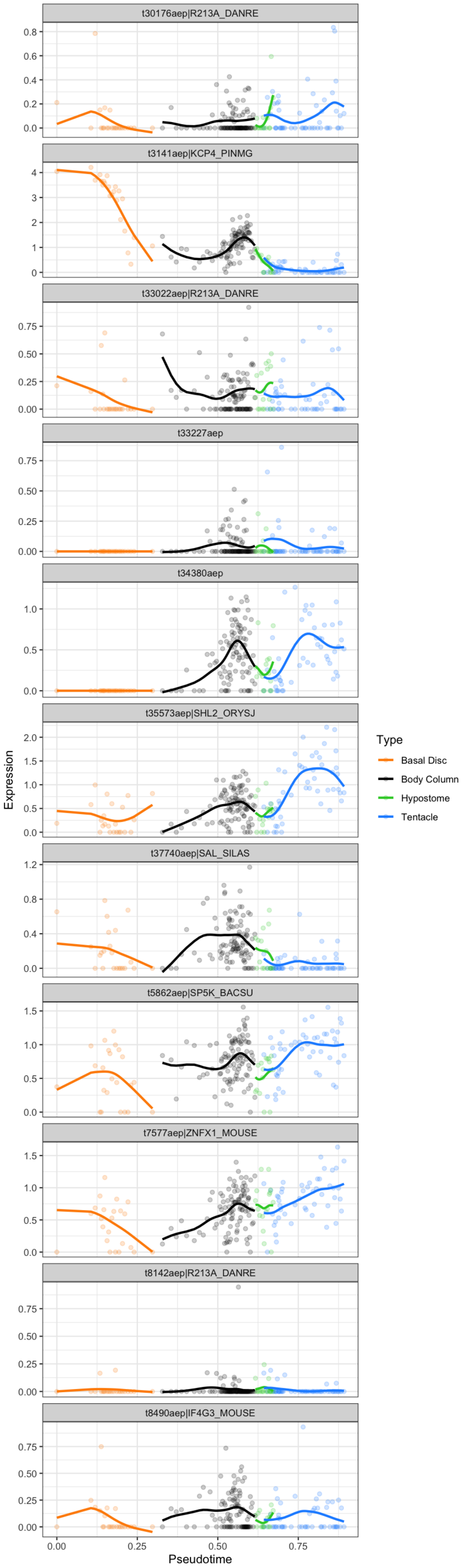
t30176aep|R213A_DANRE
t3141aep|KCP4_PINMG
t33022aep|R213A_DANRE
t33227aep
t34380aep
t35573aep|SHL2_ORYSJ
t37740aep|SAL_SILAS
t5862aep|SP5K_BACSU
t7577aep|ZNFX1_MOUSE
t8142aep|R213A_DANRE
t8490aep|IF4G3_MOUSE
Expression
Pseudotime
Type
Basal Disc
Body Column
Hypostome
Tentacle

**Software versions**

This document was computed on Fri Aug 09 19:25:35 2019 with the following R package versions.

```
R version 3.5.3 (2019-03-11)
Platform: x86_64-apple-darwin15.6.0 (64-bit)
Running under: macOS Mojave 10.14.5

Matrix products: default
BLAS: /Library/Frameworks/R.framework/Versions/3.5/Resources/lib/libRblas.0.dylib
LAPACK: /Library/Frameworks/R.framework/Versions/3.5/Resources/lib/libRlapack.dylib

locale:
[1] en_US.UTF-8/en_US.UTF-8/en_US.UTF-8/C/en_US.UTF-8/en_US.UTF-8

attached base packages:
[1] stats     graphics  grDevices utils     datasets  methods   base

other attached packages:
[1] Seurat_2.3.4  cowplot_0.9.4 URD_1.0.3     Matrix_1.2-16 ggplot2_3.2.0
[6] knitr_1.22

loaded via a namespace (and not attached):
  [1] reticulate_1.11.1          R.utils_2.8.0
  [3] tidyselect_0.2.5           htmlwidgets_1.3
  [5] grid_3.5.3                 trimcluster_0.1-2.1
  [7] ranger_0.11.2              BiocParallel_1.14.2
  [9] Rtsne_0.15                 munsell_0.5.0
 [11] destiny_2.10.2             codetools_0.2-16
 [13] ica_1.0-2                  withr_2.1.2
 [15] colorspace_1.4-1           Biobase_2.40.0
 [17] rstudioapi_0.9.0           stats4_3.5.3
 [19] ROCR_1.0-7                 robustbase_0.93-3
 [21] dtw_1.20-1                 vcd_1.4-4
 [23] VIM_4.8.0                  TTR_0.23-4
 [25] gbRd_0.4-11                labeling_0.3
 [27] Rdpack_0.10-1              lars_1.2
 [29] GenomeInfoDbData_1.1.0     polyclip_1.10-0
 [31] bit64_0.9-7                farver_1.1.0
 [33] xfun_0.5                   ggthemes_4.2.0
 [35] diptest_0.75-7             R6_2.4.0
 [37] GenomeInfoDb_1.16.0        RcppEigen_0.3.3.5.0
 [39] hdf5r_1.0.1                flexmix_2.3-15
 [41] bitops_1.0-6               DelayedArray_0.6.6
 [43] assertthat_0.2.1           SDMTools_1.1-221
 [45] scales_1.0.0               ggraph_1.0.2
 [47] nnet_7.3-12                gtable_0.3.0
 [49] npsurv_0.4-0               rlang_0.4.0
 [51] scatterplot3d_0.3-41       splines_3.5.3
 [53] lazyeval_0.2.2             acepack_1.4.1
 [55] checkmate_1.9.1            yaml_2.2.0
 [57] reshape2_1.4.3             abind_1.4-5
 [59] backports_1.1.4            Hmisc_4.2-0
```

```
 [61] tools_3.5.3                 gplots_3.0.1.1
 [63] RColorBrewer_1.1-2          proxy_0.4-23
 [65] BiocGenerics_0.26.0         ggridges_0.5.1
 [67] Rcpp_1.0.1                  plyr_1.8.4
 [69] base64enc_0.1-3             zlibbioc_1.26.0
 [71] purrr_0.3.2                 RCurl_1.95-4.12
 [73] rpart_4.1-13                pbapply_1.4-0
 [75] viridis_0.5.1               S4Vectors_0.18.3
 [77] zoo_1.8-4                   SummarizedExperiment_1.10.1
 [79] haven_2.1.0                 ggrepel_0.8.1
 [81] cluster_2.0.7-1             magrittr_1.5
 [83] data.table_1.12.2           openxlsx_4.1.0.1
 [85] gmodels_2.18.1              lmtest_0.9-36
 [87] RANN_2.6.1                  mvtnorm_1.0-10
 [89] fitdistrplus_1.0-14         matrixStats_0.54.0
 [91] hms_0.4.2                   lsei_1.2-0
 [93] evaluate_0.13               smoother_1.1
 [95] rio_0.5.16                  mclust_5.4.2
 [97] readxl_1.3.1                IRanges_2.14.12
 [99] gridExtra_2.3               compiler_3.5.3
[101] tibble_2.1.3                KernSmooth_2.23-15
[103] crayon_1.3.4                R.oo_1.22.0
[105] htmltools_0.3.6             segmented_0.5-3.0
[107] Formula_1.2-3               snow_0.4-3
[109] tidyr_0.8.3                 tweenr_1.0.1
[111] formatR_1.7                 MASS_7.3-51.1
[113] fpc_2.1-11.1                boot_1.3-20
[115] car_3.0-3                   R.methodsS3_1.7.1
[117] gdata_2.18.0                parallel_3.5.3
[119] metap_1.1                   igraph_1.2.4.1
[121] GenomicRanges_1.32.7        forcats_0.4.0
[123] pkgconfig_2.0.2             foreign_0.8-71
[125] laeken_0.5.0                sp_1.3-1
[127] foreach_1.4.4               XVector_0.20.0
[129] minpack.lm_1.2-1            bibtex_0.4.2
[131] stringr_1.4.0               digest_0.6.20
[133] tsne_0.1-3                  rmarkdown_1.12
[135] cellranger_1.1.0            htmlTable_1.13.1
[137] curl_3.3                    kernlab_0.9-27
[139] gtools_3.8.1                modeltools_0.2-22
[141] jsonlite_1.6                nlme_3.1-137
[143] carData_3.0-2               viridisLite_0.3.0
[145] pillar_1.4.2                lattice_0.20-38
[147] httr_1.4.0                  DEoptimR_1.0-8
[149] survival_2.43-3             glue_1.3.1
[151] xts_0.11-2                  zip_2.0.3
[153] png_0.1-7                   prabclus_2.2-7
[155] iterators_1.0.10            bit_1.1-14
[157] ggforce_0.2.2               class_7.3-15
[159] stringi_1.4.3               mixtools_1.1.0
[161] doSNOW_1.0.16               latticeExtra_0.6-28
[163] caTools_1.17.1.2            dplyr_0.8.3
[165] irlba_2.3.3                 e1071_1.7-2
[167] ape_5.2
```
